# To prosper, live long: understanding the sources of reproductive skew and extreme reproductive success in structured populations

**DOI:** 10.1101/2024.03.07.583934

**Authors:** Robin E. Snyder, Stephen P. Ellner

## Abstract

In many species, a few individuals produce most of the next generation. How much of this reproductive skew is driven by variation among individuals in fixed traits, how much by external factors, and how much by random chance? And what does it take to have truly exceptional lifetime reproductive output (LRO)? In the past, we and others have partitioned the variance of LRO as a proxy for reproductive skew. Here we explain how to partition LRO skewness itself into contributions from fixed trait variation; four forms of “demographic luck” (birth state, fecundity luck, survival trajectory luck, and growth trajectory luck); and two kinds of “environmental luck” (birth environment, and environment trajectory). Each of these is further partitioned into contributions at different ages. We also determine what we can infer about individuals with exceptional LRO. We find that reproductive skew is largely driven by random variation in lifespan, and exceptional LRO generally results from exceptional lifespan. Other kinds of luck frequently bring skewness down, rather than increasing it. In populations where fecundity varies greatly with environmental conditions, getting a good year at the right time can be an alternate route to exceptional LRO, so that LRO is less predictive of lifespan.

## Introduction

Even in the absence of social dominance, the distribution of lifetime reproductive output (LRO) is often highly skewed, with a few individuals producing most of the offspring (e.g. Eusemann and Liesebach (2021); Gerzabek et al. (2017); Goodwin et al. (2016); Le Boeuf et al. (2019)). Reproductive skew can have substantial evolutionary consequences. If the reasons for large reproductive inequality are heritable there is a large opportunity for selection. If not, then highly skewed genotype-independent variation in realized fitness can change the consequences of genetic drift for outcomes such as the probability of allele fixation and the mean time to fixation (Eldon and Stephan, 2023; Tuljapurkar and Zuo, 2023). Large reproductive skew also has potential management implications: if we can predict which individuals are more likely to dominate reproduction, these are the ones we should target in an intervention, whether we aim to preserve a population or to extirpate it. We also just marvel at exceptional individuals and wonder how they came to be that way.^1^ Some researchers have looked for variation in strategy or quality that allows the most successful individuals to dominate reproduction (e.g. Annett and Pierotti (1999); Péron (2023)); however, our past work suggests that exceptional success may be mostly random, resulting from some combination of rapid early growth or maturation, a favorable environment at the right time, unusually large clutch sizes, and a long life (Snyder and Ellner, 2018, 2022; Snyder et al., 2021). Even then, we can ask exactly how an individual needs to be lucky to end up far out on the reproductive tail, since not all of these forms of luck may be equally important for having exceptional LRO.

In past work we have shown how to partition the variance of LRO implied by a density-independent structured population model into contributions from trait variation, luck in survival, luck in growth, luck in favorable/unfavorable environments, and luck in fecundity at different ages (Snyder and Ellner, 2022; Snyder et al., 2021). We, like others, found that trait variation (i.e. variation in any unchanging life-long attribute, such as genotype, phenotype, birth weight, the location of a sessile organism) always contributes less — usually much less — to the variance in LRO than the contribution of random chance (Broekman et al., 2020; Hartemink and Caswell, 2018; Jenouvrier et al., 2018; Snyder and Ellner, 2018; Steiner and Tuljapurkar, 2012).^2^ Luck dominates because of the many possible trajectories through life: even with identical traits and a common environment, different individuals will typically experience a different series of size or stage transitions and die at different ages, leading to variation in lifetime reproduction. As a result, the highly successful are not necessarily exceptional in any way other than their reproductive success (Chen et al., 2019; Liu et al., 2019). They’re just lucky.

The variance of LRO measures the breadth of possible outcomes, but to understand why the distribution of LRO is so lopsided, we need to similarly partition the skewness of LRO into contributions at different ages from different kinds of luck. In this paper, we show how to do that, and apply the new methods to a set of case studies with contrasting life histories. We ask what produces the tail in LRO. To what degree is reproductive skew driven by trait differences and to what degree by luck? To the extent that skew is driven by luck, is it luck in survival, in growth, in environment, in fecundity? And we ask what it takes to end up in the far right tail of the distribution. If the right question isn’t so much “Why is this individual special?” as “How did this individual get so lucky?”, we can still ask what kind of luck it takes to be especially successful.

It is important to note that we are analyzing the lives of individuals, not the output of a cohort. In populations where temporal variation in environmental conditions affects demographic rates, we can interpret these results as representing the distribution of outcomes across a cohort of individuals only if each individual in the cohort experiences their own independent sequence of environment states. This may be nearly true if environmental variation is spatiotemporal with a fine spatial grain — patchy fires or local light environments in a forest may be good examples (Coutts et al., 2021; Metcalf et al., 2009).

As with reproductive variance, we find in empirical examples that reproductive skew results mostly from luck, rather than from trait differences. Unlike LRO variance, which can be produced by various forms of luck depending on life history, reproductive skew is generated mostly by differences in lifespan. Relatedly, we find that individuals with exceptional LRO generally have exceptionally long lives, although the degree to which LRO constrains lifespan varies: when fecundity varies substantially with environment conditions, getting a good year can partially substitute for a long life. If fecundity varies wildly with environment, the typical clutch size in an exceptionally good year may be enough to guarantee exceptional LRO. In that case, being in the right environment at the right time can provide another route into the right-hand tail of the LRO distribution.

## Methods

### Overview

Our approach to partitioning skewness in lifetime outcomes into contributions from different kinds of luck at different ages is conceptually very similar to our approach for partitioning variance (Snyder and Ellner, 2022; Snyder et al., 2021). We are deriving information about the role of luck from a demographic model which has been parameterized from data. The underlying model is a discrete-time density-independent matrix model, integral projection model, or agent-based model incorporating population structure. While usually used to project entire populations over time, these models are built from individual state-fate relationship models that describe what happens to an individual over the course of one year (or one time step of the model), as a function of their current state (e.g. size or stage). What are their odds of survival? What is their expected fecundity this year? If they live, what is the probability distribution for their state or size at the next census?

For our analysis, each time step in the life of an individual is conceptually divided up into a series of sub-steps updating first reproductive output, then survival, then growth to a new size/stage, and finally the state of the environment, if the environment is time-varying, as shown in Fig. 1. As noted in the Introduction, if we interpret our analysis as calculating the distribution of realized outcomes across a cohort of many individuals (rather than the probability distribution of possible outcomes for one individual), we must assume that each individual experiences an environmental sequence that is independent of its neighbors: we do not yet know how to account for environments that are correlated across individuals. We then re-write the original model, which moves directly from age *a* to age *a*+1, as a model where these transitions in fecundity, survival, etc. occur separately and sequentially between ages *a* and *a* +1. This requires expanding the state space, so that individual state also includes the information: which of those transitions have you most recently experienced? The expanded model’s time step is then a portion of a year. For that expanded model, results of van Daalen and Caswell (2017) let us compute the first, second, and third moments of future LRO and lifespan given the individual’s current state, and thus the moments, variance, and skewness conditional on the individual’s history up to that moment. Learning what actually happened during one more unpredictable transition (e.g. learning that the individual lived from age 5 to age 6) changes the conditional moments. “Learning what actually happened “ is equivalent to “removing the luck”: chance is replaced by certainty. Thus, the change in variance or skewness when we learn what actually happened measures how much the luck in that transition contributed to lifetime total variance or skewness. Fecundity luck is ascertained by learning how many offspring an individual produced at that age; survival trajectory luck by knowing whether an individual survived or died at that age; growth trajectory luck by knowing what size/stage the individual transitioned to; and environment trajectory luck by knowing the next state of the environment. To get one characteristic value for the population of the contribution of each kind of luck at each age, rather than a different value for each individual, we average across individuals based on the age-dependent state distribution, including those individuals that are already dead.

**Figure 1:**
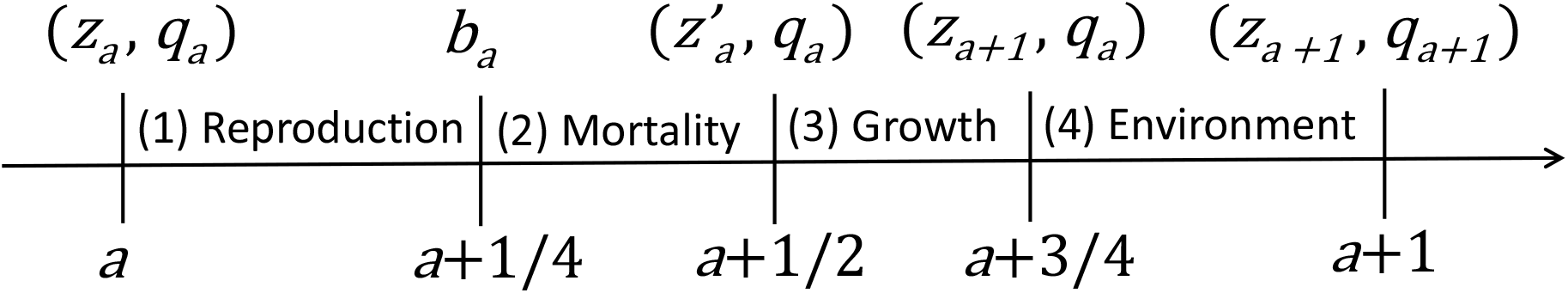
The assumed sequence of events for partitioning luck by transition type and age. At age *a* each individual has individual state *z*_*a*_ and possibly environment state *q*_*a*_. Within the time step carrying an individual from age *a* to age *a*+1 the order of events is (1) reproduction between *a* and *a*+1/4, (2) mortality between *a*+1/4 and *a*+1/2, (3) growth (or state transition) between *a*+1/2 and *a*+3/4, and (4) update of environment state between *a*+3/4 and *a*+1, in models with environmental variability. (1) *z*_*a*_ and *q*_*a*_ determine the offspring production *B*_*a*_ at age *a*, from which a realized value *b*_*a*_ is drawn at age *a*+1/4, representing some arbitrary point between ages *a* and *a*+1. (2) After reproduction, each individual either survives 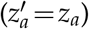 or dies 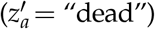, (3) All surviving individuals then transition to their subsequent state *z*_*a*+1_. (4) In models with Markovian environmental variation, all transition probabilities between ages *a* and *a*+1 are affected by the environment state *q*_*a*_. The final event within each time step is a random transition to the next environment state *q*_*a*+1_.

**Figure 2:**
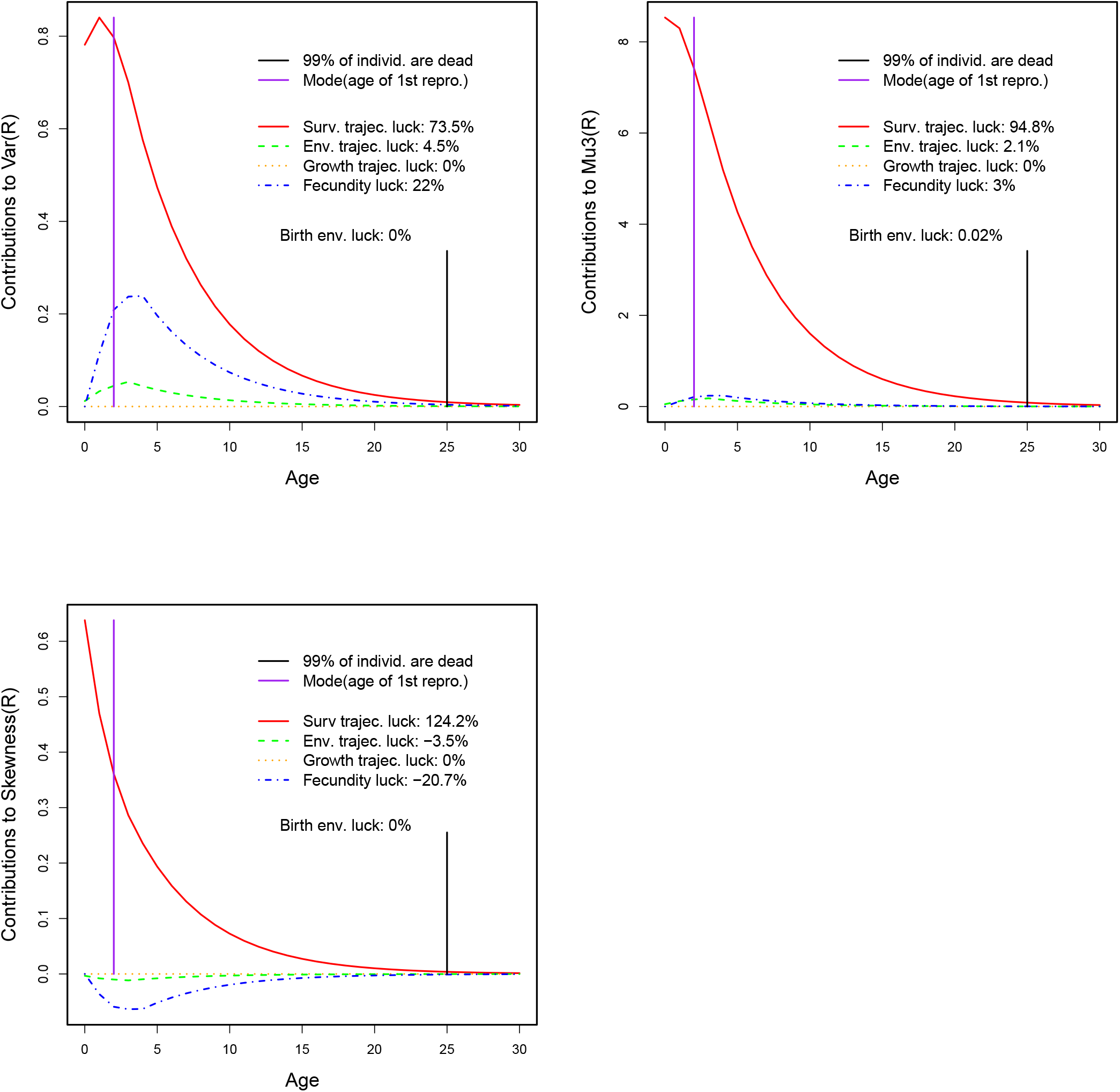
Variance, third central moment, and skewness of lifetime reproductive output for snow geese. Lines show the contributions of different forms of luck as a function of age. Top left: Partition of variance. Top right: Partition of third central moment. Bottom: Partition of skewness. The percent contributions are calculated with respect to total skewness, not the sum of the absolute magnitudes of contributions. Generated by `snowgooseVarMu3Partition.R` and `snowgooseVarSkewnessPartition3.R`.

**Figure 3:**
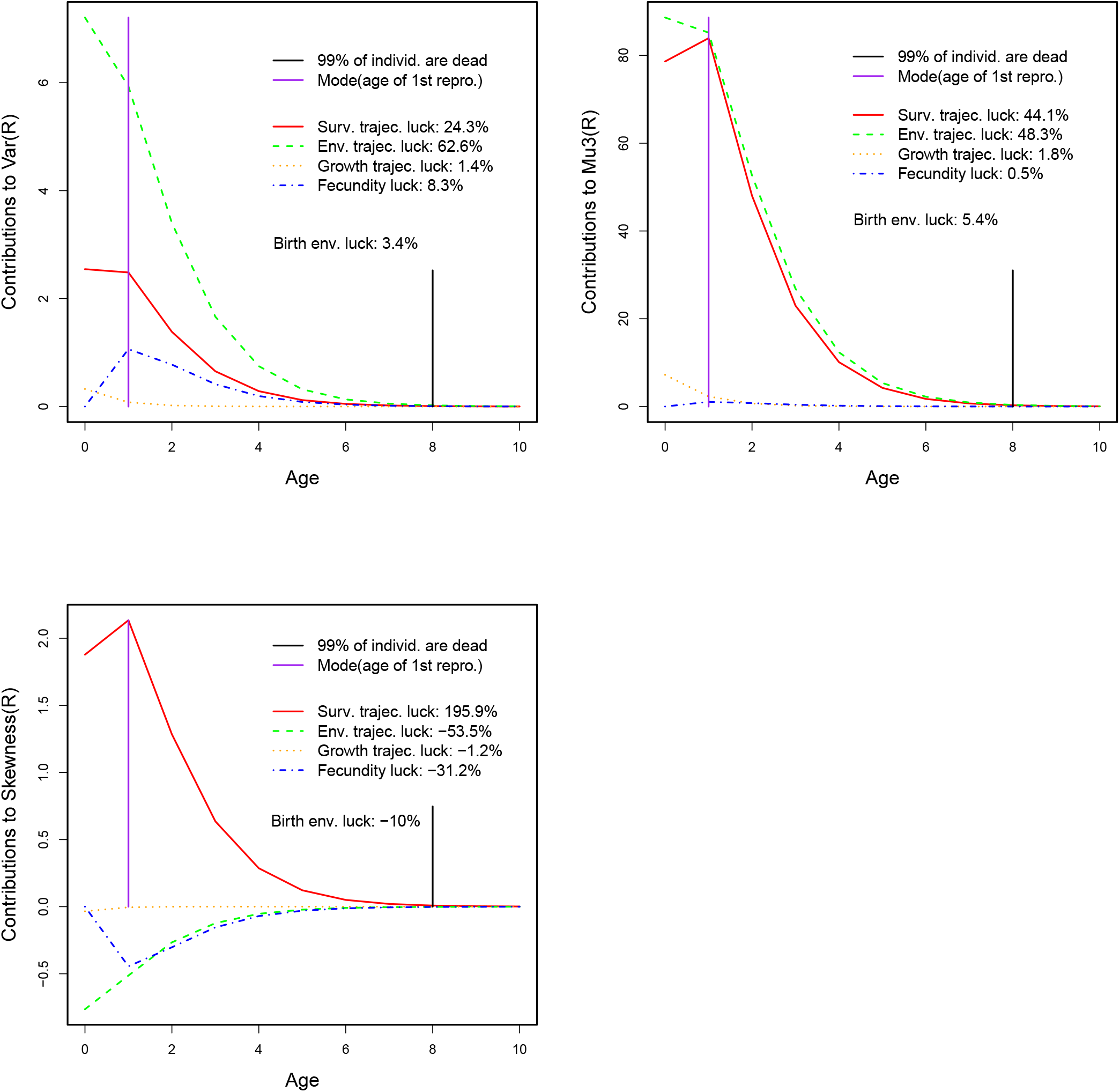
Variance, third central moment, and skewness of lifetime reproductive output for sculpins. Lines show the contributions of different forms of luck as a function of age. Top left: Partition of variance. Top right: Partition of third central moment. Bottom: Partition of skewness. The percent contributions are calculated with respect to total skewness, not the sum of the absolute magnitudes of contributions. Generated by `sculpinVarSkewnessPartition2.R` and `sculpinVarMu3Partition.R`.

Because we found that reproductive skew is largely driven by luck in survival, we assumed that having exceptionally large LRO is mostly a matter of living exceptionally long. To characterize the connection between reproductive success and lifespan, we calculate the distribution of lifespan conditional on achieving a particular LRO: how long do those in the reproductive tail live? The first step is to expand the model’s state space so that individual state includes the individual’s total reproductive output up to the present. Defining “success” as having some number (or at least some number) of offspring, we calculate the state transition probabilities conditional on achieving success (Kemeny and Snell, 1960; Snyder and Ellner, 2016). By iterating the state-transition matrix we obtain the probability of survival to ages 1,2,3,*…* and therefore the distribution of lifespan conditional on being successful.

The rest of this section fills in the mathematical details and provides some computing formulas. If you want those right now, read on. If not, you can safely skip ahead to the *Results from case studies* section.

### Notation and assumptions

We use the notation that 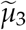 denotes skewness, while *µ*_3_ is the unscaled third *central* moment. Skewness is a normalized form of the third central moment that provides a scale-free measure of the asymmetry of a probability distribution: for a random variable *X*, 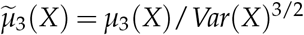. Because we use skewness as a measure of a distribution’s asymmetry, we define skewness to equal zero when the variance is zero and the standard formula gives 0/0 (undefined).

As always in a discrete time model, it is necessary to specify the sequence of events within a time step. Here we assume the order shown in Fig. 1: between ages *a* and age *a*+1, living individuals experience first reproduction, then risk of mortality, then individual state transition (“growth”), and finally an update in the environment state (if that is present in the model). This diagram assumes a pre-breeding census; a post-breeding census would have reproduction occurring last in each time step, rather than first. In SI section S3 we describe the relatively minor changes needed for a post-breeding census. It is important to note that individuals born within one time step only join the population at the start of the next time step, as new age-0 individuals, and after that undergo the depicted sequence of events every year.

We define

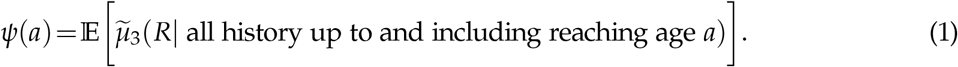

On the right-hand side, the 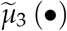 in square brackets denotes the skewness of the conditional distribution of lifetime reproductive output *R* given the individual’s entire history up to and including reaching age *a*, including past states and realized clutch sizes.^3^ The expectation is the average of that quantity, across the probability distribution of individual histories up to and including reaching age *a*. Conditioning *R* on past reproductive output, in addition to past states, differs from our past work.

The variance of a sum of independent variables is the sum of the variances, and we assume that clutch sizes at different ages are independent conditional on the state trajectory, so the contribution of fecundity luck to *Var*(*R*) is just the sum of age-specific clutch size variances. Higher moments of sums do not break down so neatly, so we need to calculate the contribution of fecundity luck age by age, just like the other forms of luck. Conditioning on past reproductive output as well as past states keeps the past and future independent (eq. 4). To that end, we let *B*_*a*_ denote clutch size (“births”) at age *a* so that

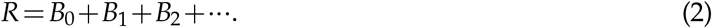

Below, we use *B*_*a*_ to denote births at each age considered as a random variable and *b*_*a*_ to denote the realized number of births in a particular life trajectory. Similarly *Z*_*a*_ denotes individual state at each age considered as a random value and *z*_*a*_ the realized state; we use *Q*_*a*_, *q*_*a*_ in the same way for the environment state when that is present in the model and *Y*_*a*_, *y*_*a*_ for the vector of all model state variables (*z, q*, and any other state variables in the model). For some of our calculations the individual state needs to include *k*, the individual’s total reproductive output up to the present. Including *k* greatly increases the size of transition probability matrices and creates new opportunities for indexing errors, so it should be done only when it is essential to answer the question at hand.

We assume that transitions between environments states, if they occur, are Markovian. The environment can be temporally autocorrelated, but each year’s environment can only depend on the previous year’s environment.

### Age-partitioning of skewness contributions from different luck types

At birth, an age 0 individual already has some history: their birth state *z*_0_ and birth environment *q*_0_. We use *ψ*(−1) to denote the unconditional skewness 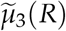 — the skewness of LRO *R* at an imaginary pre-birth state before birth state and birth environment have been assigned. If there are multiple possible birth states and/or birth environments, *ψ*(−1) needs to be computed using the law of total cumulance and the vector of conditional skewness given birth state and environment (see Supplementary Information section *Computing contributions to skewness from trait value, birth state, and birth environment*).

Our decomposition of LRO skewness is the telescoping sum identity

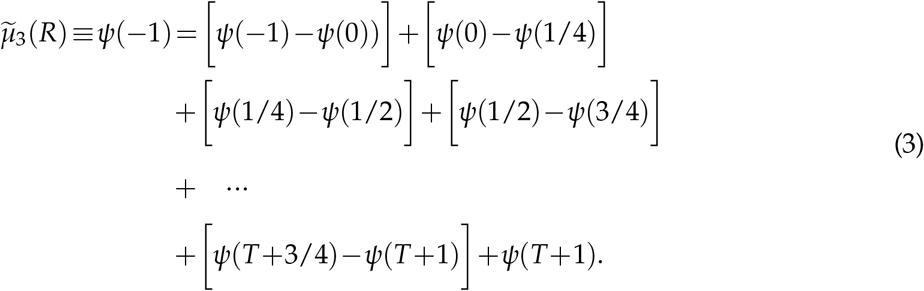

As *T* → ∞ the remainder *ψ*(*T* +1) → 0, because the conditional distribution in eqn. (1) is dominated by the probability that the individual dies before age *T*. If that happens, then *R* = *b*_1_ + *b*_2_ + *…* + *b*_*T*_ which is conditionally constant at *T*+1, with zero skewness. *ψ*(*T*+1) therefore cannot be larger than the probability that the individual is still alive at *T*, times the (finite) maximum skewness of *R* as a function of initial state. Thus *ψ*(*t*) → 0 at least geometrically fast, so it is safe to neglect terms in (1) beyond the age at which nearly all individuals are dead.

In eqn. (3) the first difference in square brackets on the right hand side is the change in expected skewness resulting from knowing birth state *z*_0_ and birth environment *q*_0_. The second term is the change resulting from knowing the first realized fertility *f*_0_. The third is the change resulting from knowing whether the individual survived to time step 1 — and so on.

We now need to evaluate each *ψ*(*s*). To do that, we create an extended state space model in which the different demographic transitions — fecundity update, survival update, growth update, environment update — occur separately and sequentially within a unit of “clock time” that advances individual age from *a* to *a*+1. Going from age *a* to age *a*+1 now takes four intermediate steps. For maximum generality, we assume here that individuals are characterized by an individual state vector *y* consisting of their state (“size”) *z*, environment *q*, and number of offspring up to the current time *k* (“kids”), and indicate the changes for models where *k* or *q* may be absent from *y*.

Adding a constant to a random variable does not change its skewness. Thus, the skewness of lifetime total reproduction *R* conditional on past reproduction *b*_0_,*b*_1_,*…*,*b*_*a*_ and states *y*_0_,*y*_1_,*…*,*y*_*a*_ is

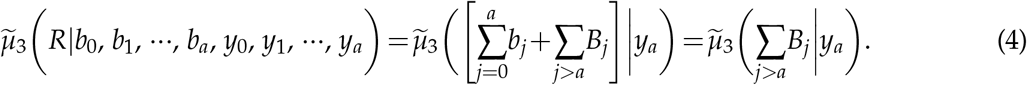

The important aspect of eqn. (4) is that even though *R* includes past as well as future reproduction, the skewness of *R* conditional on an individual’s history up to the present depends only on future reproductive success, not on past reproductive success. The same is true for any intermediate point in our subdivision of the time step. And, computing the skewness of future LRO conditional on current state in our extended state space is a solved problem (van Daalen and Caswell, 2017). Those calculations do not require *k* (reproduction to date) to be a component of *y* because *z*_*a*_ and *q*_*a*_ together determine the probability distribution of future states and reproductive outputs.

Let **F** and **P** denote the fecundity and state transition (i.e., survival & growth) matrices, respectively, of the original projection matrix **A** = **P**+**F**. Moments of LRO are computed from **P** and the “reward matrices” **R**_*j*_ (having the same size as **A**) giving the *j*^*th*^ non-central moments of the “reward” (number of offspring) associated with each possible transition, following the methods described in van Daalen and Caswell (2017). Reward matrices are a means to calculate the non-central moments of the LRO distribution. The *i, j*th entry of a reward matrix represents the “reward” (e.g. mean reproduction, second non-central moment) associated with the transition from state *j* to state *i*. The **R**_*j*_ are calculated from **F** using assumptions about the form of the offspring number distribution, where the column sums of **F** give the mean of the distribution as function of parent state. We assume that the offspring number distribution is determined by current state only, not by current and subsequent state^4^, so that, assuming the *i*,*k*^*th*^ element of a matrix represents transitions from state *k* to state *i*, all entries in the *k*th column of **R**_*j*_ are equal to the *j*th moment of the clutch size for a parent in state *k*.

Our extended state space has dimension 4 *×* the original dimension, to accommodate the four phases of the time step. Only living states are included, so that we can directly use formulas for that situation from van Daalen and Caswell (2017). How we nonetheless account for the dead is explained after eqn. (7). If the original number of living states is *n*, the first *n* entries of the state vector are devoted to the state of the individual at the start of the time period, the second *n* entries to the state of the individual just after the fecundity update, the third *n* entries to the state just after the survival update, and the last *n* entries to the state just after the growth update.

To create the transition and reward matrices on the extended state space, we make the following definitions, also used in Snyder and Ellner (2022)^5^:

1. **F**_*•*_ is the matrix (or discretized integral projection model (IPM) kernel) that only updates cumulative offspring number *k*. If cumulative offspring number is part of the state variable, the entries of **F**_*•*_ involve the probability distribution for clutch size, the mean of which is taken from the fecundity matrix **F**. If cumulative offspring number is not part of the state variable in the original model, then **F**_*•*_ is the identity matrix.
2. **S**_*•*_ is the matrix that only updates survival versus death – there is no change in individual state *z*. This is a diagonal matrix with state- and environment-specific survival probabilities on the diagonal.
3. **G**_*•*_ is the matrix that only includes growth (i.e., individual state transitions), conditional on survival.
4. **Q**_*•*_ is the matrix that only updates the environment state.

Note that **Q**_*•*_**G**_*•*_**S**_*•*_**F**_*•*_ = **A**. The transition matrix on the extended state space is then

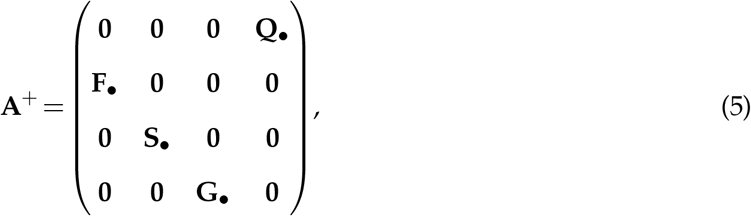

so that **Q**_*•*_, **F**_*•*_, etc. each act only on one *n*-component block of the state vector, and each time we multiply by **A**^+^, we move the non-zero components of the state vector to the next *n*-component block, corresponding to updating the system in one of the four intermediate steps portrayed in Fig. 1. E.g., applying **A**^+^ to the state at the beginning of the time step updates fecundity and moves the non-zero portions of the state vector from the first *n* components to the second *n* components: **A**^+^((*k*_*a*_,*z*_*a*_,*q*_*a*_),**0, 0, 0**) =(**0**, (*k*_*a*+1_,*z*_*a*_,*q*_*a*_), **0, 0**). The corresponding reward matrices are

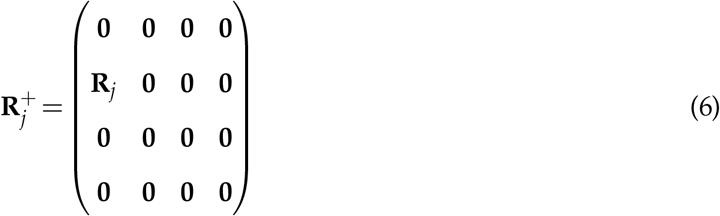

Now we can evaluate skewness at various points in the update from age *a* to age *a*+1 — i.e. the various *ψ*(*a*). We let *y* denote a possible state in the extended state space, and *y*_0_ a possible state at birth. 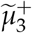 will denote skewness of *R* as a function of state on the extended state space (and hence potentially finding skewness partway through a time step), and 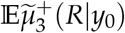 is the expected skewness of *R* conditional on initial state on the extended state space being *y*_0_.

1. The expected skewness at the start of a time period is 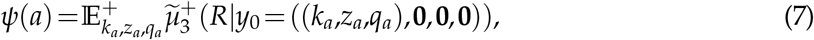 where we use 𝔼^+^ to denote a weighted average over the living states in which the weights sum to the probability of being alive at that age. The distribution of living states (*k*_*a*_,*z*_*a*_,*q*_*a*_) is **A**^*a*^*m*_0_, where *m*_0_ is the initial state distribution, so 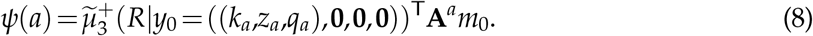
2. The expected skewness just after the fecundity update is 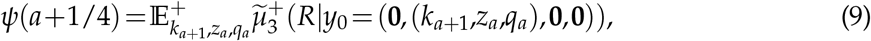 where the distribution of living states (*k*_*a*+1_,*z*_*a*_,*q*_*a*_) is **F**_*•*_**A**^*a*^*m*_0_.
3. The expected skewness just after the survival update is 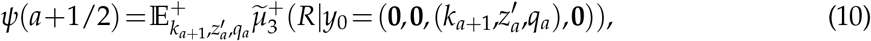 where the distribution of living states 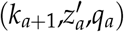 is **S**_*•*_**F**_*•*_**A**^*a*^*m*_0_.
4. The expected skewness just after the growth update is 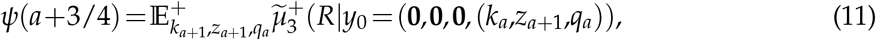 where the distribution of living states (*k*_*a*+1_,*z*_*a*+1_,*q*_*a*_) is **G**_*•*_**S**_*•*_**F**_*•*_**A**^*a*^*m*_0_.
5. The expected skewness just after the environment update is the expected skewness at the start of the next time period, *ψ*(*a*+1).

Fecundity luck, survival trajectory luck, growth trajectory luck, and environment trajectory luck are the changes in expected skewness when we make the relevant demographic transition. So, for example, survival trajectory luck at age *a* is the change in expected skewness from just after the fecundity update at age *a* to just after the survival update at age *a*: *ψ*(*a*+1/4)−*ψ*(*a*+1/2).

The extended-state transition matrix **A**^+^ corresponds to the order of events within a time-step assumed in Fig. 1, but the approach works for any order with minor changes. For example, with the order growth → reproduction → mortality → environment update, we have

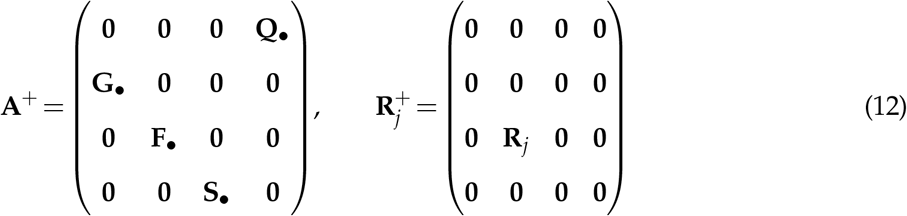

and the living state distributions are **G**_*•*_**A**^*a*^*m*_0_, **F**_*•*_**G**_*•*_**A**^*a*^*m*_0_, and **S**_*•*_**F**_*•*_**G**_*•*_**A**^*a*^*m*_0_, respectively after the growth, fecundity, and mortality updates.

For models without environmental variation, the only difference is that the time step is divided into three intermediate steps, updating fecundity, survival, and growth. We walk the reader through this process in Supplementary Information section *Models without environmental variation*.

### Birth state and birth environment luck

In the partition (3), *ψ*(−1)−*ψ*(0) is the change in expected skew resulting from an individual being assigned a birth state *z*_0_ and a birth environment *q*_0_. It represents the total effect of birth state luck and birth environment luck, that is

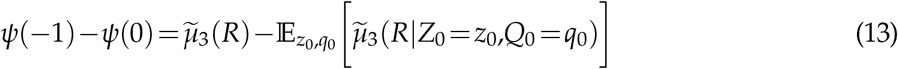

where the expectation is over the bivariate distribution of *z*_0_ and *q*_0_.

If birth state and birth environment are assigned independently, then *ψ*(−1) − *ψ*(0) can be partitioned into separate contributions from birth state luck and birth environment luck. This will always be the case if the environment is independent and identically distributed rather than actually depending on past environments. If the environment is not independent and identically distributed, it will be true if there is exact mixing at birth, that is, if the offspring state distribution is the same for all parents in all environment states.

However, in other cases there is no natural way of choosing an “order of events” for assignment of birth state and birth environment. Our assumed order of events within time-steps (Fig. 1) has reproduction occurring before an environment update, so for our case studies here we assume that birth state is assigned first, then birth environment. The skewness contributions from birth state *z* and birth environment *q* are then, respectively,

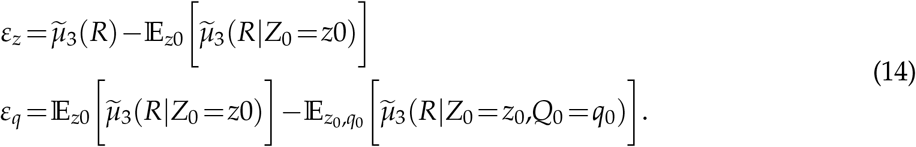

Our calculations take as “given” the distribution of birth state and environment, but in some populations either or both of those may depend on the state and environment of the individual’s parent, say if good environments or more robust parents yield larger offspring on average. In such situations the user has several options. They can use the distribution of offspring state and environment for a typical parent (e.g., the parent of a randomly chosen newborn when the population is at its stationary stage/state/environment distribution); they can evaluate and compare the birth state and environment contributions for different types of parent in different environments; or they can extend our analysis by adding an age *a* = −2 at which the state and environment of the individual’s parent is chosen from some relevant distribution.

### Determining the contribution of traits

If the original model includes variation among individuals in fixed traits (which we denote *x*), we measure the total contribution of traits to skewness by letting trait assignment be the first event represented in our telescoping sum. That is,

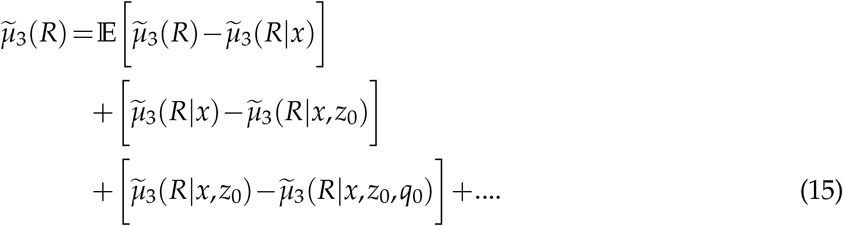

To perform this calculation, we suppose that everyone is born into an undifferentiated ur-state, as we did for birth state and birth environment. Individuals first acquire a trait value, then an initial size/stage, then an initial environment. The change in skewness that comes from conditioning on the trait value is the total contribution of traits to skewness.

The ur-state skewness *ψ*(−1) and the terms in the decomposition of *ψ*(−1)−*ψ*(0) are easiest to calculate by using an extended state space that includes pre-birth states. This is conceptually like the extended transition matrix eqn. (5), in that individuals sequentially acquire their trait value, initial state, and initial environment; the details are given the Supporting Material, section *Computing contributions to skewness from trait value, birth state, and birth environment*.

### Calculating the lifespan distribution conditional on LRO

We will find that survival trajectory luck dominates contributions to the skewness of LRO in our empirical case-studies. To determine if those with exceptional LRO are special in part because they live exceptionally long, we will want to calculate the lifespan distribution conditional on LRO. To do this, we again subdivide the time step exactly as in Fig. 1. Although we are not partitioning luck, the order of updates still affects our answer. This subdivision allows us to ensure that reproduction happens before survival, as we have assumed. The individual state space is expanded to include *k*, the number of offspring produced up to the present. For example, if individuals were originally characterized by size *z* and environment *q*, they are now characterized by *z, q*, and *k*. We then create **F**_*•*_, **S**_*•*_, **G**_*•*_, and **Q**_*•*_ matrices for this new, extended-state kernel and assemble them into transition matrix **A**^+^ as in eq. 5. LRO is then an aspect of the individual’s state at death, and we can use previously developed methods for conditioning transition probabilities on state at death (Snyder and Ellner, 2016).

Specifically, we calculate **A**^+^ conditional on “success,” where success is defined as having a given LRO, *R*_*T*_. We do this by first defining a modified transition kernel with two absorbing states — either an individual dies with a value of *k* equal to *R*_*T*_ or higher, or they die with *k*<*R*_*T*_. We calculate (by standard methods) the probability of ending in the former absorbing state given that an individuals’ current state is a transient (non-absorbing) state *z, q*_*s*_(*z*; *R*_*T*_). The probability of having LRO of precisely *R*_*T*_ is then

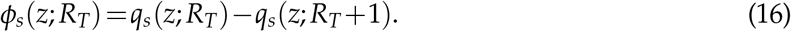

The kernel conditional on “success” is then

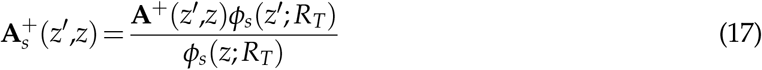

(Iosifescu, 1980, Ch. 3), where we have returned to writing the kernel as a function of the current state *z* and the state at the next time step, *z*^*t*^. Details of these calculations can be found in Snyder and Ellner (2016).

To determine the lifespan distribution conditional on success, we repeatedly multiply the initial state distribution conditional on success by the conditional kernel 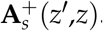. The initial distribution conditional on success is

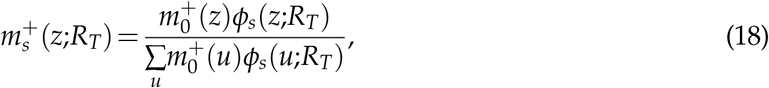

where 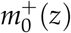 is the (unconditional) distribution of initial states on the expanded state space (for IPMs the sum in the denominator is replaced by an integral). The probability that an individual survives to *at least* age *L*, conditional on having an LRO of *R*_*T*_ is then

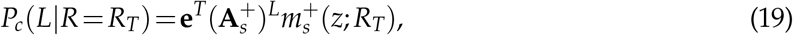

where **e** is a column of 1s. The conditional probability of having a lifespan of precisely *L* years is then

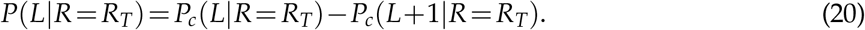

## Results from case studies

We have applied our methods to several case studies chosen to span a wide range of life history attributes. Basic life history information about each of the species and populations is presented in Table 1. Some of our case study organisms have slow, steady reproductive strategies — small clutch sizes that do not vary much with environmental conditions — including fulmars (a long-lived sea bird) and snow geese. For others, fecundity varies moderately or strongly with the environment, e.g. sculpins and *Umbonium*, a sea snail. Our case studies include long-lived organisms like fulmars and kittiwakes and short-lived organisms like sculpins and *Lomatium*, a perennial plant. In two of the models (kittiwakes and fulmars) individuals are cross-classified by an unchanging trait. In kittiwakes the trait is a latent “quality” variable, with high-quality individuals having both higher adult survival and higher adult breeding probability. Fulmars are grouped into three fixed behavioral syndromes governing survival, breeding probability, and breeding success.

**Table 1:** Life history characteristics of case study species. SI indicates a species used only in the Supplementary Material. Location is specified if the originating data base contains models for multiple locations. “Expected clutch sizes” is the range of expected clutch sizes across environment states for the most fecund stage.

| Name | Expected clutch sizes<br>of most fecund stage | Mean lifespan |
| --- | --- | --- |
| Snow goose *Anser caerulescens* | 0.155–0.975 | 4.6 |
| Woolly sculpin, False Point population<br>(*Clinocottus analis*) | 0.02–21.5 | 1.3 |
| Southern fulmar, *Fulmarus glacioides* | 1 | 15.0 |
| Black-legged kittiwake, *Rissa tridactyla* | 2.5 | 10.8 |
| *Umbonium costatum* (marine snail with planktonic larvae) | 0.04–6.15 | 3.7 |
| SI: Lesser kestrel *Falco naumanni* | 0.1 – 0.45 | 2.4 |
| SI: *Lomatium bradshawii* from Rose Prairie site 1 | 0–2.7 | 2.2 |
| SI: Flathead knob-scaled lizard<br>from tropical site (*Xenosaurus platyceps*) | 2.45 | 2.0 |

Our kittiwake model combines a model for average vital rates from Steiner et al. (2010), with estimates of trait variation and its effects from Cam et al. (2002). We make the conservative assumption that breeding probability and survival are perfectly deterministic functions of individual quality, and are therefore perfectly correlated — this is conservative for our purposes because it maximizes the potential impact of trait variation. The fulmar model comes from Jenouvrier et al. (2022). The other models are taken from the COMADRE and COMPADRE databases (Salguero-Gómez et al., 2016; Salguero-Gómez et al., 2015).

We are missing information on clutch size distributions for most species. Fulmars produce one chick if they breed successfully, and kittiwakes produce one chick in their lower fecundity stage or two or three chicks (assumed to have equal probability) in their higher fecundity stage. For all other case studies, we assume that annual clutch size follows a Poisson distribution, with reported fecundities representing expected values.

### What produces the tail in LRO: partitioning skewness

#### Reproductive skew is driven mostly by luck, not inherent differences between individuals

In previous work we and others found that reproductive variance is dominated by various forms of luck, not trait variation (see the *Introduction*). The same conclusion holds true for skewness in our current case studies. If we consider the absolute magnitudes of contributions to skewness, trait variation represents only 16% of the total contributions for the fulmars and 31% for the kittiwakes (Figs. 4 and 5). Note that trait variation can increase or decrease reproductive skew.

**Figure 4:**
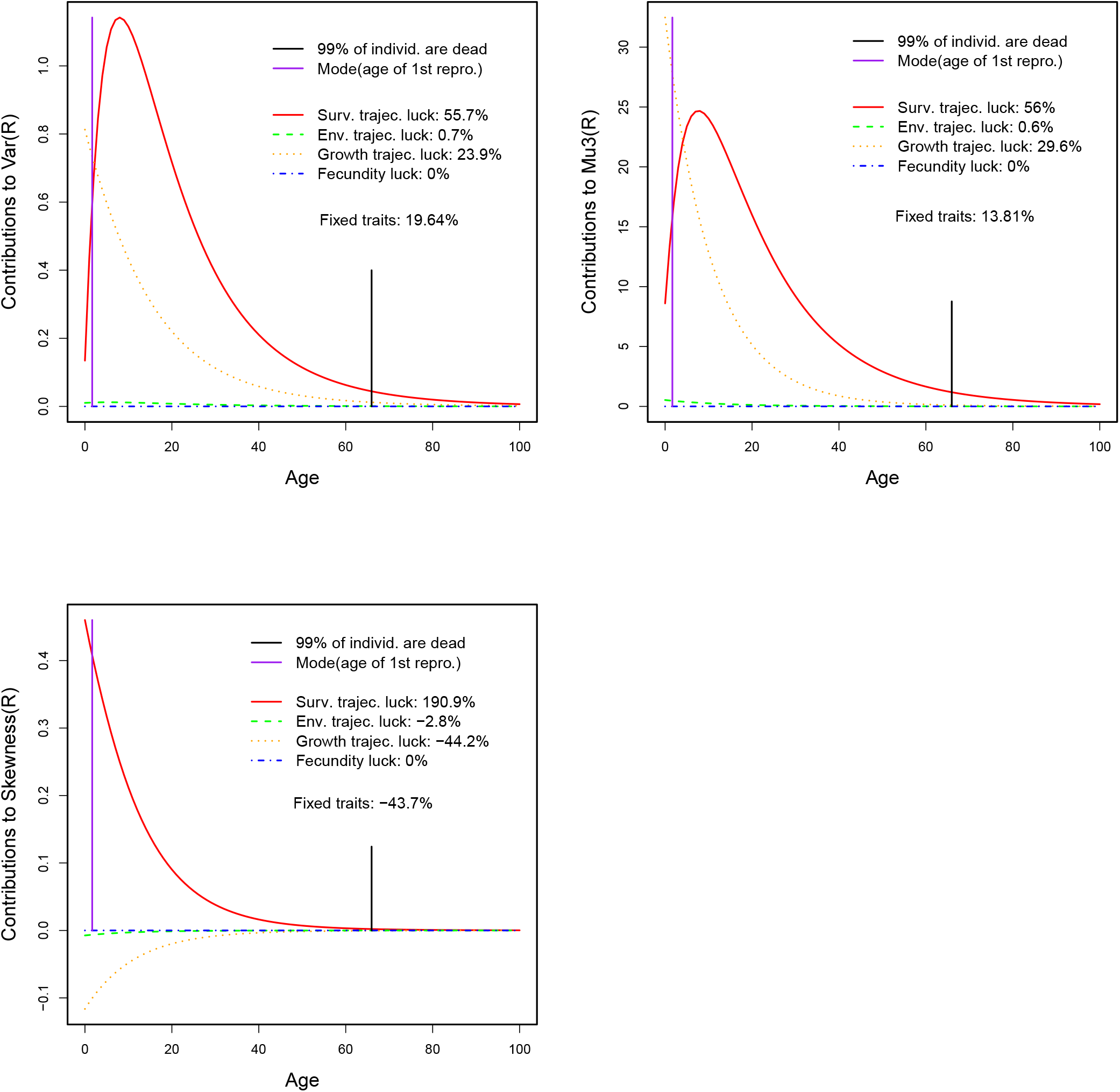
Variance, third central moment, and skewness of lifetime reproductive output for fulmars. Lines show the contributions of different forms of luck as a function of age. Top left: Partition of variance. Top right: Partition of third central moment. Bottom: Partition of skewness. The percent contributions are calculated with respect to total skewness, not the sum of the absolute magnitudes of contributions. Note that we can calculate only the total contribution of fixed traits, not an age partition. Generated by `fulmarVarSkewnessPartition.R` and `fulmarVarMu3Partition.R`.

**Figure 5:**
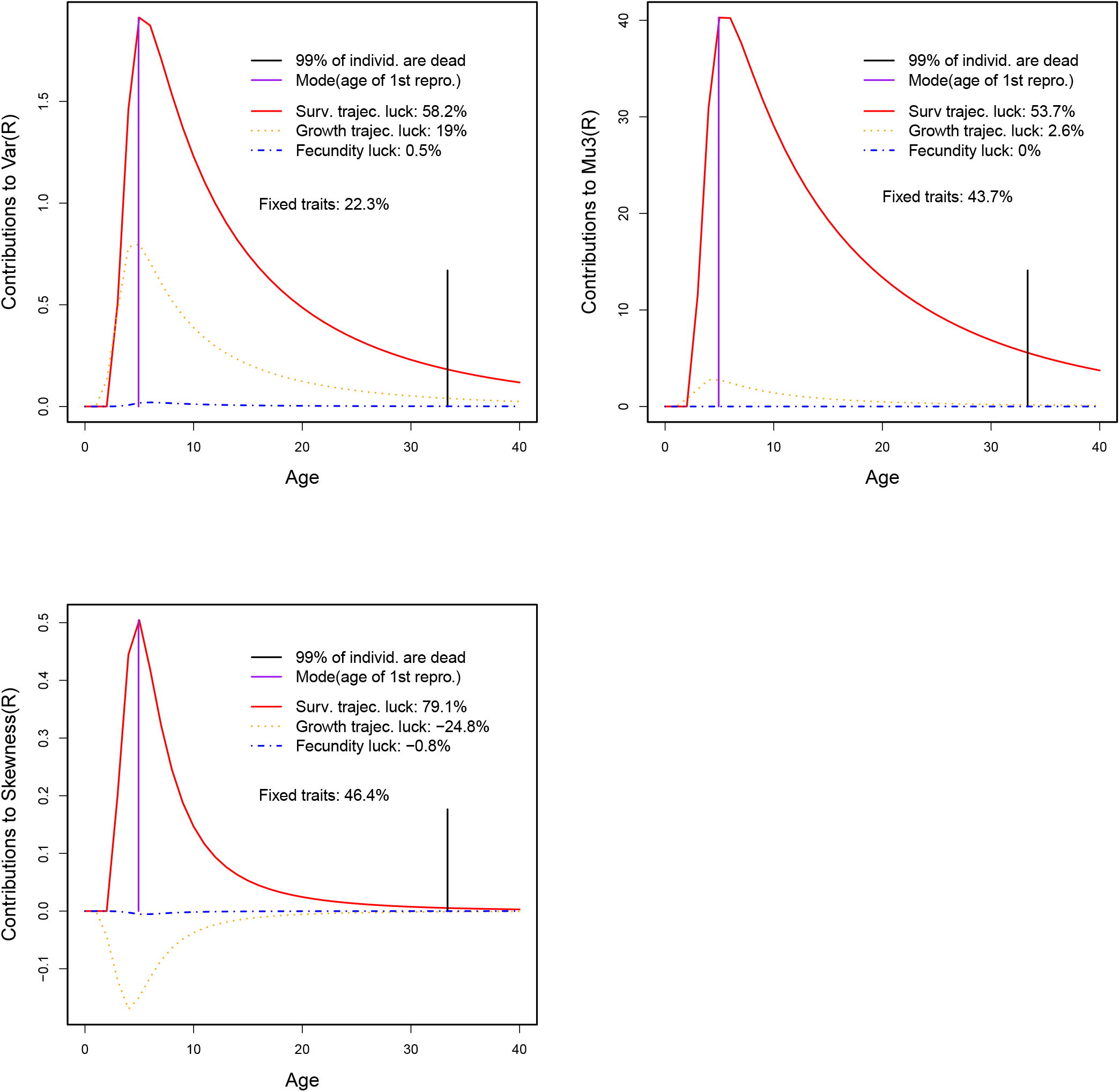
Variance, third central moment, and skewness of lifetime reproductive output for kittiwakes. Lines show the contributions of different forms of luck as a function of age. Top left: Partition of variance. Top right: Partition of third central moment. Bottom: Partition of skewness. The percent contributions are calculated with respect to total skewness, not the sum of the absolute magnitudes of contributions. Note that we can calculate only the total contribution of fixed traits, not an age partition. Generated by `kittiwakeVarSkewnessPartition.R` and `kittiwakeMu3SkewnessPartition.R`.

### Unlike reproductive variance, reproductive skew is driven mostly by variation in lifespan

The dominant contribution to reproductive variance, *Var*(*R*), depends on life history. For organisms with especially labile growth, such as a shrub that can grow or shrink, reproductive variance may be driven by growth trajectory luck (Snyder and Ellner, 2022), while environment trajectory luck typically drives reproductive variance for those whose demographic transitions depend strongly on environmental conditions (e.g. Figs 3 and S4). Survival trajectory luck dominates reproductive variance for organisms with slow, steady growth and reproduction, such as trees (Snyder and Ellner, 2022) and long-lived birds that produce 1–3 chicks per year (e.g. Figs. 2, 4, 5). In contrast, skewness is always driven by survival trajectory luck: compare the variance and skewness panels in Figs. 2–5, S2–S5. Considering the absolute magnitude of contributions to skewness, the second largest contribution in our case studies was at most 59% that of survival trajectory luck (trait variation in kittiwakes) and most were substantially smaller (Figs. 2–5, S2–S5). Survival trajectory luck represents the contributions that come from an individual dying or surviving at each age, so the dominance of survival trajectory luck contributions means that reproductive skew is driven by randomness in lifespan.

#### Differences in lifespan increase reproductive skew, while other forms of luck decrease it

Growth trajectory luck, environment trajectory luck, fecundity luck, and birth state and environment luck can all increase the third central moment of LRO, and in most cases they do, at least to some extent (Figs. 2–5, S2–S5). However, they also increase the variance of LRO, by enough so that skewness (which is the third central moment normalized by variance^3/2^) is actually decreased. So among all the different forms of luck, only survival trajectory luck contributes positively to reproductive skew.

### What does it take to get into the LRO tail?

Partitioning reproductive skew gives us a sense of what drives the lopsidedness in LRO, but that is not the same as asking what it takes to be extremely successful. What can we infer about individuals with extremely high LRO?

The dominant contribution of survival trajectory luck to skewness suggests that a long life is key to success. Clearly, if an individual can have at most one offspring per year, then the only way to have a large LRO is to live a long time, reproducing many times. But other forms of luck may be able to partially substitute for luck in survival. If a species has Poisson-distributed clutch sizes with a large mean (and hence a large variance) or if clutch sizes in good years are much larger than those in other years, then an individual might achieve extreme success either by living a long time, or by not living quite as long but instead getting lucky with fecundity or environment.

To explore what we can infer about the extremely successful, we calculated the distribution of lifespan conditional on LRO. Fig. 6 shows plots for snow geese, *Umbonium*, and sculpins. Snow geese have small clutches in all years (the expected clutch size in the most fecund state varies between 0.155 and 0.975); *Umbonium* has moderate environmental variation in fecundity (maximum expected clutch size varies between 0.04 and 6.15); and sculpins are wildly fecund in good years and rarely reproduce in bad years (maximum expected clutch size varies between 0.02 and 21.5). Given their slow, steady reproductive strategy, the only way for snow geese to achieve exceptional reproductive success is to live exceptionally long, and this is reflected in the relatively tight distribution of lifespans. For example, if we know that an individual had an LRO in the 95th percentile, we are 90% certain that its lifespan was between the 83rd and 97th percentiles. On the other hand, reproductive success implies less about the lifespan of *Umbonium* individuals. An *Umbonium* with LRO in the 95th percentile is 90% likely to have a lifespan between the 74th and 97th percentiles. This wider central interval occurs in part because *Umbonium* can achieve success by experiencing a good year, rather than by having an extra reproductive bout or two in average years. Sculpins have such wildly varying fecundity that there are two routes to exceptional success. Either an individual can live a long time, possibly substituting a good year for some longevity, or they can get one of the rare exceptionally good years when they first hit reproductive maturity (as early as age 1), and have the average number of offspring for that stage and year type, which is just shy of 15 in the best years. This possibility of hitting the environmental jackpot at age 1 is what produces the second red area at lifespan = 2 and LRO equal to 13–15 (newborn individuals are age 0, so dying at age 1 means that lifespan = 2); other routes to LRO in that range are very unlikely, requiring a long life with several (also uncommon) medium-quality years. If we eliminate environmental variation in fecundity, the corresponding heat-map for sculpins is very different (Fig. S6).

**Figure 6:**
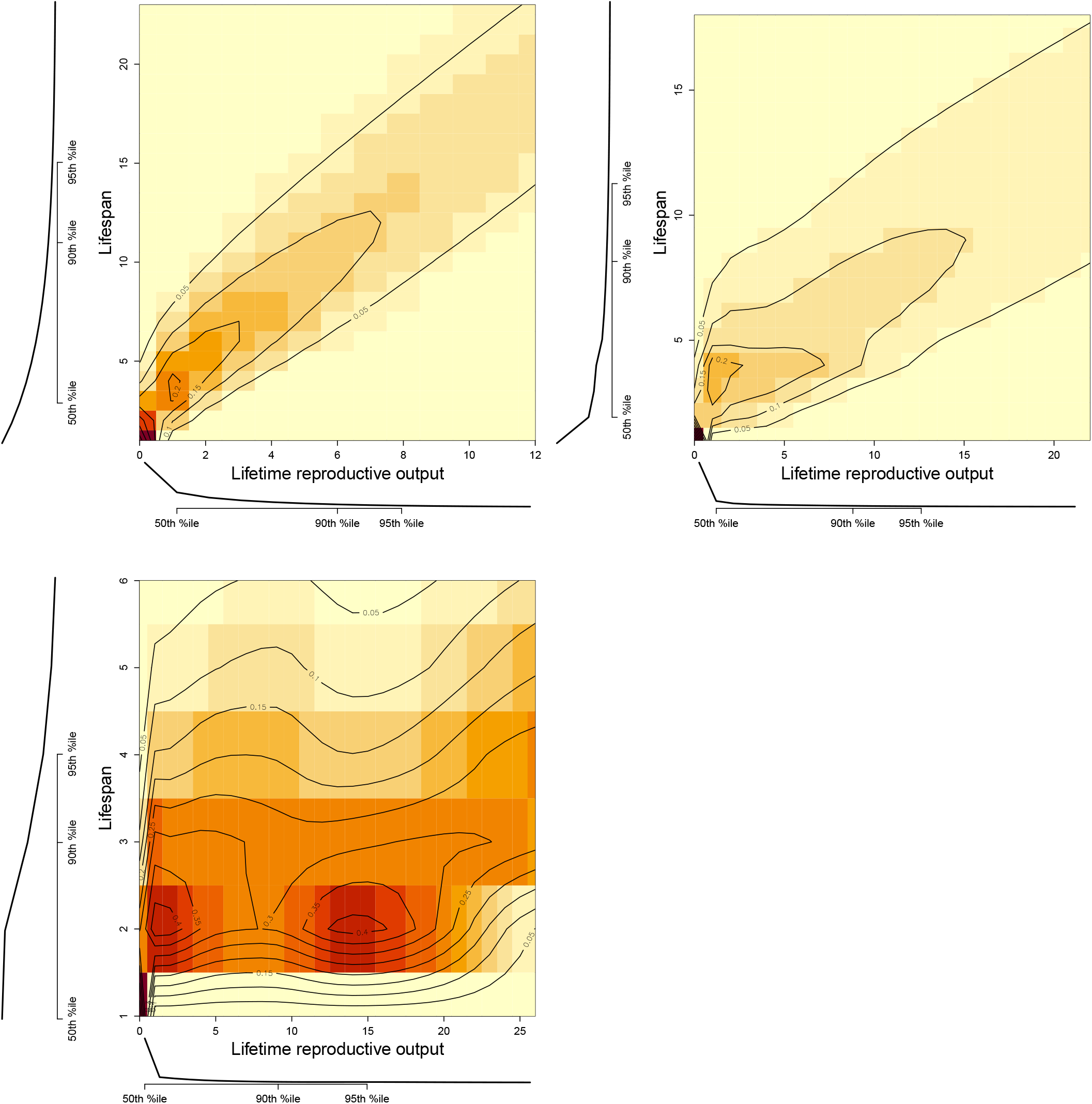
*Pr*(Lifespan | *R*) for snow geese (top left), *Umbonium* (tossp right), and sculpins (bottom right). Note that an individual who dies before reaching age 1 is assigned a lifespan of 1. Generated by `plotSnowgooseLifespanCondLRO3.R`, `plotUmboniumLifespanCondLRO2.R` and `plotSculpinLifespanCondLRO3.R`, which get data from `snowgooseLifespanCondLRO.R`, `umboniumLifespanCondLRO.R`, and `sculpinLifespanCondLRO.R`.

To further explore the effect of environmental variation in fecundity, we altered the demographic parameters of the *Umbonium* model. Fig. 7 shows the 90% central intervals for lifespan for *Umbonium* with environmental variation in fecundity (yellow) and without (orange) (fecundity in each stage is set to its environmental average). Eliminating the possibility of hitting a jackpot year does indeed narrow the central interval: if you can’t achieve extreme success by reproducing in an exceptionally good year, you have to do it by living longer.

**Figure 7:**
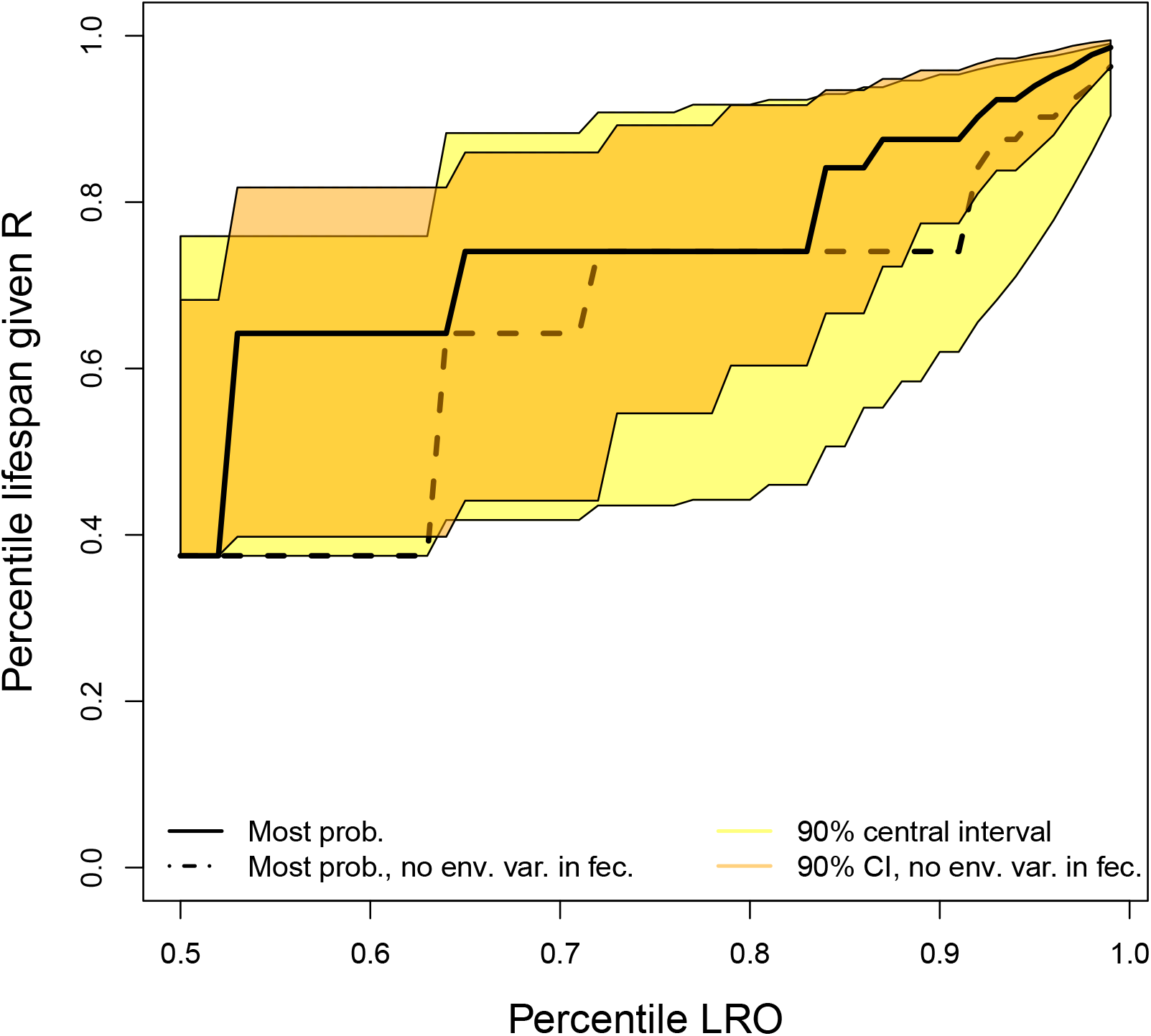
90% central intervals for *Pr*(Lifespan | *R*) for *Umbonium* with environmental variation in fecundity (expected clutch size for most fecund stage 0.04–6.15) and when expected clutch size for each stage is set to its environmental average. Generated by `compareUmboniumLifespanCIVsLROP-ercentile.R`, using data from `umboniumLifespanCIVsLROPercentile.R`, which gets data from `umboniumLifespanCondLRO.R`.

It is also the case that species with shorter lifespans (like sculpins) have wider central intervals for conditional lifespan. Similarly, the 90% central interval for kestrels (yellow, in Fig. 8) becomes narrower if we modify the model to have 20% higher survival in all stages (orange). In Supplementary Material section *Mean lifespan and central intervals for lifespan given LRO*, we show that an inverse relationship between mean lifespan and width of central intervals for conditional lifespan (on a percentile scale) is an expected consequence of a property of the models we are using — specifically, the property that age-dependent average survival and fecundity converge with increasing age to unchanging asymptotic values (Cochran and Ellner, 1992). This convergence necessarily results from convergence to the stable state distribution of the state transition matrix or kernel, and convergence of age-dependent survival is often even faster, because survival varies relatively little among states typical of adults. As a result, lifespan (or adult lifespan) typically is roughly geometrically distributed. In the idealized situation where lifespan is geometrically distributed, and mean fecundity is age-independent, the width of a conditional central interval for lifespan given LRO goes down with mean lifespan roughly in proportion to 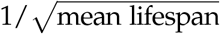, all else being equal.

**Figure 8:**
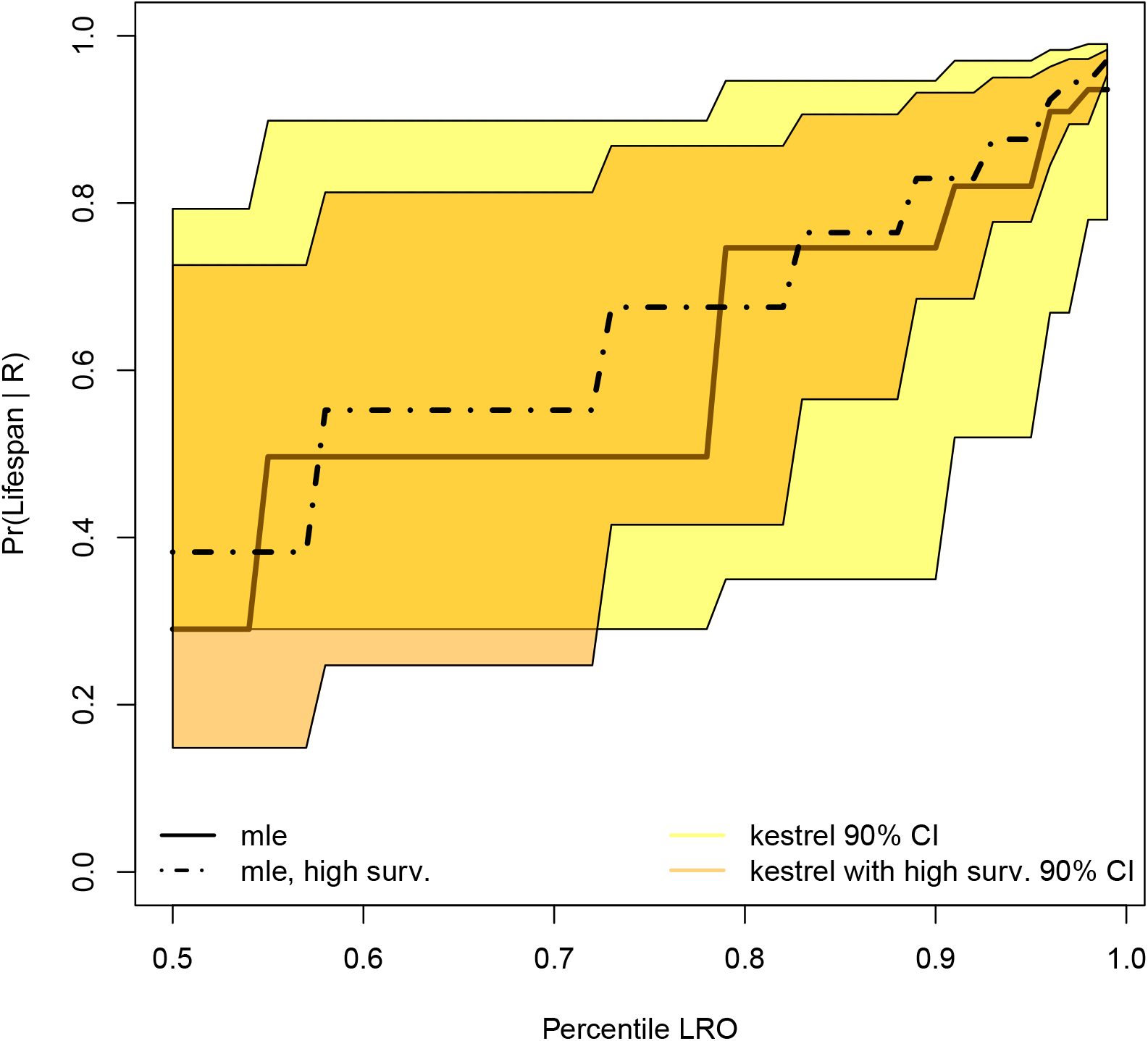
90% central intervals for *Pr*(Lifespan | *R*) for kestrels with a normal lifespan (yellow, mean lifespan 2.4) and kestrels with 20% higher survival in all stages (orange, mean lifespan 5.7). Figure generated by `compareKestrelLifespanCIVsLROPercentile2.R`, using data from `kestrelLifespanCIVsLROPercentile.R`, which gets data from `kestrelLifespanCondLRO.R`.

## Discussion

In previous work we used the sleight-of-hand of citing empirical evidence for reproductive skew as some of the motivation for theoretical papers about reproductive variance. In this paper we focused on the defining feature of reproductive skew: that a small number of individuals in a long right tail of the LRO distribution dominate reproduction. We use age-partitioning to identify the kinds of luck that contribute to existence of a fat right tail in the LRO distribution, and calculate the lifespan distribution conditional on LRO to examine what it takes to get into that tail.

In our case studies, we find that reproductive skew, like the variance, is driven largely by luck rather than trait variation. In fact, trait variation can actually decrease, rather than increase, the skewness in LRO. We suspect that trait variation will generally increase skewness when it reflects underlying variation in individual “quality” such that high-quality individuals have across-the-board better demographic performance, and may decrease skewness when it reflects life history trade-offs. In our case studies, the kittiwakes exhibit quality variation: high-quality birds have both higher survival and higher breeding probability. Variation among the fulmars comes from life history trade-offs. For example, individuals in group 2 are more likely to skip breeding and have lower juvenile survival than other groups, but breeders have higher survival.

Unlike reproductive variance, which can be dominated by luck in survival, growth, or environment depending on life history, reproductive skew is always dominated by luck in survival. Extreme reproductive success is mostly about living a long time. Luck in growth and environment and fecundity frequently contribute positively to the third central moment of LRO, *µ*_3_(*R*), but skewness is the third central moment scaled by the variance (skewness= *µ*_3_(*R*)/*Var*(*R*)^3/2^). The non-survival forms of luck contribute more to variance than to the third central moment, which decreases skewness.

Knowing that reproductive skew is dominated by luck in survival, we calculated the distribution of lifespan conditional on LRO. As we raise the threshold for success to higher and higher percentiles of the LRO distribution, we find that successful individuals have higher and higher lifespans, as expected. But how tightly lifespan is constrained by LRO depends on life history. In populations with large mean clutch size in good years, high LRO can result from getting a good year rather than living an extra few years, so high LRO is less tightly connected to long life. The link between exceptional LRO and exceptional lifespan was observed to be tighter in longer-lived species. However, this trend may just result from an often unrealistic property of the models we analyzed (and of typical matrix or integral projection models without explicit age-dependence): as individuals age, age-dependent average survival and fecundity become constant, without ever declining due to senescence. When average lifespan is high, the lucky few can then live very much longer than their less fortunate peers, and those extra-long lives are then the main cause of extreme success. We do not yet know if the lack of senescence in most stage-structured models is a serious problem. We have begun to consider examples where we have LRO data and a conventional senescence-free fitted matrix or integral projection model, and to compare predictions with observations.

Another important limitation of our analyses is that each individual is assumed to be an independent realization of the Markovian trip “from the cradle to the crypt” (Goodman, 1984) implied by the model’s survival, state transition, and clutch-size probability distributions. Variation in outcomes due to interactions between individuals — for example, the establishment and dynamics of social rankings and their implications — would only be included in the very rare case where social rank (either static or dynamic) is one of the attributes by which individuals are classified in the model. None of our case studies have this feature. For populations where social interactions are a major component of reproductive inequality, the methods here could serve as a “null model” analysis, quantifying the inequality that would develop even in the absence of social interaction effects.

Reproductive skew can have important implications for evolutionary dynamics, but we do not yet know as much as we would like. Tuljapurkar and Zuo (2023) found that the traditional approximation for the fixation probability of a rare, weakly beneficial allele in terms of the mean and variance of LRO can be very inaccurate when the allele-specific LRO distribution is highly skewed and bimodal. General theory has started to appear, but to our knowledge is limited to evolutionary models with extremely high reproductive skew, either infinite or tending to infinity along with population size in order to obtain analytic results. All these models lead to limiting coalescent processes in which multiple mergers can occur in a single generation, unlike the Kingman coalescent for Wright-Fisher models. One line of models, descended from Eldon and Wakeley (2006), assumes that each individual in each generation has a very small probability of producing a very large number of offspring (some fixed fraction *ψ* of the total population) and otherwise produces one offspring (for example, Der et al. (2012); Eldon and Stephan (2023)). The other, descended from Schweinsberg (2003), assumes that the offspring distribution has a power-law tail (with *p*(*u*) ∼ *u*^−(1+*α*)^,1 < *α* ≤ 2) implying infinite third moment (for example, Hallatschek (2018); Okada and Hallatschek (2021)). Constant population size in both cases is maintained by sampling *N* offspring at random to form the next generation.

For both offspring distributions, appropriately scaled forward-in-time allele dynamics in the large population limit are a diffusion process with occasional large jumps when the offspring of one individual make up a substantial fraction of the next generation. Eldon and Stephan (2023) found that the probability of slightly beneficial rare alleles fixing is reduced, but among alleles that do fix, the mean time to fixation is shortened. Our intuition for their finding is that the allele is likely to fix rapidly if an individual carrying the allele becomes one of the “lucky few” with exceptionally high LRO, and otherwise the allele is likely to be lost through drift. Hallatschek (2018); Okada and Hallatschek (2021) also found that extreme skew in LRO reduced the fixation probability of new beneficial mutations, relative to a Wright-Fisher model. However none of these studies, nor any others that we know of, is a “controlled experiment” comparing outcomes as LRO skewness is varied with the mean and variance held constant. We encourage population geneticists to further explore the effects of reproductive skew on evolutionary dynamics, with more realistic assumptions about the variance and skewness of the distribution of lifetime reproductive success. As a simple and possibly tractable approximation to the LRO variation in the models we have considered, we suggest offspring distributions with a substantial probability of zero offspring, while those that reproduce have a random number of offspring with strongly skewed distribution with finite but large variance and skew.

## Supporting information

Associated R code

## Acknowledgments

This research was supported by National Science Foundation grants DEB-1933497 (S.P.E.) and DEB-1933612 (R.E.S.). The Institute Paul Emile Victor (Programme IPEV 109), and Terres Australes et Antarctiques Françaises provided logistical and financial support for fulmar data collection by Barbraud and Jenouvrier. We thank Karen Abbott, Tom E.X. Miller and the Editor and anonymous reviewers for helpful comments.

## Supplementary Material

### S1 Computing contributions to skewness from trait value, birth state, and birth environment

In order to calculate the first few terms of the telescoping sum partition of skewness, we switch to an extended state space where the full extended state consists of the triple (*x*,*z*,*q*), trait *x ×* individual state *z ×* environment *q*. We also need to include several pre-birth states, so that we can condition on first *x*, then *x* and *z*_0_, then *x, z*_0_, and *q*_0_.

In this section we first describe the extended state space, and how to construct the necessary transition probability matrices on the extended state space. We then explain how those are used to compute the various age-dependent contributions to skewness for the original model. We use here notation for matrix models, assuming that an integral projection model has been discretized using a bin-to-bin method that results in an approximating large matrix model. Getting the matrices right is largely a matter of getting the indexing right when transferring information from the original state space matrices to the extended state space matrices, so our explanations are in the form of “pseudo-code” for how the matrices would be built in your language of choice. In what follows, we use `typewriter font` to denote code snippets, and we use the same variable names (e.g. `mz`, `bigmz`) that are used in our provided code. Thus, *ab* would denote the product of *a* with *b*, but `mz` is a single variable, and we use * to indicate a product of two variables in code, such as `mz * mq`.

If there are `mx` possible trait values, `mz` possible state values, and `mq` possible environments, define `bigmz = mz * mq`, the number of possible state-environment combinations. We then have the following groups of states that together comprise the necessary extended state space:

- *W*_1_: state 1: the single ur-state, ℵ. This represents an individual prior to birth, when no attributes have been assigned.
- *W*_2_: states 2 through (1 + `mx`), all possible trait values. This represents an individual prior to birth when only the trait value *x* has been assigned.
- *W*_3_: states 2 + `mx` through 1 + `mx` + `mx*mz`: all possible combinations of trait and state. This represents an individual prior to birth, when its trait and birth state have been assigned.
- *W*_4_: states 2 + `mx`+ `mx*mz` through 1 + `mx` + `mx*mz` + `mx*bigmz`: all possible combinations of trait, state, and environment. This represents living (post-birth) individuals, with all attributes assigned and with state and environment changing from one time-step to the next.

The transition matrix for the collection of all states then has the form:

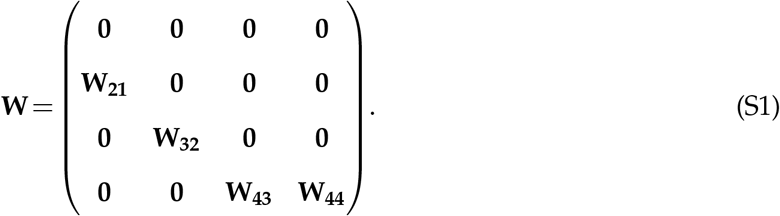

The submatrices are as follows:

- Going from ur-state to trait: **W**_**21**_ is a `mx` *×* 1 matrix that contains the trait distribution.
- Going from trait-only to traits and birth state:

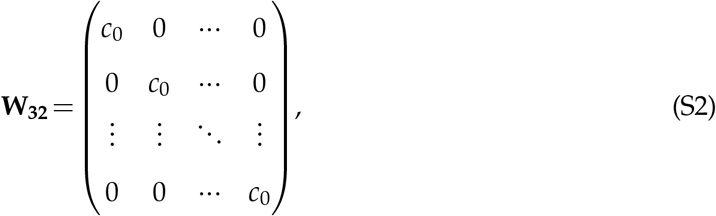

where *c*_0_ is a column vector with the birth state distribution. If the birth state distribution depends on trait value, then column *j* of **W**_**32**_ would be the birth state distribution for trait value *j*.

- Going from trait and birth state to trait, birth state, and birth environment:

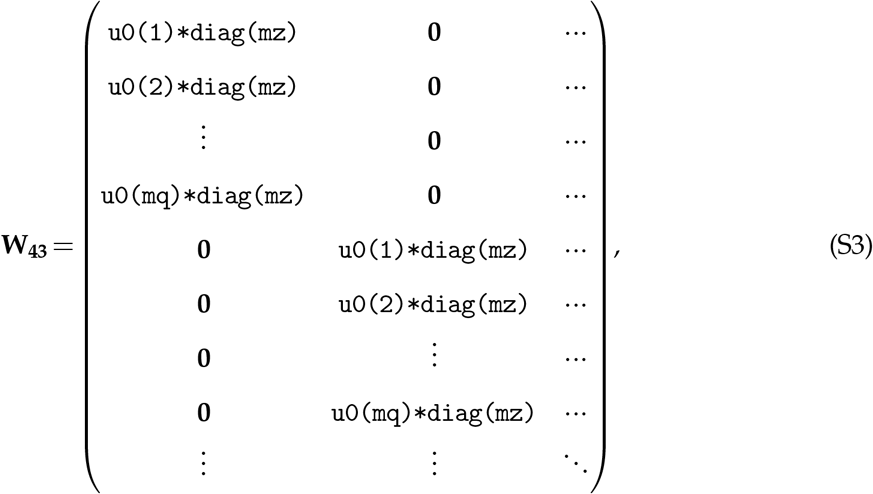

where *u*0(*j*) is the *j*th component of the stationary environment distribution, `diag(mz)` is an identity matrix of size `mz`, and we repeat the `u0(j)*diag(mz)` subblocks for as many trait values as we have.

- Going from trait, state, and environment to trait, state, and environment at ages 0,1,2,*…*:

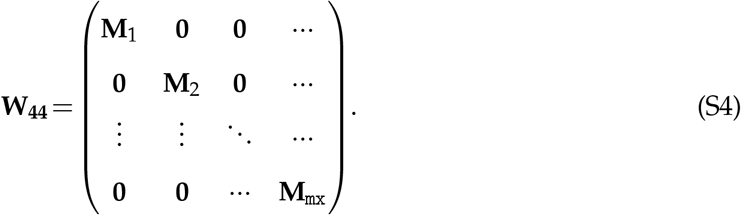

Our initial condition is the ur-state ℵ, which is a vector of the same length as the number of rows or columns of **W** with a 1 in the first component and zeros elsewhere. Multiplying ℵ by **W** gives us the pre-birth state with only trait assigned, multiplying by **W**^2^ gives us the pre-birth state with trait and birth state assigned, and multiplying by **W**^3^ gives us the pre-birth state with trait, birth state, and birth environment assigned.

**The contribution of traits to skewness** is the reduction in skewness from the unconditional skewness, 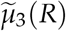, to the skewness conditional on trait, 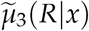:

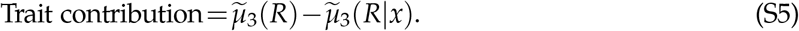

If there are multiple possible birth states and/or birth environments, the unconditional third moment of the pre-birth state, *µ*_3_(*R*), needs to be computed using the law of total cumulance. For the third moment, this is

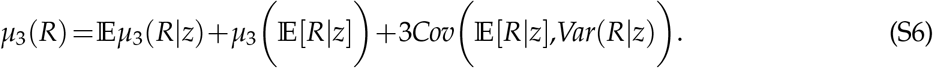

We then find the unconditional skewness by normalizing by the variance: 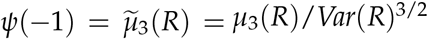.

To calculate the skewness conditional only on trait, we find find the skewness conditional on initial state and average over the pre-birth state with only trait assigned:

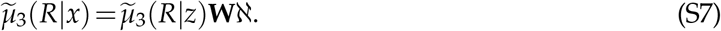

**The contribution of birth state to skewness** is the reduction in skewness from the skewness conditional on trait to the skewness conditional on trait and birth state:

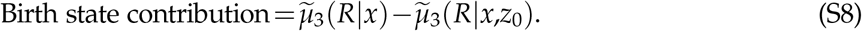

To calculate 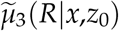 we average the skewness conditional on initial state over the pre-birth state with only trait and birth state assigned:

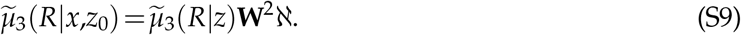

**The contribution of birth environment to skewness** is the reduction in skewness from the skewness conditional on trait and birth state to the skewness conditional on trait, birth state, and birth environment:

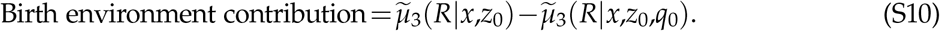

To calculate 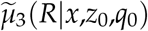, we average the skewness conditional on initial state over the pre-birth state with trait, birth state, and birth environment assigned:

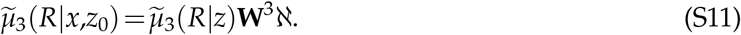

### S2 Models without environmental variation

In a model without environmental variation, the general formulas immediately above can be used with **Q**_*•*_ equal to the identity matrix, and *q* (and possibly *k*) not a component of the extended state state. Alternatively, the calculations can be done on a smaller state space with transition and reward matrices

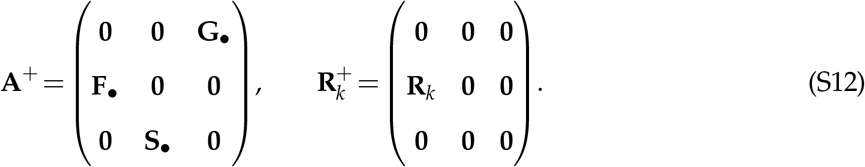

1. The expected skewness at the start of the time period is 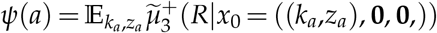, where the distribution of (*k*_*a*_,*z*_*a*_) is **A**^*a*^*m*_0_.
2. The expected skewness just after the fecundity update is 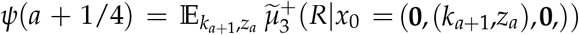, where the distribution of 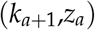 is **F**_*•*_**A**^*a*^*m*_0_.
3. The expected skewness just after the survival update is 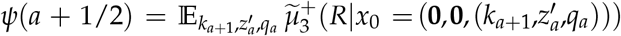, where the distribution of 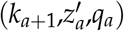 is **S**_*•*_**F**_*•*_**A**^*a*^*m*_0_.
4. The expected skewness just after the growth update is the expected skewness at the start of the next time period, *ψ*(*a*+1).

### S3 Post-breeding census models

In models based on a post-breeding census, reproduction occurs after survival and growth, so we need to change the extended state matrices and subsequent calculations in sections “Age-partitioning of skewness contributions from different luck types” and “Models without environmental variation.” For example, for a post-breeding case study with no environmental variation, the extended state transition matrix is

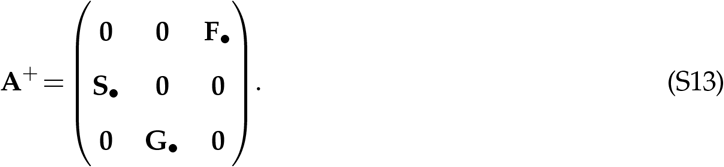

The survival, growth, and fecundity updates to skewness are as follows:

1. The expected skewness at the start of a time period is 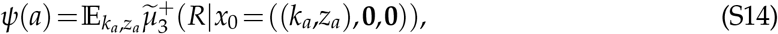 where the distribution of (*k*_*a*_,*z*_*a*_) is **A**^*a*^*m*_0_, where *m*_0_ is the initial state or state distribution.
2. The expected skewness just after the survival update is 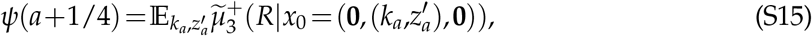 where the distribution of 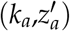 is **S**_*•*_**A**^*a*^*m*_0_.
3. The expected skewness just after the growth update is 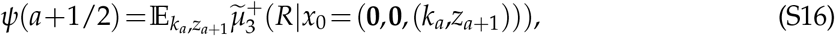 where the distribution of (*k*_*a*_,*z*_*a*+1_) is **G**_*•*_**S**_*•*_**A**^*a*^*m*_0_.
4. The expected skewness just after the fecundity update is the expected skewness at the start of the next time period, *ψ*(*a*+1).

### S4 Mean lifespan and central intervals for lifespan given LRO

Here we give a heuristic explanation for the general pattern that the width of the 90% Central Interval (CI) for lifespan conditional on LRO is wider for short-lived species. We say “heuristic” because we analyze a highly idealized approximation to what is actually observed in our models. In our models, age-dependent average survival and fecundity converge with age to age-independent constant values. Here we assume that survival and the probability distribution of fecundity are constant through an individual’s entire life, and moreover are independent from year to year.

We use *λ* to denote individual lifespan (age at death) considered as a random variable, and *R* to denote LRO considered as a random variable.

As a rough approximation to the situation where adult average vital rates are nearly independent of age, suppose that for all of their life an individual has the same probability distribution for annual clutch size *f*_*a*_, with mean 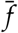 and variance 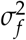. Then

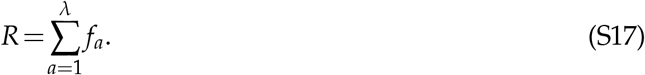

Invoking the Central Limit Theorem (and ignoring any temporal correlations in annual fecundity), the conditional distribution of *R* given *λ* can be approximated by a Normal distribution:

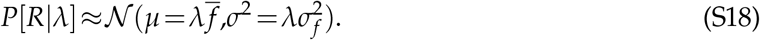

In this approximation, *R* is now a continuous random variable, not discrete. The approximation holds in the same sense that (for example) a Poisson distribution with large mean is approximated by a Gaussian distribution with the same mean and variance.

Let *φ*(*z*; *µ, σ*^2^) denote a Normal probability density function with mean *µ*, variance *σ*^2^. The joint distribution for *R* and *λ* is then given by

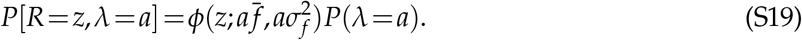

Then

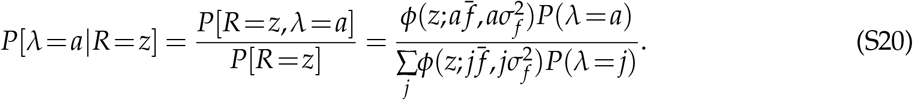

We want to ask how the right-hand size varies with *a*, because we are interested in the width of the Conditional Interval for lifespan *λ*. The denominator is independent of *a*. In the numerator, suppose that *σ*_*f*_ is “not too large”, and the variance of *λ* is “not too small” in the sense that *P*(*λ* = *a*) does not vary greatly across the range of likely *a* values. Then the fastest-changing component in the numerator (as *a* is varied) will be the term

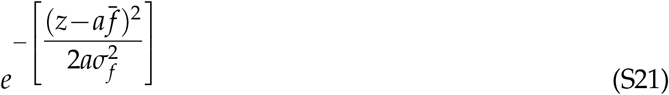

coming from the Gaussian density in the numerator. This term comes from a Gaussian where 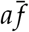 is the mean, and *z* is a possible value. But let’s turn that around, and think of it as coming from a Gaussian where *z* is the mean, 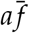 is a possible value, and the standard deviation of the distribution is 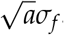.

In particular, the value of (S21) will be small whenever 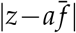 is more than 2 or 3 times 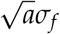. Note, to think in that way, we have to suppose that *σ*_*f*_ is small enough that the value of (S21) falls off so rapidly as 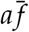 departs from the “mean” *z*, that the denominator 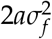 can be regarded as constant over the range where (S21) is not minutely small. Let us do that, and see what it gets us; we will then come back to ask what “small enough” means.

Coming back to (S20), the behavior of (S21) as a function of *a* says that the likely values of lifespan *a*, conditional on LRO *R* = *z*, satisfy

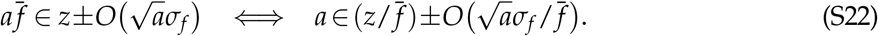

The most likely lifespan given *R* = *z* is approximately 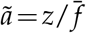, and (S22) says that likely values *a* for the lifespan cannot be too far from *ã*: with high probability, the conditional lifespan *a* is close to the most likely lifespan. We can therefore re-write the right-hand expression in (S22) as

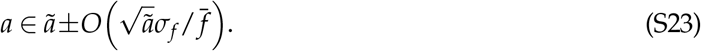

Eqn. (S22) therefore can be interpreted as follows: in the probability distribution of lifespan *λ* conditional on a particular value of LRO *R*, the width of the range of likely values (and thus the CI width on absolute scale) is approximately proportional to the square-root of the most likely lifespan given *R*.

Figure S1AB shows a simulated example, in which 80% of newborns survive to age 1, lifespan conditional on survival to age 1 has a geometric distribution with survival probability 0.9 or 0.95, and starting at age 2 adults breed each year and produce a Poisson number of new recruits with mean 0.5. Mean lifespan is 8.02 in panel A), and 16.03 in panel B) — these numbers vary slightly from run to run. The vertical lines are drawn at the 95th percentiles of LRO in each panel. Note that the axis scales in panels A) and B) are the same, based on the range of values in B). The relationship between lifespan and LRO is nearly the same in the two panels, because the number of offspring per year of breeding is the same. The two-fold difference in mean lifespan between panels A) and B) therefore produces a roughly two-fold difference in the 95th percentile of LRO, and a roughly two-fold difference in the typical lifespan conditional on LRO being at the 95th percentile (median lifespan 52 versus 25). The 90% central CI for lifespan conditional on LRO being at its 95th percentile (shown by the dashed red horizontal lines) is wider in panel B) by a factor of 1.53, close to the value 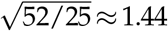 predicted theoretically based on the ratio of median lifespans, because 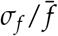 is the same in A) and B).

**Figure S1:**
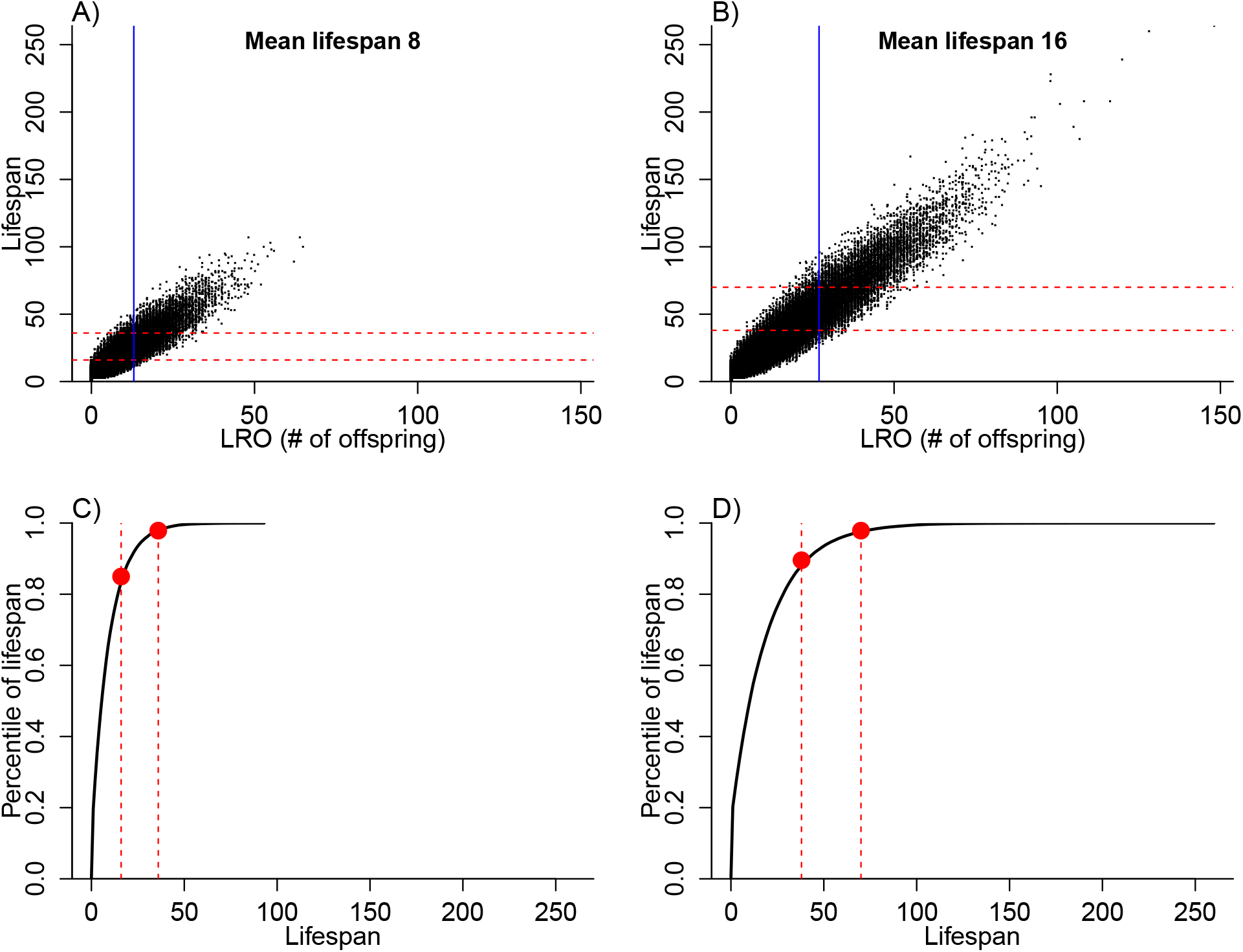
**A), B)**: Simulated joint distribution of LRO and lifespan, with two different values of annual adult survival probability producing a two-fold difference in mean lifespan. Values for 250,000 simulated individuals are plotted for each survivorship. In both cases, 80% of individuals survive to age 1, and those that do survive then have “adult” lifespan with a geometric distribution on the positive integers. Breeding begins at age 2 and continues annually until death, with Poisson(0.5) annual per-capita offspring production. The sold blue vertical lines are drawn at the 95th percentile of LRO. The dashed red horizontal lines are the boundaries of the 90% central interval for lifespan conditional on LRO being at its 95th percentile. In panels **C)** and**D)**, the 90% central intervals for conditional lifespan (dashed red lines) are translated from lifespan (*x*-axis) to percentiles of lifespan (*y*-axis), based on the relationships between lifespan and percentile of lifespan for the two different values of adult survival (solid black curves). Figure made by script `survival effect.R`.

Finally, we need to ask what the central CI becomes when we express it in terms of percentiles of lifespan, rather than lifespan. This is illustrated in panels C) and D), which correspond to A) and B) respectively. The dashed vertical red lines, in each panel, are drawn at the upper and lower endpoints of the 90% central CI for lifespan, conditional on LRO being at its 95th percentile. The black curve in each panel shows percentile of lifespan as a function of lifespan. The vertical locations of the two solid red circles in each panel therefore mark the upper and lower endpoints of the central CI on the percentile scale. With longer lifespan (panel D versus panel C), the central CI is wider in terms of lifespan (i.e., the dashed red lines are are further apart horizontally), but narrower in terms of percentiles of lifespan (i.e, the solid red circles dots are closer together vertically). So on the percentile scale, it is the shorter lifespan that results in the wider conditional CI for lifespan. As noted above, doubling the lifespan increases the CI width on lifespan scale by a factor of 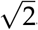. However, the relationship between percentile of lifespan and absolute lifespan becomes less steep (panel D versus panel C), by a factor of roughly 2 — i.e., to get the same range of percentile lifespans in panel D, we need to double the range of absolute lifespans. This is because doubling the mean lifespan means stretching the approximately geometric distribution by a factor of 2. As a result, the CI width on the percentile of lifespan scale is smaller by a factor of 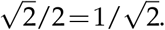.

So in general, the implication of (S23) is that if mean lifespan is increased by a factor *c*> 1, all else being equal, the CI width on percentile of lifespan scale goes down by a factor of 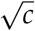. Longer mean lifespan therefore gives narrower 90% CIs on the percentile of lifespan scale, as we observe in our cases studies, all else being equal.

Intuitively, our final conclusion is driven by the fact that in (S18), the conditional mean of LRO given lifespan *λ* grows in proportion to *λ*, while the conditional standard deviation grows as 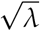.^6^ So long as that is true, and the between-year correlation in fecundity (resulting from between-year correlation of individual state) is weak enough for the Central Limit Theory to apply, eqn. (S18) will provide a good approximation, leading to our final conclusion.

Now let us return to the assumption, made in deriving (S22), that *σ*_*f*_ is “small enough”. We can write (S21) as

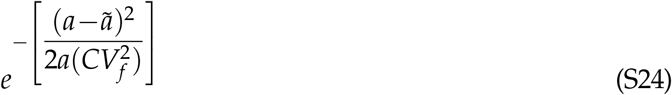

Where 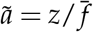 is approximately the most likely conditional lifespan, and *CV*_*f*_ is the coefficient of variation of *f*. “Small enough” means that (S24) can be approximated by

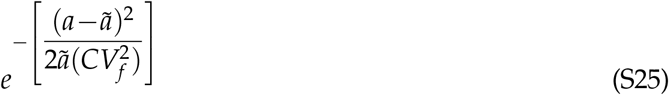

as a function of *a*, in the very weak sense that the two functions have approximately equal-size regions where they are not negligibly small. Numerically, we find this approximation improves with larger values of *ã*and smaller values of *CV*_*f*_, and it is accurate enough for our purposes so long as *ã*≥ 5 and *CV*_*f*_ ≤ 1.

## S5 Supplemental figures

**Figure S2:**
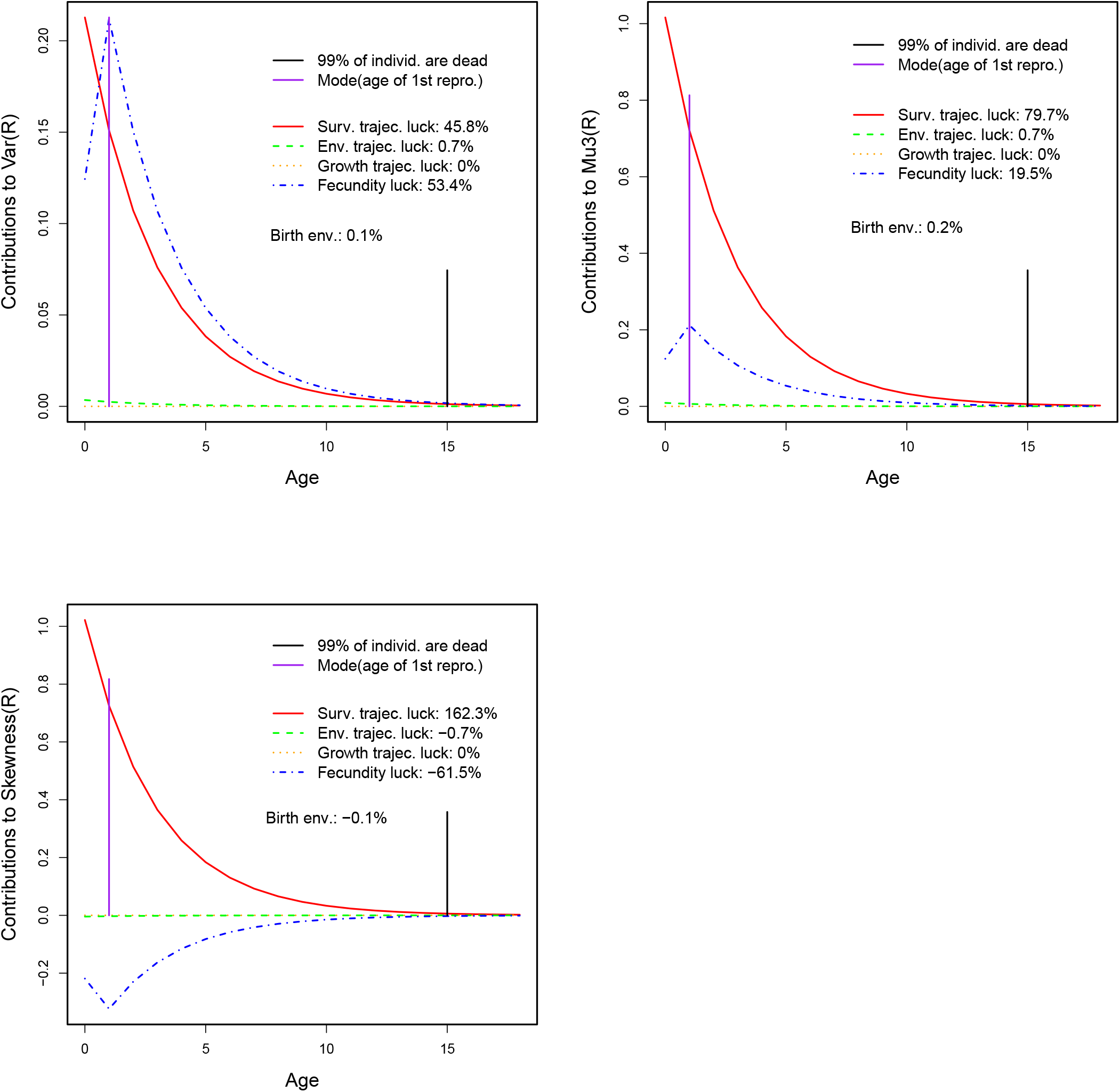
Variance, third central moment, and skewness of lifetime reproductive output for lesser kestrels. Lines show the contributions of different forms of luck as a function of age. Top left: Partition of variance. Top right: Partition of third central moment. Bottom: Partition of skewness. The percent contributions are calculated with respect to total skewness, not the sum of the absolute magnitudes of contributions. Generated by `kestrelVarMu3Partition.R` and `kestrelVarSkewnessPartition.R`.

**Figure S3:**
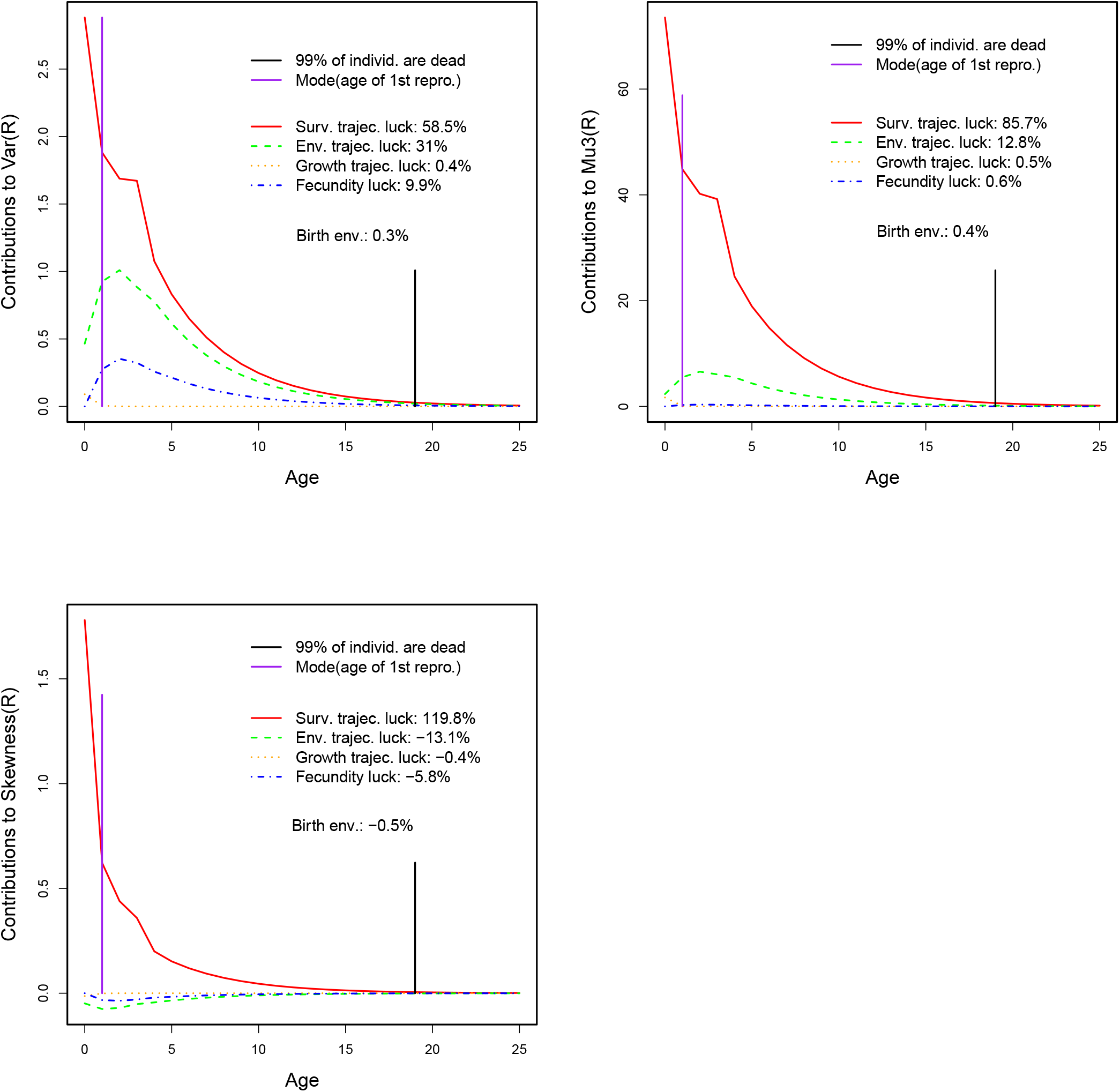
Variance, third central moment, and skewness of lifetime reproductive output for *Umbonium costatum*. Lines show the contributions of different forms of luck as a function of age. Top left: Partition of variance. Top right: Partition of third central moment. Bottom: Partition of skewness. The percent contributions are calculated with respect to total skewness, not the sum of the absolute magnitudes of contributions. Generated by `umboniumVarMu3Partition.R` and `umboniumVarSkewnessPartition.R`.

**Figure S4:**
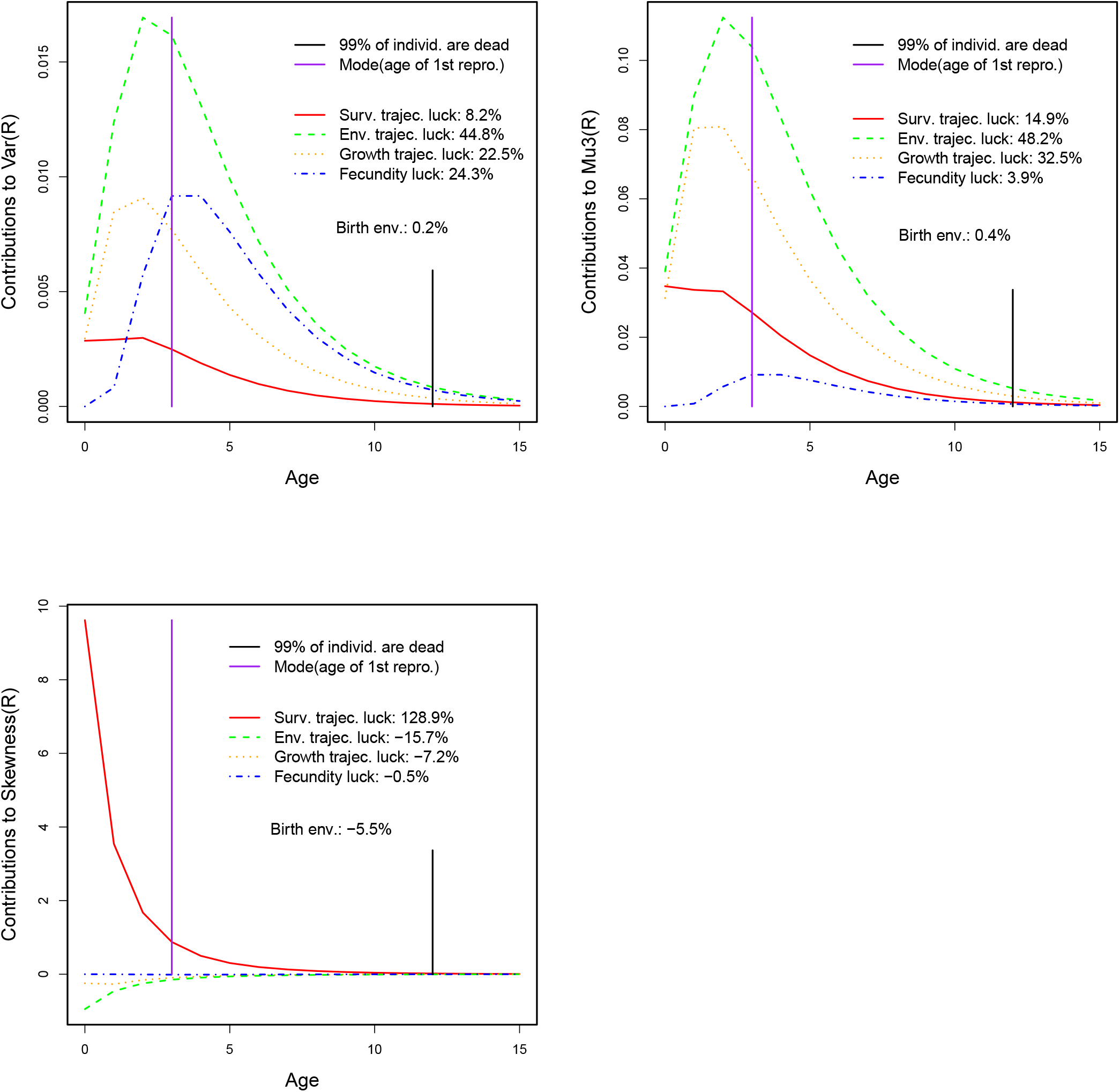
Variance, third central moment, and skewness of lifetime reproductive output for *Lomatium bradshawii*. Top left: Partition of variance. Lines show the contributions of different forms of luck as a function of age. Top right: Partition of third central moment. Bottom: Partition of skewness. The percent contributions are calculated with respect to total skewness, not the sum of the absolute magnitudes of contributions. Generated by `lomatiumVarMu3Partition.R` and `lomatiumSkewnessPartition5.R`.

**Figure S5:**
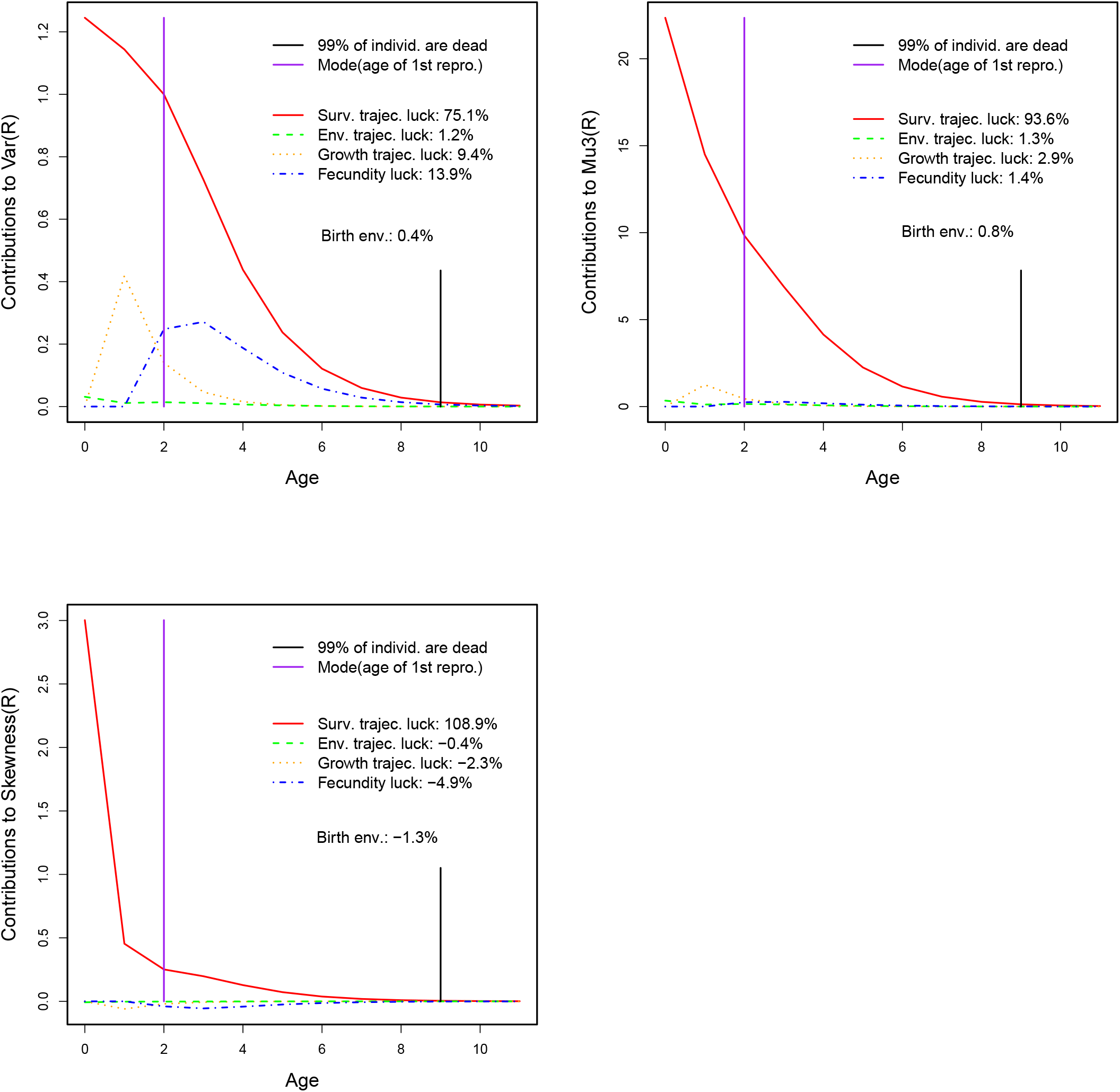
Variance, third central moment, and skewness of lifetime reproductive output for *Xenosaurus platyceps*. Lines show the contributions of different forms of luck as a function of age. Top left: Partition of variance. Top right: Partition of third central moment. Bottom: Partition of skewness. The percent contributions are calculated with respect to total skewness, not the sum of the absolute magnitudes of contributions. Generated by `xenosaurusVarMu3Partition.R` and `xenosaurusVarSkewnessPartition.R`.

**Figure S6:**
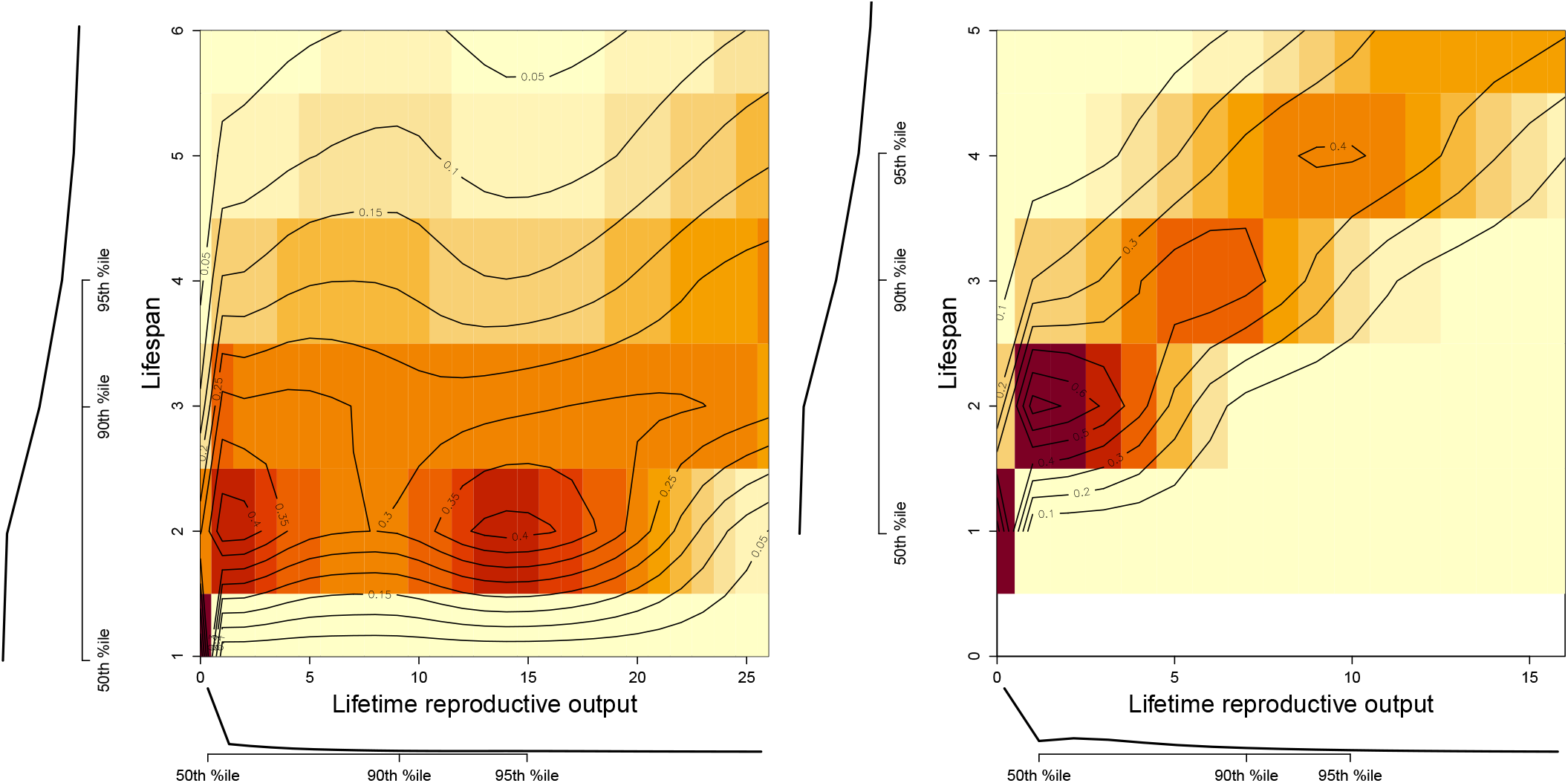
*Pr*(Lifespan | *R*) for sculpins with environmental variation in fecundity (left) and without (fecundity in each stage is set to its environmental average) (bottom right). Generated by `plotSculpinLifespanCondLRO3.R`, which gets data from `sculpinLifespanCondLRO.R`.

## Footnotes

1 For example, T. Shaw, *The oldest known common loons find success at Seney National Wildlife Refuge*, www.fws.gov/story/oldest-known-common-loons, accessed Dec. 7, 2023.

2 Note that what we have called luck and traits, Caswell and collaborators have called individual stochasticity and individual

3 We use “clutch size” to mean the number of offspring produced by a single female in one year (or one time step of the original model), regardless of taxon.

4 The distribution of offspring number can become correlated with future states in subtle ways. See the section “Where these calculations break down and what to do about it” in Snyder and Ellner (2022) for details.

5 The subscript dots are ways of distinguishing the survival, growth, etc. matrices that work on the extended state space from those that work in the original state space.

6 This is why (S21) has 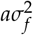 as the variance of the Gaussian, not 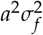.

