## Supplementary material for "To prosper, live long: understanding the sources of reproductive skew and extreme reproductive success in structured populations": Associated R code: compareKestrelLifespanCIVsLROPercentile2.R

```
##########################################################################
##
## plotLifespanCIVsLROPercentile.R: Plots the 90% confidence interval
## of lifespan conditional on LRO.  Both LRO and lifespan are
## expressed as percentiles so that we can compare across species.
##
## kestrelCILifespan.dat and kestrelCILifespanHighSurv.dat are
## generated by kestrelLifespanCIVsLROPercentile.R.
##
##########################################################################

rm(list=ls(all=TRUE)); graphics.off();
##require ("viridisLite")
##require ("colorspace")
require ("RColorBrewer")

colorVec = c(adjustcolor("yellow", alpha.f=0.5),
             adjustcolor("orange", alpha.f=0.5),
             adjustcolor("purple", alpha.f=0.5))

dKestrel = read.table ("kestrelCILifespan.dat", header=T)
dKestrelHighSurv = read.table ("kestrelCILifespanHighSurv.dat", header=T)

ymax = max (dKestrel$top5, dKestrelHighSurv$top5)

plot (dKestrel$Rpercentiles, dKestrel$bottom5, type="l",
      ylim=c(0, 1), col=colorVec[1], lty=1,
      xlab="Percentile LRO", ylab="Pr(Lifespan | R)")
lines (dKestrel$Rpercentiles, dKestrel$top5, type="l",
       col=colorVec[1], lty=2)
polygon (c(dKestrel$Rpercentiles, rev(dKestrel$Rpercentiles)),
         c(dKestrel$bottom5, rev(dKestrel$top5)),
         col=colorVec[1])
lines (dKestrel$Rpercentiles, dKestrel$mle, type="l",
       lty=1, lwd=3)

lines (dKestrelHighSurv$Rpercentiles, dKestrelHighSurv$bottom5, type="l",
       col=colorVec[2], lty=1)
lines (dKestrelHighSurv$Rpercentiles, dKestrelHighSurv$top5, type="l",
       col=colorVec[2], lty=2)
polygon (c(dKestrelHighSurv$Rpercentiles, rev(dKestrelHighSurv$Rpercentiles)),
         c(dKestrelHighSurv$bottom5, rev(dKestrelHighSurv$top5)),
         col=colorVec[2])
lines (dKestrelHighSurv$Rpercentiles, dKestrelHighSurv$mle, type="l",
       lty=4, lwd=3)

legend (x="bottomright",
##        legend=c("kestrel 90% CI (mean lifespan 2.4)",
##                 "kestrel with high surv. 90% CI (mean lifespan 5.7)"),
        legend=c("kestrel 90% CI",
                 "kestrel with high surv. 90% CI"),
        col=colorVec, lwd=2, bty="n")

legend (x="bottomleft",
        legend=c("mle", "mle, high surv."), lty=c(1,4), lwd=2, bty="n")

##dev.copy2pdf (file="compareKestrelLifespanCIVsLROPercentile2.pdf")
```
