## Supplementary material for "To prosper, live long: understanding the sources of reproductive skew and extreme reproductive success in structured populations": Associated R code: compareUmboniumLifespanCIVsLROPercentile.R

```
##########################################################################
##
## plotLifespanCIVsLROPercentile.R: Plots the 90% confidence interval
## of lifespan conditional on LRO.  Both LRO and lifespan are
## expressed as percentiles so that we can compare across species.
##
## umboniumCILifespan.dat and umboniumCILifespanNoEnvVarInFec.dat are
## generated by unboniumLifespanCIVsLROPercentile.R.
##
##########################################################################

rm(list=ls(all=TRUE)); graphics.off();
##require ("viridisLite")
##require ("colorspace")
require ("RColorBrewer")

numSpecies = 3
##colorVec = brewer.pal (numSpecies, "RdPu")
colorVec = c(adjustcolor("yellow", alpha.f=0.5),
             adjustcolor("orange", alpha.f=0.5),
             adjustcolor("purple", alpha.f=0.5))

dUmbonium = read.table ("umboniumCILifespan.dat", header=T)
dUmboniumNoEnvVarInFec = read.table ("umboniumCILifespanNoEnvVarInFec.dat", header=T)

ymax = max (dUmbonium$top5, dUmboniumNoEnvVarInFec$top5)

par (cex.lab=1.4)
plot (dUmbonium$Rpercentiles, dUmbonium$bottom5, type="l",
      ylim=c(0, 1), col=colorVec[1], lty=1,
      xlab="Percentile LRO", ylab="Percentile lifespan given R")
lines (dUmbonium$Rpercentiles, dUmbonium$top5, type="l",
       col=colorVec[1], lty=2)
polygon (c(dUmbonium$Rpercentiles, rev(dUmbonium$Rpercentiles)),
         c(dUmbonium$bottom5, rev(dUmbonium$top5)),
         col=colorVec[1])
lines (dUmbonium$Rpercentiles, dUmbonium$mle, type="l",
       lty=2, lwd=3)

lines (dUmboniumNoEnvVarInFec$Rpercentiles, dUmboniumNoEnvVarInFec$bottom5, type="l",
       col=colorVec[2], lty=1)
lines (dUmboniumNoEnvVarInFec$Rpercentiles, dUmboniumNoEnvVarInFec$top5, type="l",
       col=colorVec[2], lty=2)
polygon (c(dUmboniumNoEnvVarInFec$Rpercentiles, rev(dUmboniumNoEnvVarInFec$Rpercentiles)),
         c(dUmboniumNoEnvVarInFec$bottom5, rev(dUmboniumNoEnvVarInFec$top5)),
         col=colorVec[2])
lines (dUmboniumNoEnvVarInFec$Rpercentiles, dUmboniumNoEnvVarInFec$mle, type="l",
       lty=1, lwd=3)

legend (x="bottomright",
        legend=c("90% central interval",
                 "90% CI, no env. var. in fec."),
        col=colorVec, lwd=2, bty="n")

legend (x="bottomleft",
        legend=c("Most prob.", "Most prob., no env. var. in fec."), lty=c(1,4), lwd=2, bty="n")

##dev.copy2pdf (file="compareUmboniumLifespanCIVsLROPercentile.pdf")
```
