## Supplementary material for "To prosper, live long: understanding the sources of reproductive skew and extreme reproductive success in structured populations": Associated R code: fulmarVarSkewnessPartition.R

```
#############################################################################
##
## fulmarVarSkewnessPartition: Age-partitions luck and env. var. for
## fulmars.  Assume that the three sea ice conditions occur with equal
## frequency and no temporal autocorrelation for now.  Uses
## genericPartitionVarSkewness6.R, our now-canonical partitioning
## code.
##
## Some general definitions:
##
## mz: number of stages/sizes
##
## numEnv: number of environments/year types
##
## P: survival/growth transition matrix
##
## c0: initial distribution of sizes/stages
##
## F: fecundity matrix
##
## Q: environment transition matrix
## 
## M: "megamatrix" governing size x environment transitions
##
## m0: initial distribution of size x environment
##
#############################################################################

rm(list=ls(all=TRUE)); graphics.off(); 
require(Matrix); require(statmod);

demogDir = "./fulmarMatrices/"
if(.Platform$OS.type=="windows") {
    demogDir = "c:/repos/fanova/envVar/fulmarMatrices"
    codeDir = "c:/repos/fanova/envVar" 
}

source ("Utilities.R")
source ("partitioningUtilities.R")
source ("genericPartitionVarSkewness6.R")
source ("megamatrixFunctions.R")

lineTypes = c(1,2,3,4)
names(lineTypes) = c("surv", "env", "growth", "fec")

colors = c("red", "green", "orange", "blue")
names(colors) = c("surv", "env", "growth", "fec")

debugging = TRUE

noEnvVar = FALSE  ## Turn off environmental variation for debugging
equallyFrequentEnv = FALSE  ## each env. occurs 1/3 of the time
stickyEnv = FALSE ## positively autocorrelated env.
if (noEnvVar)  ## If no env. var., this is the one env. we have.
  fixedEnv = 2

noTraitVar = FALSE  ## everyone is born into group fixedTrait
if (noTraitVar)
  fixedTrait = 2

ourFecundityDef = TRUE  # successful breeders have 1 chick, everyone else
                        # has zero

mz = 4;   # size of the age/stage transition matrix
numTraits = 3  # number of groups
numEnv = 3  # number of environmental levels
bigmz = numEnv*mz

### really maxAge+1, because we let age start at 0 
##maxAge = 500
maxAge = 1000

## Set up birth size distribution --- all are born as pre-breeders
c0 = c(1, rep(0, mz-1));

## "mixing distribution" (group distribution) from eq. 3, Jenouvrier
## et al., JAE 2017

if (noTraitVar) {
  groupDist = c(0, 0, 0)
  groupDist[fixedTrait] = 1
} else {
  groupDist = c(0.14, 0.67, 0.19)
}

## Read in demographic matrices.
setwd (demogDir)

Parray = Farray = array (0, dim=c(numTraits, numEnv, mz, mz))
Slist = Glist = list (numEnv)

## Low sea ice
if (ourFecundityDef) {
  Farray[1,1,1,2] = Farray[2,1,1,2] = Farray[3,1,1,2] = 1
} else {
  Farray[1,1,,] = as.matrix(read.table("F.lowSICgp1.txt"))
  Farray[2,1,,] = as.matrix(read.table("F.lowSICgp2.txt"))
  Farray[3,1,,] = as.matrix(read.table("F.lowSICgp3.txt"))
}
Parray[1,1,,] = as.matrix(read.table("U.lowSICgp1.txt"))
Parray[2,1,,] = as.matrix(read.table("U.lowSICgp2.txt"))
Parray[3,1,,] = as.matrix(read.table("U.lowSICgp3.txt"))

## ordinary sea ice
if (ourFecundityDef) {
  Farray[1,2,1,2] = Farray[2,2,1,2] = Farray[3,2,1,2] = 1
} else {
  Farray[1,2,,] = as.matrix(read.table("F.ordSICgp1.txt"))
  Farray[2,2,,] = as.matrix(read.table("F.ordSICgp2.txt"))
  Farray[3,2,,] = as.matrix(read.table("F.ordSICgp3.txt"))
}
Parray[1,2,,] = as.matrix(read.table("U.ordSICgp1.txt"))
Parray[2,2,,] = as.matrix(read.table("U.ordSICgp2.txt"))
Parray[3,2,,] = as.matrix(read.table("U.ordSICgp3.txt"))

## High sea ice
if (ourFecundityDef) {
  Farray[1,3,1,2] = Farray[2,3,1,2] = Farray[3,3,1,2] = 1
} else {
  Farray[1,3,,] = as.matrix(read.table("F.highSICgp1.txt"))
  Farray[2,3,,] = as.matrix(read.table("F.highSICgp2.txt"))
  Farray[3,3,,] = as.matrix(read.table("F.highSICgp3.txt"))
}
Parray[1,3,,] = as.matrix(read.table("U.highSICgp1.txt"))
Parray[2,3,,] = as.matrix(read.table("U.highSICgp2.txt"))
Parray[3,3,,] = as.matrix(read.table("U.highSICgp3.txt"))

setwd ("../")

## Sea ice transition matrix.  Currently we assume that the three sea
## ice conditions occur with equal frequency and no temporal
## autocorrelation.
if (noEnvVar) {
  Q = diag(3)
} else if (equallyFrequentEnv) {
  Q = matrix (1/3, 3, 3)
} else {
  Q = matrix (rep(c(0.1, 0.8, 0.1), 3), 3, 3)
}

if (stickyEnv) {s = 0.5; Q = diag(s, numEnv) + (1 - s)*Q}

## Initial env. distribution = stationary environment distribution
if (noEnvVar) {
  u0 = rep(0, numEnv)
  u0[fixedEnv] = 1
} else {
  u0=domEig(Q)$w; 
}

## The initial cross-classified distribution
m0 = matrix (outer (c0, as.vector(u0)), bigmz, 1)

## Allocate some memory
survTrajecSkewnessCondX = growthTrajecSkewnessCondX = envTrajecSkewnessCondX =
  fecSkewnessCondX = matrix (0, numTraits, maxAge)
surv = fecUpdateSkewness = survUpdateSkewness =
  growthUpdateSkewness = envUpdateSkewness = numeric (maxAge)
reproArrivalMode = lifespan = birthEnvSkewnessCondX =
  birthEnvVarCondX = numeric (numTraits)
expRCondX = totVarCondX = totSkewnessCondX = totMu3CondX =
  numeric (numTraits)

survTrajecVarCondX = growthTrajecVarCondX = envTrajecVarCondX = fecVarCondX =
  matrix (0, numTraits, maxAge) 
survUpdateVar = growthUpdateVar = envUpdateVar = fecUpdateVar =
  numeric (maxAge)
##urVarCondTrait = birthStateVarCondX = birthEnvVarCondX = numeric
##(numTraits)


## start set-up for reward matrices.  We need to specify these and
## pass them in, since clutch sizes are not Poisson-distributed.
esbigmz = 4*bigmz
bsbigmz = 1 + mz + bigmz

esF = esR1 = esR2 = esR3 = matrix (0, esbigmz+1, esbigmz+1)
bsF = bsR1 = bsR2 = bsR3 = matrix (0, bsbigmz+1, bsbigmz+1)

## Define M.  Our state is breeding class x sea ice level, with one
## matrix per trait value.
Marray = array (NA, dim=c(numTraits, bigmz, bigmz))
for (x in 1:numTraits) {
  for (i in 1:numEnv) {
    for (j in 1:numEnv) {
      Marray[x, (i-1)*mz + 1:mz, (j-1)*mz + 1:mz] = Parray[x,j,,]*Q[i,j]
    }
  }
}

bigFarray = array (0, dim=c(numTraits, bigmz, bigmz))
for (x in 1:numTraits) {
  for (i in 1:numEnv) {
    for (j in 1:numEnv) {
      ## Fecundity depends on the parent's environment, j, but the
      ## chick may be born into any of the three enviroments.
      bigFarray[x, (i-1)*mz + 1:mz, (j-1)*mz + 1:mz] =
        Farray[x,j,,] * Q[i,j]
    }
  }
}

## Loop over traits
for (x in 1:numTraits) {

  Plist = Flist = list ()
  for (q in 1:numEnv) {
    Plist[[q]] = Parray[x,q,,]
    Flist[[q]] = Farray[x,q,,]
  }

  ## define reward matrices.  F doesn't actually depend on x, but I'm
  ## playing this by the book.
  F = bigFarray[x,,]
  esF[bigmz + 1:bigmz, 1:bigmz] = F
  bsF[1, 1 + mz + 1:bigmz] = colSums (F)

  for (j in 1:esbigmz) 
    esR1[,j] = sum(esF[,j])
  for (j in 1:bsbigmz) 
    bsR1[,j] = sum(bsF[,j])

  esR2 = esR1^2
  esR3 = esR1^3
  bsR2 = bsR1^2
  bsR3 = bsR1^3  

  out = getVarSkewnessPartitionsEnvVar (Plist, Flist, Q, m0, maxAge,
                                        esR1, esR2, esR3,
                                        bsR1, bsR2, bsR3)

  birthEnvSkewnessCondX[x] = out$birthEnvSkewness
  survTrajecSkewnessCondX[x,] = out$survTrajecSkewness
  growthTrajecSkewnessCondX[x,] = out$growthTrajecSkewness
  envTrajecSkewnessCondX[x,] = out$envTrajecSkewness
  fecSkewnessCondX[x,] = out$fecSkewness
  ## totSkewnessCondX matches the equivalent in the old code
  totSkewnessCondX[x] = out$totSkewness
  lifespan[x] = out$lifespan

  birthEnvVarCondX[x] = out$birthEnvVar
  survTrajecVarCondX[x,] = out$survTrajecVar
  growthTrajecVarCondX[x,] = out$growthTrajecVar
  envTrajecVarCondX[x,] = out$envTrajecVar
  fecVarCondX[x,] = out$fecVar
  ## totVarCondX matches urVarCondTrait in the old code!
  totVarCondX[x] = out$totVar
  totVarCondX2 = birthEnvVarCondX[x] +
    sum(survTrajecVarCondX[x,] + growthTrajecVarCondX[x,] +
        envTrajecVarCondX[x,] + fecVarCondX[x,])

  totMu3CondX[x] = totSkewnessCondX[x]*totVarCondX[x]^1.5

  ## Get expected LRO while you're at it.

  ## Define megamatrices M and F
  F = M = matrix (0, bigmz, bigmz)
  for (i in 1:numEnv) {
    for (j in 1:numEnv) {
      M[(i-1)*mz + 1:mz, (j-1)*mz + 1:mz] = Plist[[j]]*Q[i,j]
      F[(i-1)*mz + 1:mz, (j-1)*mz + 1:mz] = Flist[[j]]*Q[i,j]
    }
  }
  N = solve (diag(bigmz) - M)
  expRCondX[x] = (colSums (F %*% N)) %*% m0

  ## And get modal time to reproductive state
  absorbingStates = (0:2)*mz + 2

  out = getModalTimeToHitState (absorbingStates, Plist, Flist,
                                Q, m0, maxAge)
  reproArrivalMode[x] = out$hitTimeMode

  ## Sanity check on variance. 
  if (debugging) {
    expRCondXZ = colSums (F %*% N)
    pb = Farray[x,1,1,]
    b = rep(1, bigmz) ## one chick per pair if they do reproduce, but that's
    ## accounted for in pb
    for (q in 2:numEnv)
      pb = c(pb, Farray[x,q,1,])
    
    sigbsq = pb*(1 - pb)
    rbarPib = expRCondXZ %*% M     # \bar{r} \pi_b
    r2 = (sigbsq + (pb*b)^2 + 2*pb*b*rbarPib) %*% N
    varRCondXZ = r2 - expRCondXZ^2
    expZVarRCondXZ = varRCondXZ %*% m0
    varZExpRCondXZ = sum(m0*expRCondXZ^2) - sum(m0*expRCondXZ)^2
    cat ("Var. check:", totVarCondX[x], "should = ",
         expZVarRCondXZ + varZExpRCondXZ, 
         "should = ", totVarCondX2, "\n")
  }
}

traitAve = function (x) {
  return (sum(groupDist*x))
}

traitVar = function (x) {
  return (sum(groupDist*x^2) - traitAve(x)^2)
}

traitMu3 = function (x) {
  return (sum(groupDist*(x - traitAve(x))^3))
}

## Average over traits
birthEnvSkewness = traitAve (birthEnvSkewnessCondX)
fecSkewness = apply (fecSkewnessCondX, 2, traitAve)
survTrajecSkewness = apply (survTrajecSkewnessCondX, 2, traitAve)
growthTrajecSkewness = apply (growthTrajecSkewnessCondX, 2, traitAve)
envTrajecSkewness = apply (envTrajecSkewnessCondX, 2, traitAve)

birthEnvVar = traitAve (birthEnvVarCondX)
fecVar = apply (fecVarCondX, 2, traitAve)
survTrajecVar = apply (survTrajecVarCondX, 2, traitAve)
growthTrajecVar = apply (growthTrajecVarCondX, 2, traitAve)
envTrajecVar = apply (envTrajecVarCondX, 2, traitAve)

aveLifespan = traitAve (lifespan)
aveReproArrivalMode = traitAve (reproArrivalMode)

totSurvTrajecSkewness = sum (survTrajecSkewness)
totGrowthTrajecSkewness = sum (growthTrajecSkewness)
totEnvTrajecSkewness = sum (envTrajecSkewness)
totFecSkewness = sum (fecSkewness)

totSurvTrajecVar = sum (survTrajecVar)
totGrowthTrajecVar = sum (growthTrajecVar)
totEnvTrajecVar = sum (envTrajecVar)
totFecVar = sum (fecVar)

## Use the law of total cumulance to get the full variance and
## skewness.

## Variance
expXVarRCondX = traitAve (totVarCondX)
varXExpRCondX = traitVar (expRCondX)
## totVar agrees with old code!
totVar2 = expXVarRCondX + varXExpRCondX

######################################################################
## To get total skewness or variance, we need to create transition
## matrices and reward matrices for a model in which state is
## classified by trait x stage x env.  I'll use the prefix "ts" to
## suggest "triple state" as a way to distinguish these matrices.
##
## And since we need to start off in the ur-state, we need to include
## pre-birth states (the ur-trait, ur-stage, and
## ur-environment)), so the full prefix is bsts...
##
## OLD BUG?  I think this is fixed. [the bsts vectors give the right
## moments if we condition on x, z, q, but not if we only condition
## on, e.g., x.  And yet bstsM %*% bstsM %*% bstsM %*% urState does
## give you the appropriate triply-classified initial state (the outer
## product of m0 and groupDist).]
##
## A guide to index ranges: 1 = the ur state.  2--4: trait is assigned
## but nothing else.  5--16: trait and z are assigned, but not q.
## 17--28: trait 1, (z,q).  29--40: trait 2, (z,q).  41--52: trait 3,
## (z,q).
##
## See SI section S1 for a guide to these calculations.
##
######################################################################

tsbigmz = numTraits * bigmz
bstsbigmz = 1 + numTraits + numTraits*mz + tsbigmz

tsM = as.matrix(bdiag (list(Marray[1,,], Marray[2,,], Marray[3,,])))
tsF = as.matrix(bdiag (list(bigFarray[1,,], Marray[2,,],
                            Marray[3,,])))
bstsF = bstsM = matrix (0, bstsbigmz, bstsbigmz)
## From ur-stage to just traits (submatrix W_{21} in SI)
bstsM[1 + (1:numTraits),1] = groupDist
## From just traits to traits and stage (submatrix W_{32} in SI)
for (x in 1:numTraits) 
  bstsM[1 + numTraits + (x-1)*mz + 1:mz,1 + x] = c0
## From traits and stage to traits and stage and environment (m0)
## (submatrix W_{43} in SI)
for (x in 1:numTraits) {
  for (q in 1:numEnv) 
    bstsM[1 + numTraits + numTraits*mz + (x-1)*bigmz + (q-1)*mz + 1:mz,
          1 + numTraits + (x-1)*mz + 1:mz] =
      diag (u0[q], mz)
}
## From normal stage x env. state to normal stage x env. state
## (submatrix W_{44} in SI)
bstsM[1 + numTraits + numTraits*mz + 1:tsbigmz,
      1 + numTraits + numTraits*mz + 1:tsbigmz] = tsM

## You only start reproducing once you're past the ur-stages.  I guess
## reproduction should go to the ur-stage x ur-environment state.  Not
## that it matters much, since all we need are the column sums of bsF.
bstsF[1, 1 + numTraits + numTraits*mz + 1:tsbigmz] =
  rep(colSums (F), numTraits)

## The following reward matrix calculations come from van Daalen,
## S. F., and H. Caswell. 2017. Lifetime reproductive output:
## individual stochasticity, variance, and sensitivity
## analysis. Theoretical Ecology 10:355 – 374.

## set up bstsMplus, the transition matrix with absorbing state (dead)
bstsMplus = matrix (0, bstsbigmz+1, bstsbigmz+1)
bstsMplus[1:bstsbigmz, 1:bstsbigmz] = bstsM
bstsMplus[bstsbigmz+1, 1:bstsbigmz] = 1 - colSums(bstsM)
bstsMplus[bstsbigmz+1, bstsbigmz+1] = 1

## Set up bstsZ, which cleaves off dead states
bstsZ = cbind (diag(bstsbigmz), rep(0, bstsbigmz))

## Set up the vector 1_s, where s = bigmz + 1
bstsOneS = rep(0, bstsbigmz+1)
bstsOneS[bstsbigmz+1] = 1

## set up reward matrices
bstsR1 = bstsR2 = bstsR3 = matrix (0, bstsbigmz+1, bstsbigmz+1)
bstsR1Stripped = bstsR2Stripped = matrix (0, bstsbigmz, bstsbigmz)
##bstsBreedingStages = 1 + numTraits + numTraits*mz +
##  (0:(numTraits*numEnv - 1))*mz + 2 

## First moment of clutch size.  Note that stage 2 individuals have
## 1 chick and everyone else has zero.
##for (j in bstsBreedingStages) 
##  bstsR1[,j] = sum(bstsF[,j])
## Simpler
foo = colSums(bstsF)
for (j in 1:bstsbigmz)
  bstsR1[,j] = foo[j]

bstsR2 = bstsR1^2
bstsR3 = bstsR1^3

## R1Stripped has the absorbed state cleaved off
bstsR1Stripped = bstsR1[1:bstsbigmz,1:bstsbigmz]
bstsR2Stripped = bstsR2[1:bstsbigmz,1:bstsbigmz]

## Ex(R)
bstse = rep (1, bstsbigmz+1)
bstsN = solve (diag(bstsbigmz) - bstsM)
## I get the correct bstsRho1Vec for traits 1 and 3 (and presumably
## 2), all values of (z,q).  (E.g. compare bstsRho1Vec[17:28] with
## expRCondXZQ for x = 1 in fulmarSurvGrowthPluckPartition.R.)

## I get the correct value for each trait averaged over z and q (compare
## bstsRho1Vec[2:4] with expRCondX)!

## bstsRho1Vec[13:16] should be \Ex_q Ex(R | x=3, z, q), which should
## be the same as matrix (expRCondXZQ, 4, 3) %*% u0 when x = 3 in
## fulmarSurvGrowthPluckPartition.R and it is!
bstsRho1Vec = t(bstsN) %*% bstsZ %*% t(bstsMplus * bstsR1) %*% bstse

## Ex(R^2 | x) I get the correct bstsRho2Vec for traits 1 and 3 (and
## presumably 2), all values of (z,q).  (E.g. compare
## bstsRho2Vec[17:28] with r2 for x = 1.)
bstsRho2Vec = t(bstsN) %*% (bstsZ %*% t(bstsMplus * bstsR2) %*% bstse +
                        2*t(bstsM * bstsR1Stripped) %*% bstsRho1Vec)

## Ex(R^3 | x)
bstsRho3Vec = t(bstsN) %*% (bstsZ %*% t(bstsMplus * bstsR3) %*% bstse +
                        3*t(bstsM*bstsR2Stripped) %*% bstsRho1Vec +
                        3*t(bstsM*bstsR1Stripped) %*% bstsRho2Vec)

bstsRho1Vec = t(bstsRho1Vec)
bstsRho2Vec = t(bstsRho2Vec)
bstsRho3Vec = t(bstsRho3Vec)

## I get the correct bstsMu2Vec for trait 3, all values of (z,q).
## (Compare bstsMu2Vec[41:52] with varRCondXZQ for x = 3 in
## fulmarSurvGrowthPluckPartition.R.)
bstsMu2Vec = bstsRho2Vec - bstsRho1Vec^2
bstsMu3Vec = bstsRho3Vec - 3*bstsRho1Vec*bstsRho2Vec + 2*bstsRho1Vec^3
bstsSkewnessVec = bstsMu3Vec / bstsMu2Vec^1.5
## Fix the division by zero
##foo = which (bstsMu2Vec == 0)
##bstsSkewnessVec[foo] = 0

urState = rep(0, bstsbigmz)
urState[1] = 1
totSkewness = bstsSkewnessVec %*% urState
totVar = bstsMu2Vec %*% urState

## Sanity check: passes!
cat ("Checking totVar:", totVar, "should = ", totVar2, "\n")

traitSkewness = totSkewness - traitAve (totSkewnessCondX)
## Woot!  2022 paper says that traits contribute 19.6% of the variance
## and that's what we get here too!
traitVar = totVar - traitAve (totVarCondX)

par (lwd=2, cex.lab=1.4)

ymax = max(survTrajecSkewness, growthTrajecSkewness,
           envTrajecSkewness, fecSkewness)
ymin = min(survTrajecSkewness, growthTrajecSkewness,
           envTrajecSkewness, fecSkewness) 
plot(0:100, survTrajecSkewness[1:101], col=colors["surv"],
      lty=lineTypes["surv"],
     xlab="Age", ylab="Contributions to Skewness(R)", type="l",
     ylim=c(ymin, ymax))
lines (0:100, growthTrajecSkewness[1:101], col=colors["growth"],
      lty=lineTypes["growth"])
lines (0:100, envTrajecSkewness[1:101],
       col=colors["env"],
      lty=lineTypes["env"])
lines (0:100, fecSkewness[1:101],
       col=colors["fec"],
       lty=lineTypes["fec"])
lines (rep(aveLifespan, 2), c(0, 0.27*ymax), col="black",
       lty=1, lwd=2)
lines (rep(aveReproArrivalMode, 2), c(0, ymax), col="purple",
       lty=1, lwd=2)

legend (x=35, y=0.99*ymax, legend=c("99% of individ. are dead",
                                     "Mode(age of 1st repro.)"),
        col=c("black", "purple"),
        lty=1, bty="n", lwd=2, cex=1.2)

legend (x=35, y=0.8*ymax,
        legend=c(paste("Surv. trajec. luck: ",
                       round(100*totSurvTrajecSkewness/totSkewness, digits=1),
                       "%", sep=""),
                 paste("Env. trajec. luck: ",
                       round(100*totEnvTrajecSkewness/totSkewness, digits=1),
                       "%", sep=""),
                 paste("Growth trajec. luck: ",
                       round(100*totGrowthTrajecSkewness/totSkewness, digits=1),
                       "%", sep=""),
                 paste("Fecundity luck: ",
                       round(100*totFecSkewness/totSkewness, digits=1),
                       "%", sep="")),
        col=colors, lty=lineTypes, bty="n", lwd=2, cex=1.2)


ydiff = ymax - ymin

text (x=64, y=ymin+0.53*ydiff, col="black", cex=1.2,
        paste ("Fixed traits: ",
               round (100*traitSkewness/totSkewness, digits=2),
               "%", sep=""))

dev.copy2pdf (file="fulmarSkewnessPartition.pdf")

## And variance

dev.new ()
par( cex.lab=1.4, lwd=2)

## Now average over traits and include contribution to skewness from
## traits
ymax = max(survTrajecVar, growthTrajecVar,
           envTrajecVar, fecVar)
ymin = min(survTrajecVar, growthTrajecVar,
           envTrajecVar, fecVar) 
plot(0:100, survTrajecVar[1:101], col=colors["surv"],
      lty=lineTypes["surv"],
     xlab="Age", ylab="Contributions to Var(R)", type="l",
     lwd=2, ylim=c(ymin, ymax))
lines (0:100, growthTrajecVar[1:101], col=colors["growth"],
      lty=lineTypes["growth"])
lines (0:100, envTrajecVar[1:101],
       col=colors["env"],
       lty=lineTypes["env"])
lines (0:100, fecVar[1:101],
       col=colors["fec"],
       lty=lineTypes["fec"])
lines (rep(aveLifespan, 2), c(0, 0.35*ymax), col="black",
       lty=1, lwd=2)
lines (rep(aveReproArrivalMode, 2), c(0, ymax), col="purple",
       lty=1, lwd=2)

ydiff = ymax - ymin

legend (x=40, y=0.97*ymax, legend=c("99% of individ. are dead",
                                    "Mode(age of 1st repro.)"),
        col=c("black", "purple"),
        lty=1, bty="n", lwd=2, cex=1.2)

legend (x=40, y=0.8*ymax,
        legend=c(paste("Surv. trajec. luck: ",
                       round(100*totSurvTrajecVar/totVar, digits=1),
                       "%", sep=""),
                 paste("Env. trajec. luck: ",
                       round(100*totEnvTrajecVar/totVar, digits=1),
                       "%", sep=""),
                 paste("Growth trajec. luck: ",
                       round(100*totGrowthTrajecVar/totVar, digits=1),
                       "%", sep=""),
                 paste("Fecundity luck: ",
                       round(100*totFecVar/totVar, digits=1),
                       "%", sep="")),
        col=colors, lty=lineTypes, bty="n", lwd=2, cex=1.2)

text (x = 70, y=ymin+0.48*ydiff, col="black", cex=1.2,
      paste ("Fixed traits: ",
             round (100*traitVar/totVar, digits=2),
             "%", sep=""))

dev.copy2pdf (file="fulmarVarPartition.pdf")

###########################################

dev.new(width=8, height=7)
par(mar=c(5,4,4,4) + 0.1, cex.lab=1.4, mgp=c(2.2,1,0), # Leave space for z axis
    mfrow=c(2,2), lwd=2)
for (i in 1:3) {
  ymax = max(survTrajecSkewnessCondX[i,], growthTrajecSkewnessCondX[i,],
             envTrajecSkewnessCondX[i,], fecSkewnessCondX[i,])
  ymin = min(survTrajecSkewnessCondX[i,], growthTrajecSkewnessCondX[i,],
             envTrajecSkewnessCondX[i,], fecSkewnessCondX[i,]) 
  plot(0:100, survTrajecSkewnessCondX[i,1:101], col="black",
       lty=1, xlab="Age", ylab="Contributions to Skewness(R)", type="l",
       lwd=2, main=paste("Group", i, sep=" "), ylim=c(ymin, ymax))
  lines (0:100, growthTrajecSkewnessCondX[i,1:101], col="orange",
         lty=1, lwd=2, main=paste("Group", i, sep=" "))
  lines (rep(lifespan[i], 2), c(0, 0.5*ymax), col="black",
         lty=2, lwd=2)
  lines (0:100, envTrajecSkewnessCondX[i,1:101],
         col="green3", lty=1, type="l", lwd=2)
  lines (0:100, fecSkewnessCondX[i,1:101],
         col="blue", lty=1, type="l", lwd=2)
  if (i == 3) {
    legend (x="topright",
            legend=c("Survival trajectory",
                     "Growth trajectory",
                     "Env. trajectory",
                     "Fecundity",
                     "99% of individ. are dead"),
            col=c("black", "orange", "green", "blue", "black"),
            lty=c(1,1,1,1,2), lwd=2, bty="n")
  }
}

## Now average over traits and include contribution to skewness from
## traits
ymax = max(survTrajecSkewness, growthTrajecSkewness,
           envTrajecSkewness, fecSkewness)
ymin = min(survTrajecSkewness, growthTrajecSkewness,
           envTrajecSkewness, fecSkewness) 
plot(0:100, survTrajecSkewness[1:101], col="black",
     lty=1, xlab="Age", ylab="Contributions to Skewness(R)", type="l",
     lwd=2, main="All groups", ylim=c(ymin, ymax))
lines (0:100, growthTrajecSkewness[1:101], col="orange",
       lty=1, lwd=2)
lines (rep(lifespan[i], 2), c(0, 0.5*ymax), col="black",
       lty=2, lwd=2)
lines (0:100, envTrajecSkewness[1:101],
       col="green3", lty=1, type="l", lwd=2)
lines (0:100, fecSkewness[1:101],
       col="blue", lty=1, type="l", lwd=2)

ydiff = ymax - ymin

text (x=70, y=ymin+0.95*ydiff, col="purple",
      paste("Birth env.: ",
            round(100*birthEnvSkewness/totSkewness, digits=2), "%", sep=""))
text (x=70, y=ymin+0.85*ydiff, col="green3",
      paste("Env. trajec.: ",
            round(100*totEnvTrajecSkewness/totSkewness, digits=2),
            "%", sep=""))
text (x=70, y=ymin+0.75*ydiff, col="blue",
      paste("Fecundity: ",
            round(100*totFecSkewness/totSkewness, digits=2),
            "%", sep=""))
text (x = 70, y=ymin+0.65*ydiff, col="red",
      paste ("Fixed traits: ",
             round (100*traitSkewness/totSkewness, digits=2),
             "%", sep=""))
text (x=70, y=ymin+0.55*ydiff, col="orange",
      paste("Growth. trajec.: ",
            round(100*totGrowthTrajecSkewness/totSkewness, digits=2),
            "%", sep="")) 
text (x=70, y=ymin+0.45*ydiff,
      paste("Surv. trajec.: ",
            round(100*totSurvTrajecSkewness/totSkewness, digits=1),
            
            "%", sep=""))


## And variance

dev.new(width=8, height=7)
par(mar=c(5,4,4,4) + 0.1, cex.lab=1.4, mgp=c(2.2,1,0), # Leave space for z axis
    mfrow=c(2,2), lwd=2)
for (i in 1:3) {
  ymax = max(survTrajecVarCondX[i,], growthTrajecVarCondX[i,],
             envTrajecVarCondX[i,], fecVarCondX[i,])
  ymin = min(survTrajecVarCondX[i,], growthTrajecVarCondX[i,],
             envTrajecVarCondX[i,], fecVarCondX[i,]) 
  plot(0:100, survTrajecVarCondX[i,1:101], col="black",
       lty=1, xlab="Age", ylab="Contributions to Var(R)", type="l",
       lwd=2, main=paste("Group", i, sep=" "), ylim=c(ymin, ymax))
  lines (0:100, growthTrajecVarCondX[i,1:101], col="orange",
         lty=1, lwd=2, main=paste("Group", i, sep=" "))
  lines (rep(lifespan[i], 2), c(0, 0.5*ymax), col="black",
         lty=2, lwd=2)
  lines (0:100, envTrajecVarCondX[i,1:101],
         col="green3", lty=1, type="l", lwd=2)
  lines (0:100, fecVarCondX[i,1:101],
         col="blue", lty=1, type="l", lwd=2)
  if (i == 3) {
    legend (x="topright",
            legend=c("Survival trajectory",
                     "Growth trajectory",
                     "Env. trajectory",
                     "Fecundity",
                     "99% of individ. are dead"),
            col=c("black", "orange", "green", "blue", "black"),
            lty=c(1,1,1,1,2), lwd=2, bty="n")
  }
}

## Now average over traits and include contribution to skewness from
## traits
ymax = max(survTrajecVar, growthTrajecVar,
           envTrajecVar, fecVar)
ymin = min(survTrajecVar, growthTrajecVar,
           envTrajecVar, fecVar) 
plot(0:100, survTrajecVar[1:101], col="black",
     lty=1, xlab="Age", ylab="Contributions to Var(R)", type="l",
     lwd=2, main="All groups", ylim=c(ymin, ymax))
lines (0:100, growthTrajecVar[1:101], col="orange",
       lty=1, lwd=2)
lines (rep(lifespan[i], 2), c(0, 0.5*ymax), col="black",
       lty=2, lwd=2)
lines (0:100, envTrajecVar[1:101],
       col="green3", lty=1, type="l", lwd=2)
lines (0:100, fecVar[1:101],
       col="blue", lty=1, type="l", lwd=2)

ydiff = ymax - ymin

text (x=70, y=ymin+0.95*ydiff, col="purple",
      paste("Birth env.: ",
            round(100*birthEnvVar/totVar, digits=2), "%", sep=""))
text (x=70, y=ymin+0.85*ydiff, col="green3",
      paste("Env. trajec.: ",
            round(100*totEnvTrajecVar/totVar, digits=2),
            "%", sep=""))
text (x=70, y=ymin+0.75*ydiff, col="blue",
      paste("Fecundity: ",
            round(100*totFecVar/totVar, digits=2),
            "%", sep=""))
text (x = 70, y=ymin+0.65*ydiff, col="red",
      paste ("Fixed traits: ",
             round (100*traitVar/totVar, digits=2),
             "%", sep=""))
text (x=70, y=ymin+0.55*ydiff, col="orange",
      paste("Growth. trajec.: ",
            round(100*totGrowthTrajecVar/totVar, digits=2),
            "%", sep="")) 
text (x=70, y=ymin+0.45*ydiff,
      paste("Surv. trajec.: ",
            round(100*totSurvTrajecVar/totVar, digits=1),
            "%", sep=""))

##dev.copy2pdf (file="fulmarVarPartition.pdf")

dev.new ()
par (cex.lab=1.4)
skewnessContributions = c(totSurvTrajecSkewness,
                          totGrowthTrajecSkewness,
                          totEnvTrajecSkewness,
                          totFecSkewness,
                          traitSkewness)
barplot (skewnessContributions,
         col=c("black", "orange", "green", "blue", "red"), main="Fulmars",
         names.arg=c("Survival", "Growth", "Env.", "Fecundity",
                     "Traits"),
         cex.names=1.4, cex.main=1.4)

dev.copy2pdf (file="fulmarSkewnessBarplot.pdf")
```
