## Supplementary material for "To prosper, live long: understanding the sources of reproductive skew and extreme reproductive success in structured populations": Associated R code: genericPartitionVarSkewness6.R

```
#########################################################################
##
## genericPartitionVarSkewness6: various utilities for calculations in
## "To prosper, live long: ..."  Does most of the actual work.
##
## Assumes that clutch size is Poisson-distributed.
##
## Dependencies: partitioningUtilities.R, megamatrixFunctions.R
##
## Some general definitions:
##
## mz: number of stages/sizes
##
## numEnv: number of environments/year types
##
## P: survival/growth transition matrix
##
## c0: initial distribution of sizes/stages
##
## F: fecundity matrix
##
## Q: environment transition matrix
## 
## M: "megamatrix" governing size x environment transitions
##
## m0: initial distribution of size x environment
##
##########################################################################

require ("Matrix")

## Useful for getting modal time to first reproduction when states are
## defined by reproductive status.
getModalTimeToHitState = function (absorbingStates,
                                   Plist, Flist, Q=NULL,
                                   m0, maxAge=100) {
  mz = dim(Plist[[1]])[1]
  if (is.null(Q)) {  ## no env. var.
    bigmz = mz
    M = P
  } else {
    numEnv = dim(Q)[1]
    bigmz = numEnv*mz
    M = matrix (0, bigmz, bigmz)
    for (i in 1:numEnv) {
      for (j in 1:numEnv) {
        M[(i-1)*mz + 1:mz, (j-1)*mz + 1:mz] = Plist[[j]]*Q[i,j]
        F[(i-1)*mz + 1:mz, (j-1)*mz + 1:mz] = Flist[[j]]*Q[i,j]
      }
    }
  }

  ## Create kernel conditional on hitting an absorbing state

  ## define transient states (i.e. not absorbing)
  transient = (1:bigmz)[-absorbingStates]
  numTransient = length(transient)

  out = makeCondKernel (M, transient)
  MCond = out$MCond 

  ## Create kernel with all absorbing states going directly to a
  ## special absorbing state.
  MAbsorbing = matrix (0, bigmz+1, bigmz+1)
  MAbsorbing[transient, transient] = MCond[transient, transient]
  for (j in transient)
    MAbsorbing[bigmz+1, j] = sum(MCond[absorbingStates, j])
  MAbsorbing[bigmz+1, absorbingStates] = 1
  MAbsorbing[bigmz+1, bigmz+1] = 1

  ## Get distribution of times to absorption
  m0Absorbing = c(m0, 0)
  MaAbsorbingM0 = matrix (0, maxAge, bigmz+1)
  MaAbsorbingM0[1,] = m0Absorbing

  ## get stage x env. distribution at each age
  for (a in 2:maxAge)
    MaAbsorbingM0[a,] = MAbsorbing %*% MaAbsorbingM0[a-1,]

  hitTimes = diff (MaAbsorbingM0[,bigmz+1])
  hitTimeMode = which.max (hitTimes)
  hitTimeDist = hitTimes / sum(hitTimes)

  return (out = list(hitTimeMode=hitTimeMode,
                     hitTimeDist=hitTimeDist))
}

## Get mode of time to first reproductive event, defined as the first
## time an individual successfully produces offspring
getModalTimeToFirstRepro = function (Plist, Flist, Q, m0, 
                                     maxAge=100) {

  mz = dim(Plist[[1]])[1]
  numEnv = dim(Q)[1]
  bigmz = numEnv*mz

  ## Define megamatrices M and F
  F = M = matrix (0, bigmz, bigmz)
  for (i in 1:numEnv) {
    for (j in 1:numEnv) {
      M[(i-1)*mz + 1:mz, (j-1)*mz + 1:mz] = Plist[[j]]*Q[i,j]
      F[(i-1)*mz + 1:mz, (j-1)*mz + 1:mz] = Flist[[j]]*Q[i,j]
    }
  }

  ## pb = prob. of breeding as a function of state
  ## Assume entries of F are means of Poisson distributions, so
  ## prob. of breeding is pb = 1 - exp(-mean)
  pb = 1 - exp(-colSums (F))

  big3mz = 3*bigmz
  bigF = matrix (0, big3mz, big3mz)
  bigM = matrix (0, big3mz, big3mz);
  
  ## Make M for the extended state space Z1 \cup Z2 \cup Z3.  Z1 = has
  ## never reproduced, Z2 = reproduced for the first time this year,
  ## Z3 = reproduced for the first time sometime in the past.  (See
  ## Snyder and Ellner Am. Nat. 2021, SI section "Conditioning on
  ## reaching maturity, attempting breeding, or producing an
  ## offspring", "Breeder definition 3".
  Z1 = 1:bigmz
  Z2 = (bigmz+1):(2*bigmz)
  Z3 = (2*bigmz+1):(3*bigmz)
  Z23 = c(Z2, Z3)
  
  # Z1 to Z1
  for (j in Z1) bigM[j, Z1] = M[j,]*(1 - pb[j])
  # Z1 to Z2 
  for (j in Z1) bigM[bigmz+j, Z1] = M[j,]*pb[j]
  # Z2 to Z3
  bigM[Z3, Z2] = M
  # Z3 to Z3
  bigM[Z3, Z3] = M

  ## expanded state space init. distribution
  bigm0 = c(m0, rep(0, 2*bigmz))

  ## make F for the extended state space Z1 \cup Z2 \cup Z3.  Note that the
  ## only non-zero entries are from Z2 and Z3 to Z1.
  bigF[1, Z23] = F[1,]

  ## Calculate aM, the probability of going from Z1 to Z2 in one step
  aM = apply (bigM[(bigmz+1):(2*bigmz), 1:bigmz], 2, sum)
  
  ## Q2 = M restricted to Z1, N2 = (I - Q2)^{-1}
  Q2 = bigM[1:bigmz,1:bigmz];

  N2 = solve(diag(bigmz) - Q2)

  ## Calculate q2Extended, the probability of reaching Z2 before
  ## death.
  q2Extended = c(aM %*% N2, rep(1, 2*bigmz))

  ## Transition matrix conditional on becoming big before death
  bigMCond = matrix (0, big3mz, big3mz)
  for (j in 1:big3mz)
    bigMCond[,j] = bigM[,j] * q2Extended / q2Extended[j]

  ## For calculating repro. states hitting time dist.  All repro. states
  ## transition to a special absorbing state.
  bigMReproAbsorbing = matrix (0, big3mz+1, big3mz+1)
  ## Transitions from never reproduced to never reproduced
  bigMReproAbsorbing[Z1, Z1] = bigMCond[Z1, Z1]
  ## Transitions from never reproduced to the absorbing state
  for (j in Z1)
    bigMReproAbsorbing[big3mz+1, j] = sum(bigMCond[Z23, j])
  ## Transitions from reproduced this year or sometime in the past to
  ## the absorbing state
  bigMReproAbsorbing[big3mz+1, Z23] = 1
  ## Transitions from the absorbing state to the absorbing state
  bigMReproAbsorbing[big3mz+1, big3mz+1] = 1

  bigMaAbsorbing = diag(1, big3mz+1)
  bigm0Absorbing = c(bigm0, 0)
  bigMaAbsorbingM0 = matrix (0, maxAge, big3mz+1)
  bigMaAbsorbingM0[1,] = bigm0Absorbing

  ## get stage x env. distribution at each age
  for (a in 2:maxAge) {
    bigMaAbsorbing = bigMReproAbsorbing %*% bigMaAbsorbing
    bigMaAbsorbingM0[a,] = bigMaAbsorbing %*% bigm0Absorbing
  }

  ## calculate big size hit time distribution
  reproArrivalAges = diff(bigMaAbsorbingM0[,big3mz+1])
  reproArrivalMode = which.max(reproArrivalAges)
  reproAgeDist = reproArrivalAges / sum(reproArrivalAges)

  return (out=list(reproArrivalMode=reproArrivalMode,
                   reproAgeDist=reproAgeDist,
                   q2Extended=q2Extended))
}

getVarSkewnessPartitionsNoEnvVar = function (P, F, c0, maxAge=100,
                                             esR1=NULL, esR2=NULL,
                                             esR3=NULL,
                                             bsR1=NULL, bsR2=NULL,
                                             bsR3=NULL, debugging=FALSE) {
  # Input parameters:
  # P is the survival/growth transition matrix (called U in COMPADRE)
  # F is the matrix of fertility transitions
  # c0 is the birth state distribution (also called mixing distribution)

  if (debugging) {
    ## some toy matrices for testing things out:
    P<- matrix(c(0.5, 0.07, 0.05, 0, 0.5, 0.21, 0, 0, 0.8), ncol=3)
    F<- matrix(c(0.2, 0, 0, 1, 0.5, 0, 5, 3, 0), ncol=3)
    
    c0 = c(0.9, 0.1, 0)
    maxAge = 100
  }

  mz = dim(P)[1]

  ## Make survival, growth, and fecundity bullet matrices
  Sbullet = diag (colSums (P))
  
  Gbullet = P;  
  for(ic in 1:ncol(Gbullet)){
        if(Sbullet[ic,ic]>0) Gbullet[,ic]=Gbullet[,ic]/Sbullet[ic,ic]
  }       
  cat(range(Gbullet%*%Sbullet-P), "should equal 0", "\n"); 
  
  ## #kids isn't part of the state definition, so Fbullet = I
  Fbullet = diag (mz)

  ## create matrices for extended state space, which I will denote with
  ## the prefix "es".

  esmz = 3*mz
  esF = esP = matrix (0, esmz, esmz)

  esP[1:mz, 2*mz + 1:mz] = Gbullet
  esP[mz + 1:mz, 1:mz] = Fbullet
  esP[2*mz + 1:mz, mz + 1:mz] = Sbullet
  
  esF[mz + 1:mz, 1:mz] = F

  ################################################################
  ## Extended state model that includes ur-stage alpha_z.  Used for
  ## calculating contributions from birth stage.  The "bs" before
  ## variable names stands for "birth state."
  ################################################################

  bsmz = 1 + mz
  ## initial state --- everybody starts off in the ur-state.
  bsC0 = rep(0, bsmz)
  bsC0[1] = 1

  bsF = bsP = matrix (0, bsmz, bsmz)
  ## From ur-state to birth state
  bsP[1 + (1:mz),1] = c0
  ## From normal state to normal state
  bsP[1 + 1:mz, 1 + 1:mz] = P

  ## You only start reproducing once you're past the ur-state.  Note
  ## that all we need are the column sums of bsF.
  bsF[1, 1 + 1:mz] = colSums (F)

  ## set up reward matrices ###############################

  ## If the reward matrices aren't specified, we assume
  ## Poisson-distributed clutch sizes
  if (is.null(esR1)) {
    esR1 = esR2 = esR3 = matrix (0, esmz+1, esmz+1)
    bsR1 = bsR2 = bsR3 = matrix (0, bsmz+1, bsmz+1)

    ## First moment of clutch size (Poisson)
    for (j in 1:esmz) 
      esR1[,j] = sum(esF[,j])
    for (j in 1:bsmz) 
      bsR1[,j] = sum(bsF[,j])

    ## Second moment of clutch size  
    esR2 = esR1 + esR1^2
    bsR2 = bsR1 + bsR1^2
    ## Third moment of clutch size
    esR3 = esR1+ 3*esR1*esR1 + esR1*esR1*esR1
    bsR3 = bsR1+ 3*bsR1*bsR1 + bsR1*bsR1*bsR1
  }
  
  ## Get moments conditional on init. state #########################

  out = calcMoments (esP, esR1, esR2, esR3)
  esRho1Vec = out$rho1Vec
  esMu2Vec = out$mu2Vec
  esSkewnessVec = out$skewnessVec

  out = calcMoments (bsP, bsR1, bsR2, bsR3)
  bsRho1Vec = out$rho1Vec
  bsMu2Vec = out$mu2Vec
  bsMu3Vec = out$mu3Vec
  bsSkewnessVec = out$skewnessVec

  ## Start calculating luck terms ###################################

  survTrajecSkewness = growthTrajecSkewness = 
    fecSkewness = numeric (maxAge)
  fecUpdateSkewness = survUpdateSkewness =
    growthUpdateSkewness =  numeric (maxAge)
  survUpdateVar = growthUpdateVar = fecUpdateVar =
    numeric (maxAge) 
  survTrajecVar = growthTrajecVar = fecVar =
    numeric (maxAge)
  surv = numeric (maxAge)

  ## for ur-state var.
  ## expectation wrt z of Var(success | z)
  expZVarCondZ = bsMu2Vec %*% bsC0
  ## variance wrt z of Exp(success | z)
  varZExpCondZ = sum(bsC0*bsRho1Vec^2) - (sum(bsC0*bsRho1Vec)^2)
  ## Law of total variance
  urVar = expZVarCondZ + varZExpCondZ
  
  justBirthStateVar = bsMu2Vec %*% bsP %*% bsC0
  birthStateVar = urVar - justBirthStateVar

  ## for ur-state skewness
  expZMu3CondZ = bsMu3Vec %*% bsC0
  ## 3rd moment wrt z of Exp(success | z)
  rho3ZExpCondZ = sum(bsC0*(bsRho1Vec)^3)
  ## 2nd moment wrt z of Exp(success | z)
  rho2ZExpCondZ = sum(bsC0*(bsRho1Vec)^2)
  ## 1st moment wrt z of Exp(success | z)
  expZExpCondZ = sum(bsC0*bsRho1Vec)
  ## 3rd central moment wrt z of Exp(success | z)
  mu3ZExpCondZ = rho3ZExpCondZ -
    3*expZExpCondZ*rho2ZExpCondZ +
    2*expZExpCondZ^3
  ## covariance wrt z of Exp(success | z) and Var(success | z)
  covZExpVarCondZ = sum(bsC0*bsRho1Vec*bsMu2Vec) -
    sum(bsC0*bsRho1Vec)*sum(bsC0*bsMu2Vec)
  ## Law of total cumulance
  urMu3 = expZMu3CondZ + mu3ZExpCondZ +
    3*covZExpVarCondZ
  urSkewness = urMu3 / urVar^1.5
  
  justBirthStateSkewness = bsSkewnessVec %*% bsP %*% bsC0
  birthStateSkewness = urSkewness - justBirthStateSkewness
  
  ## Starting the age loop
  PaC0 = c0  ## P^a c_0
  mzZero = rep(0, mz)

  ## Sanity check: passes
  if (debugging) {
    N = solve(diag(mz) - P)
    rho1Vec = colSums (F %*% N)
    cat (esRho1Vec %*% c(c0, mzZero, mzZero),
         "should = ", rho1Vec %*% c0, "\n")
  }

  fecUpdateSkewness[1] = esSkewnessVec %*% c(mzZero, PaC0, mzZero)
  survUpdateSkewness[1] = esSkewnessVec %*% 
    c(mzZero, mzZero, Sbullet %*% PaC0)
  growthUpdateSkewness[1] = esSkewnessVec %*% 
    c(P %*% PaC0, mzZero, mzZero)

  fecUpdateVar[1] = esMu2Vec %*% c(mzZero, PaC0, mzZero)
  survUpdateVar[1] = esMu2Vec %*% c(mzZero, mzZero, Sbullet %*% PaC0)
  growthUpdateVar[1] = esMu2Vec %*% c(P %*% PaC0, mzZero, mzZero)

  surv[1] = sum (PaC0)
  ## a is actually age + 1, since the first year of life is age 0
  for (a in 2:maxAge) {
    PaC0 = P %*% PaC0

    fecUpdateSkewness[a] = esSkewnessVec %*% c(mzZero, PaC0, mzZero)
    survUpdateSkewness[a] = esSkewnessVec %*% 
      c(mzZero, mzZero, Sbullet %*% PaC0)
    growthUpdateSkewness[a] = esSkewnessVec %*% 
      c(P %*% PaC0, mzZero, mzZero)

    fecUpdateVar[a] = esMu2Vec %*% c(mzZero, PaC0, mzZero)
    survUpdateVar[a] = esMu2Vec %*% c(mzZero, mzZero, Sbullet %*% PaC0)
    growthUpdateVar[a] = esMu2Vec %*% c(P %*% PaC0, mzZero, mzZero)

    ## fecundity skewness and var.
    fecSkewness[a] = growthUpdateSkewness[a-1] - fecUpdateSkewness[a]
    fecVar[a] = growthUpdateVar[a-1] - fecUpdateVar[a]
    
    ## survival trajectory skewness and var.
    survTrajecSkewness[a-1] = fecUpdateSkewness[a-1] -
      survUpdateSkewness[a-1]
    survTrajecVar[a-1] = fecUpdateVar[a-1] - survUpdateVar[a-1]
    
    ## growth trajectory skewness and var.
    growthTrajecSkewness[a-1] = survUpdateSkewness[a-1] -
      growthUpdateSkewness[a-1] 
    growthTrajecVar[a-1] = survUpdateVar[a-1] - growthUpdateVar[a-1]

    surv[a] = sum(PaC0)
  }

  ## By what age are 99% of individuals dead?
  lifespan = which(surv < 0.01)[1]

  ## And get the first value of fecSkewness and fecVar
  fecSkewness[1] = justBirthStateSkewness - fecUpdateSkewness[1]
  fecVar[1] = justBirthStateVar - fecUpdateVar[1]

  ## By what age are 99% of individuals dead?
  lifespan = which(surv < 0.01)[1]

  totSurvTrajecSkewness = sum(survTrajecSkewness)
  totGrowthTrajecSkewness = sum(growthTrajecSkewness)
  totFecSkewness = sum(fecSkewness)
  totSkewness = bsSkewnessVec %*% bsC0
  ## Sanity check: passes
  if (debugging) 
    cat (totSurvTrajecSkewness + totGrowthTrajecSkewness +
         totFecSkewness + birthStateSkewness, "should = ",
         totSkewness, "\n")

  totSurvTrajecVar = sum(survTrajecVar)
  totGrowthTrajecVar = sum(growthTrajecVar)
  totFecVar = sum(fecVar)
  totVar = bsMu2Vec %*% bsC0
  ## Sanity check: passes
  if (debugging)
    cat (totSurvTrajecVar + totGrowthTrajecVar +
         totFecVar + birthStateVar, "should = ",
         totVar, "\n")

  return (out=list(birthStateVar=birthStateVar,
                   survTrajecVar=survTrajecVar,
                   growthTrajecVar=growthTrajecVar,
                   fecVar=fecVar,
                   totVar=totVar,
                   birthStateSkewness=birthStateSkewness,
                   survTrajecSkewness=survTrajecSkewness,
                   growthTrajecSkewness=growthTrajecSkewness,
                   fecSkewness=fecSkewness,
                   totSkewness=totSkewness,
                   lifespan=lifespan))
}

getVarMu3PartitionsNoEnvVar = function (P, F, c0, maxAge=100,
                                             esR1=NULL, esR2=NULL,
                                             esR3=NULL,
                                             bsR1=NULL, bsR2=NULL,
                                             bsR3=NULL, debugging=TRUE) {
  # Input parameters:
  # P is the survival/growth transition matrix (called U in COMPADRE)
  # F is the matrix of fertility transitions
  # c0 is the birth state distribution (also called mixing distribution)

  mz = dim(P)[1]

  ## Make survival, growth, and fecundity bullet matrices, defined in
  ## "Age-partitioning of skewness contributions from different luck
  ## types"
  Sbullet = diag (colSums (P))
  
  Gbullet = P;  
  for(ic in 1:ncol(Gbullet)){
        if(Sbullet[ic,ic]>0) Gbullet[,ic]=Gbullet[,ic]/Sbullet[ic,ic]
  }       
  cat(range(Gbullet%*%Sbullet-P), "should equal 0", "\n"); 
  
  ## #kids isn't part of the state definition, so Fbullet = I
  Fbullet = diag (mz)

  ## create matrices for extended state space, which I will denote with
  ## the prefix "es".

  esmz = 3*mz
  esF = esP = matrix (0, esmz, esmz)

  esP[1:mz, 2*mz + 1:mz] = Gbullet
  esP[mz + 1:mz, 1:mz] = Fbullet
  esP[2*mz + 1:mz, mz + 1:mz] = Sbullet
  
  esF[mz + 1:mz, 1:mz] = F

  ################################################################
  ## Extended state model that includes ur-stage alpha_z.  Used for
  ## calculating contributions from birth stage.  The "bs" before
  ## variable names stands for "birth state."
  ################################################################

  bsmz = 1 + mz
  ## initial state --- everybody starts off in the ur-state.
  bsC0 = rep(0, bsmz)
  bsC0[1] = 1

  bsF = bsP = matrix (0, bsmz, bsmz)
  ## From ur-state to birth state
  bsP[1 + (1:mz),1] = c0
  ## From normal state to normal state
  bsP[1 + 1:mz, 1 + 1:mz] = P

  ## You only start reproducing once you're past the ur-state.  Note
  ## that all we need are the column sums of bsF.
  bsF[1, 1 + 1:mz] = colSums (F)

  ## set up reward matrices ###############################

  ## If the reward matrices aren't specified, we assume
  ## Poisson-distributed clutch sizes
  if (is.null(esR1)) {
    esR1 = esR2 = esR3 = matrix (0, esmz+1, esmz+1)
    bsR1 = bsR2 = bsR3 = matrix (0, bsmz+1, bsmz+1)

    ## First moment of clutch size (Poisson)
    for (j in 1:esmz) 
      esR1[,j] = sum(esF[,j])
    for (j in 1:bsmz) 
      bsR1[,j] = sum(bsF[,j])

    ## Second moment of clutch size  
    esR2 = esR1 + esR1^2
    bsR2 = bsR1 + bsR1^2
    ## Third moment of clutch size
    esR3 = esR1+ 3*esR1*esR1 + esR1*esR1*esR1
    bsR3 = bsR1+ 3*bsR1*bsR1 + bsR1*bsR1*bsR1
  }
  
  ## Get moments conditional on init. state #########################

  out = calcMoments (esP, esR1, esR2, esR3)
  esRho1Vec = out$rho1Vec
  esMu2Vec = out$mu2Vec
  esMu3Vec = out$mu3Vec

  out = calcMoments (bsP, bsR1, bsR2, bsR3)
  bsRho1Vec = out$rho1Vec
  bsMu2Vec = out$mu2Vec
  bsMu3Vec = out$mu3Vec

  ## Start calculating luck terms ###################################

  survTrajecMu3 = growthTrajecMu3 = 
    fecMu3 = numeric (maxAge)
  fecUpdateMu3 = survUpdateMu3 =
    growthUpdateMu3 =  numeric (maxAge)
  survUpdateVar = growthUpdateVar = fecUpdateVar =
    numeric (maxAge) 
  survTrajecVar = growthTrajecVar = fecVar =
    numeric (maxAge)
  surv = numeric (maxAge)

  ## for ur-state var.
  ## expectation wrt z of Var(success | z)
  expZVarCondZ = bsMu2Vec %*% bsC0
  ## variance wrt z of Exp(success | z)
  varZExpCondZ = sum(bsC0*bsRho1Vec^2) - (sum(bsC0*bsRho1Vec)^2)
  ## Law of total variance
  urVar = expZVarCondZ + varZExpCondZ
  
  justBirthStateVar = bsMu2Vec %*% bsP %*% bsC0
  birthStateVar = urVar - justBirthStateVar

  ## for ur-state skewness
  expZMu3CondZ = bsMu3Vec %*% bsC0
  ## 3rd moment wrt z of Exp(success | z)
  rho3ZExpCondZ = sum(bsC0*(bsRho1Vec)^3)
  ## 2nd moment wrt z of Exp(success | z)
  rho2ZExpCondZ = sum(bsC0*(bsRho1Vec)^2)
  ## 1st moment wrt z of Exp(success | z)
  expZExpCondZ = sum(bsC0*bsRho1Vec)
  ## 3rd central moment wrt z of Exp(success | z)
  mu3ZExpCondZ = rho3ZExpCondZ -
    3*expZExpCondZ*rho2ZExpCondZ +
    2*expZExpCondZ^3
  ## covariance wrt z of Exp(success | z) and Var(success | z)
  covZExpVarCondZ = sum(bsC0*bsRho1Vec*bsMu2Vec) -
    sum(bsC0*bsRho1Vec)*sum(bsC0*bsMu2Vec)
  ## Law of total cumulance
  urMu3 = expZMu3CondZ + mu3ZExpCondZ +
    3*covZExpVarCondZ
  
  justBirthStateMu3 = bsMu3Vec %*% bsP %*% bsC0
  birthStateMu3 = urMu3 - justBirthStateMu3
  
  ## Starting the age loop
  PaC0 = c0  ## P^a c_0
  mzZero = rep(0, mz)

  ## Sanity check: passes
  if (debugging) {
    N = solve(diag(mz) - P)
    rho1Vec = colSums (F %*% N)
    cat (esRho1Vec %*% c(c0, mzZero, mzZero),
         "should = ", rho1Vec %*% c0, "\n")
  }

  fecUpdateMu3[1] = esMu3Vec %*% c(mzZero, PaC0, mzZero)
  survUpdateMu3[1] = esMu3Vec %*% 
    c(mzZero, mzZero, Sbullet %*% PaC0)
  growthUpdateMu3[1] = esMu3Vec %*% 
    c(P %*% PaC0, mzZero, mzZero)

  fecUpdateVar[1] = esMu2Vec %*% c(mzZero, PaC0, mzZero)
  survUpdateVar[1] = esMu2Vec %*% c(mzZero, mzZero, Sbullet %*% PaC0)
  growthUpdateVar[1] = esMu2Vec %*% c(P %*% PaC0, mzZero, mzZero)

  surv[1] = sum (PaC0)
  ## a is actually age + 1, since the first year of life is age 0
  for (a in 2:maxAge) {
    PaC0 = P %*% PaC0

    fecUpdateMu3[a] = esMu3Vec %*% c(mzZero, PaC0, mzZero)
    survUpdateMu3[a] = esMu3Vec %*% 
      c(mzZero, mzZero, Sbullet %*% PaC0)
    growthUpdateMu3[a] = esMu3Vec %*% 
      c(P %*% PaC0, mzZero, mzZero)

    fecUpdateVar[a] = esMu2Vec %*% c(mzZero, PaC0, mzZero)
    survUpdateVar[a] = esMu2Vec %*% c(mzZero, mzZero, Sbullet %*% PaC0)
    growthUpdateVar[a] = esMu2Vec %*% c(P %*% PaC0, mzZero, mzZero)

    ## fecundity skewness and var.
    fecMu3[a] = growthUpdateMu3[a-1] - fecUpdateMu3[a]
    fecVar[a] = growthUpdateVar[a-1] - fecUpdateVar[a]
    
    ## survival trajectory skewness and var.
    survTrajecMu3[a-1] = fecUpdateMu3[a-1] -
      survUpdateMu3[a-1]
    survTrajecVar[a-1] = fecUpdateVar[a-1] - survUpdateVar[a-1]
    
    ## growth trajectory skewness and var.
    growthTrajecMu3[a-1] = survUpdateMu3[a-1] -
      growthUpdateMu3[a-1] 
    growthTrajecVar[a-1] = survUpdateVar[a-1] - growthUpdateVar[a-1]

    surv[a] = sum(PaC0)
  }

  ## By what age are 99% of individuals dead?
  lifespan = which(surv < 0.01)[1]

  ## And get the first value of fecMu3 and fecVar
  fecMu3[1] = justBirthStateMu3 - fecUpdateMu3[1]
  fecVar[1] = justBirthStateVar - fecUpdateVar[1]

  ## By what age are 99% of individuals dead?
  lifespan = which(surv < 0.01)[1]

  totSurvTrajecMu3 = sum(survTrajecMu3)
  totGrowthTrajecMu3 = sum(growthTrajecMu3)
  totFecMu3 = sum(fecMu3)
  totMu3 = bsMu3Vec %*% bsC0

  cat ("Got here!\n")
  ## Sanity check: passes
  if (debugging) 
    cat ("Checking Mu3 totals:", totSurvTrajecMu3 + totGrowthTrajecMu3 +
         totFecMu3 + birthStateMu3, "should = ",
         totMu3, "\n")

  totSurvTrajecVar = sum(survTrajecVar)
  totGrowthTrajecVar = sum(growthTrajecVar)
  totFecVar = sum(fecVar)
  totVar = bsMu2Vec %*% bsC0
  ## Sanity check: passes
  if (debugging)
    cat ("Checking Var totals:", totSurvTrajecVar + totGrowthTrajecVar +
         totFecVar + birthStateVar, "should = ",
         totVar, "\n")

  return (out=list(birthStateVar=birthStateVar,
                   survTrajecVar=survTrajecVar,
                   growthTrajecVar=growthTrajecVar,
                   fecVar=fecVar,
                   totVar=totVar,
                   birthStateMu3=birthStateMu3,
                   survTrajecMu3=survTrajecMu3,
                   growthTrajecMu3=growthTrajecMu3,
                   fecMu3=fecMu3,
                   totMu3=totMu3,
                   lifespan=lifespan))
}


getVarSkewnessPartitionsEnvVar = function (Plist, Flist, Q, m0,
                                           maxAge=100, esR1=NULL,
                                           esR2=NULL, esR3=NULL,
                                           bsR1=NULL, bsR2=NULL,
                                           bsR3=NULL)
{
  require (Matrix)
  debugging = TRUE
  
  mz = dim(Plist[[1]])[1]
  numEnv = dim(Q)[1]
  bigmz = numEnv*mz

  ## Define megamatrices M and F
  F = M = matrix (0, bigmz, bigmz)
  for (i in 1:numEnv) {
    for (j in 1:numEnv) {
      M[(i-1)*mz + 1:mz, (j-1)*mz + 1:mz] = Plist[[j]]*Q[i,j]
      F[(i-1)*mz + 1:mz, (j-1)*mz + 1:mz] = Flist[[j]]*Q[i,j]
    }
  }

  ## Make survival and growth matrices for each environment type
  Slist = Glist = list (numEnv)
  for (q in 1:numEnv) {
    Slist[[q]] = diag (colSums (Plist[[q]]))
    Sinv = diag (1 / diag(Slist[[q]]))
    Glist[[q]] = Plist[[q]] %*% Sinv
  }

  ## Make "bullet" matrices, defined in "Age-partitioning of skewness
  ## contributions from different luck types."  bdiag makes sparse
  ## matrices, which can't be pasted into regular matrices, so wrap
  ## these in as.matrix().
  Sbullet = as.matrix(bdiag (Slist))
  Gbullet = as.matrix(bdiag (Glist))
  Pbullet = Gbullet %*% Sbullet
  Qbullet = matrix (0, bigmz, bigmz)
  for (i in 1:numEnv) {
    for (j in 1:numEnv) {
      Qbullet[(i-1)*mz + 1:mz, (j-1)*mz + 1:mz] = diag(mz)*Q[i,j]
    }
  }
  ## This code assumes that #kids is not part of the state definition,
  ## so Fbullet = I
  Fbullet = diag(bigmz)

  ## Sanity check
  if (debugging)
    cat (sum(Qbullet %*% Gbullet %*% Sbullet - M), "should = 0.\n")

  ## create matrices for extended state space, which I will denote with
  ## the prefix "es".

  esbigmz = 4*bigmz
  esF = esM = matrix (0, esbigmz, esbigmz)

  esM[bigmz + 1:bigmz, 1:bigmz] = Fbullet
  esM[2*bigmz + 1:bigmz, bigmz + 1:bigmz] = Sbullet
  esM[3*bigmz + 1:bigmz, 2*bigmz + 1:bigmz] = Gbullet
  esM[1:bigmz, 3*bigmz + 1:bigmz] = Qbullet
  
  esF[bigmz + 1:bigmz, 1:bigmz] = F

  ################################################################
  ## Extended state model that includes ur-stage alpha_z and
  ## ur-environment alpha_q, as discussed in "Determining the
  ## contribution of traits."  Used for calculating contributions from
  ## birth stage and birth environment.  The "bs" before variable names
  ## stands for "birth state."
  ################################################################

  bsbigmz = 1 + mz + bigmz
  ## initial state
  bsM0 = rep(0, bsbigmz)
  bsM0[1] = 1

  bsF = bsM = matrix (0, bsbigmz, bsbigmz)
  ## From ur-stage to birth stage and ur-environment
  bsM[1 + (1:mz),1] = c0
  ## From birth stage and ur-environment to m0
  for (q in 1:numEnv) {
    bsM[1 + q*mz + 1:mz, 1 + 1:mz] = diag (u0[q], mz)
  }
  ## From normal stage x env. state to normal stage x env. state
  bsM[1 + mz + 1:bigmz, 1 + mz + 1:bigmz] = M
  ## You only start reproducing once you're past the ur-stages.  I guess
  ## reproduction should go to the ur-stage x ur-environment state.  Not
  ## that it matters much, since all we need are the column sums of bsF.
  bsF[1, 1 + mz + 1:bigmz] = colSums (F)

  ################################################################
  ## set up reward matrices
  ################################################################
  
  ## If the reward matrices aren't specified, we assume
  ## Poisson-distributed clutch sizes
  if (is.null(esR1)) {
    esR1 = esR2 = esR3 = matrix (0, esbigmz+1, esbigmz+1)
    bsR1 = bsR2 = bsR3 = matrix (0, bsbigmz+1, bsbigmz+1)

    ## First moment of clutch size (Poisson)
    for (j in 1:esbigmz) 
      esR1[,j] = sum(esF[,j])
    for (j in 1:bsbigmz) 
      bsR1[,j] = sum(bsF[,j])

    ## Second moment of clutch size  
    esR2 = esR1 + esR1^2
    bsR2 = bsR1 + bsR1^2
    ## Third moment of clutch size
    esR3 = esR1+ 3*esR1*esR1 + esR1*esR1*esR1
    bsR3 = bsR1+ 3*bsR1*bsR1 + bsR1*bsR1*bsR1
  }

  out = calcMoments (esM, esR1, esR2, esR3)
  esRho1Vec = out$rho1Vec
  esMu2Vec = out$mu2Vec
  esSkewnessVec = out$skewnessVec

  out = calcMoments (bsM, bsR1, bsR2, bsR3)
  bsRho1Vec = out$rho1Vec
  bsMu2Vec = out$mu2Vec
  bsMu3Vec = out$mu3Vec
  bsSkewnessVec = out$skewnessVec

  ## Start calculating luck terms
  survTrajecSkewness = growthTrajecSkewness = envTrajecSkewness =
    fecSkewness = numeric (maxAge)
  fecUpdateSkewness = survUpdateSkewness =
    growthUpdateSkewness = envUpdateSkewness = numeric (maxAge)
  survUpdateVar = growthUpdateVar = envUpdateVar = fecUpdateVar =
    numeric (maxAge) 
  survTrajecVar = growthTrajecVar = envTrajecVar = fecVar =
    numeric (maxAge)
  surv = numeric (maxAge)

  ## for ur-state var.
  ## expectation wrt z of Var(success | z)
  expZVarCondZ = bsMu2Vec %*% bsM0
  ## variance wrt z of Exp(success | z)
  varZExpCondZ = sum(bsM0*bsRho1Vec^2) - (sum(bsM0*bsRho1Vec)^2)
  ## Law of total variance
  urVar = expZVarCondZ + varZExpCondZ
  
  justBirthStageVar = bsMu2Vec %*% bsM %*% bsM0
  birthStageEnvVar = bsMu2Vec %*% bsM %*% bsM %*% bsM0

  ## eq. S8, but for variance instead of skewness
  birthStateVar = urVar - justBirthStageVar
  ## eq. S10, but for variance instead of skewness
  birthEnvVar = justBirthStageVar - birthStageEnvVar

  ## for ur-state skewness
  expZMu3CondZ = bsMu3Vec %*% bsM0
  ## 3rd moment wrt z of Exp(success | z)
  rho3ZExpCondZ = sum(bsM0*(bsRho1Vec)^3)
  ## 2nd moment wrt z of Exp(success | z)
  rho2ZExpCondZ = sum(bsM0*(bsRho1Vec)^2)
  ## 1st moment wrt z of Exp(success | z)
  expZExpCondZ = sum(bsM0*bsRho1Vec)
  ## 3rd central moment wrt z of Exp(success | z)
  mu3ZExpCondZ = rho3ZExpCondZ -
    3*expZExpCondZ*rho2ZExpCondZ +
    2*expZExpCondZ^3
  ## covariance wrt z of Exp(success | z) and Var(success | z)
  covZExpVarCondZ = sum(bsM0*bsRho1Vec*bsMu2Vec) -
    sum(bsM0*bsRho1Vec)*sum(bsM0*bsMu2Vec)
  ## Law of total cumulance
  urMu3 = expZMu3CondZ + mu3ZExpCondZ +
    3*covZExpVarCondZ
  urSkewness = urMu3 / urVar^1.5
  
  justBirthStageSkewness = bsSkewnessVec %*% bsM %*% bsM0
  birthStageEnvSkewness = bsSkewnessVec %*% bsM %*% bsM %*% bsM0

  ## eq. S8
  birthStateSkewness = urSkewness - justBirthStageSkewness
  ## eq. S10
  birthEnvSkewness = justBirthStageSkewness - birthStageEnvSkewness

  ## Starting the age loop
  MaM0 = m0
  bigmzZero = rep(0, bigmz)

  ## Sanity check --- passes
  if (debugging) {
    N = solve(diag(bigmz) - M)
    rho1Vec = colSums (F %*% N)
    cat ("Checking Exp(R):", esRho1Vec %*%
                             c(bigmzZero, bigmzZero, bigmzZero, m0),
         "should = ", rho1Vec %*% m0, "\n")
  }

  ## eq. 9
  fecUpdateSkewness[1] = esSkewnessVec %*%
    c(bigmzZero, Fbullet %*% MaM0, bigmzZero, bigmzZero)
  ## eq. 10
  survUpdateSkewness[1] = esSkewnessVec %*% 
    c(bigmzZero, bigmzZero, Sbullet %*% Fbullet %*% MaM0, bigmzZero)
  ## eq. 11
  growthUpdateSkewness[1] = esSkewnessVec %*% 
    c(bigmzZero, bigmzZero, bigmzZero, Pbullet %*% MaM0)
  ## eq. 8
  envUpdateSkewness[1] = esSkewnessVec %*%
    c(M %*% MaM0, bigmzZero, bigmzZero, bigmzZero)

  ## eq. 9 but for variance, not skewness
  fecUpdateVar[1] = esMu2Vec %*%
    c(bigmzZero, Fbullet %*% MaM0, bigmzZero, bigmzZero)
  ## eq. 10 but for variance, not skewness
  survUpdateVar[1] = esMu2Vec %*%
    c(bigmzZero, bigmzZero, Sbullet %*% MaM0, bigmzZero)
  ## eq. 11 but for variance, not skewness
  growthUpdateVar[1] = esMu2Vec %*%
    c(bigmzZero, bigmzZero, bigmzZero, Pbullet %*% MaM0)
  ## eq. 8 but for variance, not skewness
  envUpdateVar[1] = esMu2Vec %*%
    c(M %*% MaM0, bigmzZero, bigmzZero, bigmzZero)

  surv[1] = sum (MaM0)

  for (a in 2:maxAge) {
    MaM0 = M %*% MaM0

    ## eq. 9 
    fecUpdateSkewness[a] = esSkewnessVec %*%
      c(bigmzZero, Fbullet %*% MaM0, bigmzZero, bigmzZero)
    ## eq. 10
    survUpdateSkewness[a] = esSkewnessVec %*% 
      c(bigmzZero, bigmzZero, Sbullet %*% Fbullet %*% MaM0, bigmzZero)
    ## eq. 11
    growthUpdateSkewness[a] = esSkewnessVec %*% 
      c(bigmzZero, bigmzZero, bigmzZero, Pbullet %*% MaM0)
    ## eq. 8
    envUpdateSkewness[a] = esSkewnessVec %*%
      c(M %*% MaM0, bigmzZero, bigmzZero, bigmzZero)

    ## eq. 9 but for variance, not skewness
    fecUpdateVar[a] = esMu2Vec %*%
      c(bigmzZero, Fbullet %*% MaM0, bigmzZero, bigmzZero)
    ## eq. 10 but for variance, not skewness
    survUpdateVar[a] = esMu2Vec %*%
      c(bigmzZero, bigmzZero, Sbullet %*% MaM0, bigmzZero)
    ## eq. 11 but for variance, not skewness
    growthUpdateVar[a] = esMu2Vec %*%
      c(bigmzZero, bigmzZero, bigmzZero, Pbullet %*% MaM0)
    ## eq. 12 but for variance, not skewness
    envUpdateVar[a] = esMu2Vec %*%
      c(M %*% MaM0, bigmzZero, bigmzZero, bigmzZero)

    ## The following luck definitions are discussed just after eq. 11
    ## in the paper.
    
    ## fecundity skewness and var. 
    fecSkewness[a] = envUpdateSkewness[a-1] - fecUpdateSkewness[a]
    fecVar[a] = envUpdateVar[a-1] - fecUpdateVar[a]

    ## survival trajectory skewness and var.
    survTrajecSkewness[a-1] = fecUpdateSkewness[a-1] -
      survUpdateSkewness[a-1]
    survTrajecVar[a-1] = fecUpdateVar[a-1] - survUpdateVar[a-1]

    ## growth trajectory skewness and var.
    growthTrajecSkewness[a-1] = survUpdateSkewness[a-1] -
      growthUpdateSkewness[a-1] 
    growthTrajecVar[a-1] = survUpdateVar[a-1] - growthUpdateVar[a-1]
    
    ## environment trajectory skewness
    envTrajecSkewness[a-1] = growthUpdateSkewness[a-1] -
      envUpdateSkewness[a-1]
    envTrajecVar[a-1] = growthUpdateVar[a-1] - envUpdateVar[a-1]

    surv[a] = sum (MaM0)
  }

  ## And get the first value of fecSkewness and fecVar
  fecSkewness[1] = birthStageEnvSkewness - fecUpdateSkewness[1]
  fecVar[1] = birthStageEnvVar - fecUpdateVar[1]

  ## By what age are 99% of individuals dead?
  lifespan = which(surv < 0.01)[1]

  totSurvTrajecSkewness = sum(survTrajecSkewness)
  totGrowthTrajecSkewness = sum(growthTrajecSkewness)
  totEnvTrajecSkewness = sum(envTrajecSkewness)
  totFecSkewness = sum(fecSkewness)
  totSkewness = bsSkewnessVec %*% bsM0

  ## sanity check
  if (debugging) 
    cat ("Checking sum of skewness luck:",
         totSurvTrajecSkewness + totGrowthTrajecSkewness +
         totEnvTrajecSkewness + totFecSkewness +
         birthStateSkewness + birthEnvSkewness, "should = ",
         totSkewness, "\n")
  
  totSurvTrajecVar = sum(survTrajecVar)
  totGrowthTrajecVar = sum(growthTrajecVar)
  totEnvTrajecVar = sum(envTrajecVar)
  totFecVar = sum(fecVar)
  totVar = bsMu2Vec %*% bsM0
  ## Sanity check: passes
  if (debugging)
    cat ("Checking sum of variance luck:",
         totSurvTrajecVar +
         totGrowthTrajecVar +
         totEnvTrajecVar + totFecVar +
         birthStateVar + birthEnvVar, "should = ",
         totVar, "\n")

  return (out=list(birthStateVar=birthStateVar,
                   birthEnvVar=birthEnvVar,
                   survTrajecVar=survTrajecVar,
                   growthTrajecVar=growthTrajecVar,
                   envTrajecVar=envTrajecVar,
                   fecVar=fecVar,
                   totVar=totVar,
                   birthStateSkewness=birthStateSkewness,
                   birthEnvSkewness=birthEnvSkewness,
                   survTrajecSkewness=survTrajecSkewness,
                   growthTrajecSkewness=growthTrajecSkewness,
                   envTrajecSkewness=envTrajecSkewness,
                   fecSkewness=fecSkewness,
                   totSkewness=totSkewness,
                   lifespan=lifespan))
}

## This function is almost identical to
## getVarSkewnessPartitionsEnvVar, but does the partitioning of the
## 3rd central moment of LRO, not the skewness.
getVarMu3PartitionsEnvVar = function (Plist, Flist, Q, m0,
                                           maxAge=100, esR1=NULL,
                                           esR2=NULL, esR3=NULL,
                                           bsR1=NULL, bsR2=NULL,
                                           bsR3=NULL)
{
  require (Matrix)
  debugging = TRUE
  
  mz = dim(Plist[[1]])[1]
  numEnv = dim(Q)[1]
  bigmz = numEnv*mz

  ## Define megamatrices M and F
  F = M = matrix (0, bigmz, bigmz)
  for (i in 1:numEnv) {
    for (j in 1:numEnv) {
      M[(i-1)*mz + 1:mz, (j-1)*mz + 1:mz] = Plist[[j]]*Q[i,j]
      F[(i-1)*mz + 1:mz, (j-1)*mz + 1:mz] = Flist[[j]]*Q[i,j]
    }
  }

  ## Make survival and growth matrices
  Slist = Glist = list (numEnv)
  for (q in 1:numEnv) {
    Slist[[q]] = diag (colSums (Plist[[q]]))
    Sinv = diag (1 / diag(Slist[[q]]))
    Glist[[q]] = Plist[[q]] %*% Sinv
  }

  ## Make "bullet" matrices.  bdiag makes sparse matrices, which can't
  ## be pasted into regular matrices, so wrap these in as.matrix().
  Sbullet = as.matrix(bdiag (Slist))
  Gbullet = as.matrix(bdiag (Glist))
  Pbullet = Gbullet %*% Sbullet
  Qbullet = matrix (0, bigmz, bigmz)
  for (i in 1:numEnv) {
    for (j in 1:numEnv) {
      Qbullet[(i-1)*mz + 1:mz, (j-1)*mz + 1:mz] = diag(mz)*Q[i,j]
    }
  }
  ## This code assumes that #kids is not part of the state definition,
  ## so Fbullet = I
  Fbullet = diag(bigmz)

  ## Sanity check
  if (debugging)
    cat (sum(Qbullet %*% Gbullet %*% Sbullet - M), "should = 0.\n")

  ## create matrices for extended state space, which I will denote with
  ## the prefix "es".

  esbigmz = 4*bigmz
  esF = esM = matrix (0, esbigmz, esbigmz)

  esM[bigmz + 1:bigmz, 1:bigmz] = Fbullet
  esM[2*bigmz + 1:bigmz, bigmz + 1:bigmz] = Sbullet
  esM[3*bigmz + 1:bigmz, 2*bigmz + 1:bigmz] = Gbullet
  esM[1:bigmz, 3*bigmz + 1:bigmz] = Qbullet
  
  esF[bigmz + 1:bigmz, 1:bigmz] = F

  ################################################################
  ## Extended state model that includes ur-stage alpha_z and
  ## ur-environment alpha_q.  Used for calculating contributions from
  ## birth stage and birth environment.  The "bs" before variable names
  ## stands for "birth state."
  ################################################################

  bsbigmz = 1 + mz + bigmz
  ## initial state
  bsM0 = rep(0, bsbigmz)
  bsM0[1] = 1

  bsF = bsM = matrix (0, bsbigmz, bsbigmz)
  ## From ur-stage to birth stage and ur-environment
  bsM[1 + (1:mz),1] = c0
  ## From birth stage and ur-environment to m0
  for (q in 1:numEnv) {
    bsM[1 + q*mz + 1:mz, 1 + 1:mz] = diag (u0[q], mz)
  }
  ## From normal stage x env. state to normal stage x env. state
  bsM[1 + mz + 1:bigmz, 1 + mz + 1:bigmz] = M
  ## You only start reproducing once you're past the ur-stages.  I guess
  ## reproduction should go to the ur-stage x ur-environment state.  Not
  ## that it matters much, since all we need are the column sums of bsF.
  bsF[1, 1 + mz + 1:bigmz] = colSums (F)

  ################################################################
  ## set up reward matrices
  ################################################################
  
  ## If the reward matrices aren't specified, we assume
  ## Poisson-distributed clutch sizes
  if (is.null(esR1)) {
    esR1 = esR2 = esR3 = matrix (0, esbigmz+1, esbigmz+1)
    bsR1 = bsR2 = bsR3 = matrix (0, bsbigmz+1, bsbigmz+1)

    ## First moment of clutch size (Poisson)
    for (j in 1:esbigmz) 
      esR1[,j] = sum(esF[,j])
    for (j in 1:bsbigmz) 
      bsR1[,j] = sum(bsF[,j])

    ## Second moment of clutch size  
    esR2 = esR1 + esR1^2
    bsR2 = bsR1 + bsR1^2
    ## Third moment of clutch size
    esR3 = esR1+ 3*esR1*esR1 + esR1*esR1*esR1
    bsR3 = bsR1+ 3*bsR1*bsR1 + bsR1*bsR1*bsR1
  }

  out = calcMoments (esM, esR1, esR2, esR3)
  esRho1Vec = out$rho1Vec
  esMu2Vec = out$mu2Vec
  esMu3Vec = out$mu3Vec

  out = calcMoments (bsM, bsR1, bsR2, bsR3)
  bsRho1Vec = out$rho1Vec
  bsMu2Vec = out$mu2Vec
  bsMu3Vec = out$mu3Vec

  ## Start calculating luck terms
  survTrajecMu3 = growthTrajecMu3 = envTrajecMu3 =
    fecMu3 = numeric (maxAge)
  fecUpdateMu3 = survUpdateMu3 =
    growthUpdateMu3 = envUpdateMu3 = numeric (maxAge)
  survUpdateVar = growthUpdateVar = envUpdateVar = fecUpdateVar =
    numeric (maxAge) 
  survTrajecVar = growthTrajecVar = envTrajecVar = fecVar =
    numeric (maxAge)
  surv = numeric (maxAge)

  ## for ur-state var.
  ## expectation wrt z of Var(success | z)
  expZVarCondZ = bsMu2Vec %*% bsM0
  ## variance wrt z of Exp(success | z)
  varZExpCondZ = sum(bsM0*bsRho1Vec^2) - (sum(bsM0*bsRho1Vec)^2)
  ## Law of total variance
  urVar = expZVarCondZ + varZExpCondZ
  
  justBirthStageVar = bsMu2Vec %*% bsM %*% bsM0
  birthStageEnvVar = bsMu2Vec %*% bsM %*% bsM %*% bsM0
  
  birthStateVar = urVar - justBirthStageVar
  birthEnvVar = justBirthStageVar - birthStageEnvVar

  ## for ur-state skewness
  expZMu3CondZ = bsMu3Vec %*% bsM0
  ## 3rd moment wrt z of Exp(success | z)
  rho3ZExpCondZ = sum(bsM0*(bsRho1Vec)^3)
  ## 2nd moment wrt z of Exp(success | z)
  rho2ZExpCondZ = sum(bsM0*(bsRho1Vec)^2)
  ## 1st moment wrt z of Exp(success | z)
  expZExpCondZ = sum(bsM0*bsRho1Vec)
  ## 3rd central moment wrt z of Exp(success | z)
  mu3ZExpCondZ = rho3ZExpCondZ -
    3*expZExpCondZ*rho2ZExpCondZ +
    2*expZExpCondZ^3
  ## covariance wrt z of Exp(success | z) and Var(success | z)
  covZExpVarCondZ = sum(bsM0*bsRho1Vec*bsMu2Vec) -
    sum(bsM0*bsRho1Vec)*sum(bsM0*bsMu2Vec)
  ## Law of total cumulance
  urMu3 = expZMu3CondZ + mu3ZExpCondZ +
    3*covZExpVarCondZ
  
  justBirthStageMu3 = bsMu3Vec %*% bsM %*% bsM0
  birthStageEnvMu3 = bsMu3Vec %*% bsM %*% bsM %*% bsM0

  birthStateMu3 = urMu3 - justBirthStageMu3
  birthEnvMu3 = justBirthStageMu3 - birthStageEnvMu3

  ## Starting the age loop
  MaM0 = m0
  bigmzZero = rep(0, bigmz)

  ## Sanity check --- passes
  if (debugging) {
    N = solve(diag(bigmz) - M)
    rho1Vec = colSums (F %*% N)
    cat ("Checking Exp(R):", esRho1Vec %*%
                             c(bigmzZero, bigmzZero, bigmzZero, m0),
         "should = ", rho1Vec %*% m0, "\n")
  }

  fecUpdateMu3[1] = esMu3Vec %*%
    c(bigmzZero, Fbullet %*% MaM0, bigmzZero, bigmzZero)
  survUpdateMu3[1] = esMu3Vec %*% 
    c(bigmzZero, bigmzZero, Sbullet %*% Fbullet %*% MaM0, bigmzZero)
  growthUpdateMu3[1] = esMu3Vec %*% 
    c(bigmzZero, bigmzZero, bigmzZero, Pbullet %*% MaM0)
  envUpdateMu3[1] = esMu3Vec %*%
    c(M %*% MaM0, bigmzZero, bigmzZero, bigmzZero)

  fecUpdateVar[1] = esMu2Vec %*%
    c(bigmzZero, Fbullet %*% MaM0, bigmzZero, bigmzZero)
  survUpdateVar[1] = esMu2Vec %*%
    c(bigmzZero, bigmzZero, Sbullet %*% MaM0, bigmzZero) 
  growthUpdateVar[1] = esMu2Vec %*%
    c(bigmzZero, bigmzZero, bigmzZero, Pbullet %*% MaM0)
  envUpdateVar[1] = esMu2Vec %*%
    c(M %*% MaM0, bigmzZero, bigmzZero, bigmzZero)

  surv[1] = sum (MaM0)

  for (a in 2:maxAge) {
    MaM0 = M %*% MaM0

    fecUpdateMu3[a] = esMu3Vec %*%
      c(bigmzZero, Fbullet %*% MaM0, bigmzZero, bigmzZero)
    survUpdateMu3[a] = esMu3Vec %*% 
      c(bigmzZero, bigmzZero, Sbullet %*% Fbullet %*% MaM0, bigmzZero)
    growthUpdateMu3[a] = esMu3Vec %*% 
      c(bigmzZero, bigmzZero, bigmzZero, Pbullet %*% MaM0)
    envUpdateMu3[a] = esMu3Vec %*%
      c(M %*% MaM0, bigmzZero, bigmzZero, bigmzZero)

    fecUpdateVar[a] = esMu2Vec %*%
      c(bigmzZero, Fbullet %*% MaM0, bigmzZero, bigmzZero)
    survUpdateVar[a] = esMu2Vec %*%
      c(bigmzZero, bigmzZero, Sbullet %*% MaM0, bigmzZero) 
    growthUpdateVar[a] = esMu2Vec %*%
      c(bigmzZero, bigmzZero, bigmzZero, Pbullet %*% MaM0)
    envUpdateVar[a] = esMu2Vec %*%
      c(M %*% MaM0, bigmzZero, bigmzZero, bigmzZero)

    ## fecundity skewness and var.
    fecMu3[a] = envUpdateMu3[a-1] - fecUpdateMu3[a]
    fecVar[a] = envUpdateVar[a-1] - fecUpdateVar[a]

    ## survival trajectory skewness and var.
    survTrajecMu3[a-1] = fecUpdateMu3[a-1] -
      survUpdateMu3[a-1]
    survTrajecVar[a-1] = fecUpdateVar[a-1] - survUpdateVar[a-1]

    ## growth trajectory skewness and var.
    growthTrajecMu3[a-1] = survUpdateMu3[a-1] -
      growthUpdateMu3[a-1] 
    growthTrajecVar[a-1] = survUpdateVar[a-1] - growthUpdateVar[a-1]
    
    ## environment trajectory skewness
    envTrajecMu3[a-1] = growthUpdateMu3[a-1] -
      envUpdateMu3[a-1]
    envTrajecVar[a-1] = growthUpdateVar[a-1] - envUpdateVar[a-1]

    surv[a] = sum (MaM0)
  }

  ## And get the first value of fecMu3 and fecVar
  fecMu3[1] = birthStageEnvMu3 - fecUpdateMu3[1]
  fecVar[1] = birthStageEnvVar - fecUpdateVar[1]

  ## By what age are 99% of individuals dead?
  lifespan = which(surv < 0.01)[1]

  totSurvTrajecMu3 = sum(survTrajecMu3)
  totGrowthTrajecMu3 = sum(growthTrajecMu3)
  totEnvTrajecMu3 = sum(envTrajecMu3)
  totFecMu3 = sum(fecMu3)
  totMu3 = bsMu3Vec %*% bsM0

  ## sanity check
  if (debugging) ## Passes
    cat ("Checking sum of Mu3 luck:",
         totSurvTrajecMu3 + totGrowthTrajecMu3 +
         totEnvTrajecMu3 + totFecMu3 +
         birthStateMu3 + birthEnvMu3, "should = ",
         totMu3, "\n")
  
  totSurvTrajecVar = sum(survTrajecVar)
  totGrowthTrajecVar = sum(growthTrajecVar)
  totEnvTrajecVar = sum(envTrajecVar)
  totFecVar = sum(fecVar)
  totVar = bsMu2Vec %*% bsM0
  ## Sanity check: passes
  if (debugging)
    cat ("Checking sum of variance luck:",
         totSurvTrajecVar +
         totGrowthTrajecVar +
         totEnvTrajecVar + totFecVar +
         birthStateVar + birthEnvVar, "should = ",
         totVar, "\n")


  return (out=list(birthStateVar=birthStateVar,
                   birthEnvVar=birthEnvVar,
                   survTrajecVar=survTrajecVar,
                   growthTrajecVar=growthTrajecVar,
                   envTrajecVar=envTrajecVar,
                   fecVar=fecVar,
                   totVar=totVar,
                   birthStateMu3=birthStateMu3,
                   birthEnvMu3=birthEnvMu3,
                   survTrajecMu3=survTrajecMu3,
                   growthTrajecMu3=growthTrajecMu3,
                   envTrajecMu3=envTrajecMu3,
                   fecMu3=fecMu3,
                   totMu3=totMu3,
                   lifespan=lifespan))
}

#####################################################################
## Unlike distLifespanCondR, this version doesn't require you to add
## age as a state variable.  Keeps the memory needs to a dull roar.
##
## This version uses the kernel conditional on LRO = R_T, not LRO >=
## R_T.  (That version is in genericPartitionVarSkewness4.R and gives
## the same answers.)
##
#####################################################################

distLifespanCondR2 = function (Plist, Flist, Q,
                                  m0, maxKids, maxAge,
                                  percentileCutoff = 0.99,
                                  B=NULL, Fdist="Poisson") {
  require (Matrix)
  debugging = TRUE
  
  mT = maxKids + 1
  mA = maxAge + 1
  
  mz = dim(Plist[[1]])[1]
  numEnv = dim(Q)[1]
  bigmz = numEnv*mz

  ## Define megamatrices M and F
  F = M = matrix (0, bigmz, bigmz)
  for (i in 1:numEnv) {
    for (j in 1:numEnv) {
      M[(i-1)*mz + 1:mz, (j-1)*mz + 1:mz] = Plist[[j]]*Q[i,j]
      F[(i-1)*mz + 1:mz, (j-1)*mz + 1:mz] = Flist[[j]]*Q[i,j]
    }
  }

  if (is.null(B)) 
    ## B[i,j] is the probability that a class-j individual has i-1 kids.
    ## We assume Poisson-distributed number of offspring.
    ## The columns of B should sum to 1 and they do.
    B = mk_B (maxKids, F)

  ## Construct A, the transition matrix for a (number of kids) x stage x
  ## env. model.   
  out = make_AxT (B, M, mT)
  A = out$A
  mzA = bigmz*mT
  K = out$K

  #####################################################################
  ## making the bullet matrices
  #####################################################################

  ## Make "bullet" matrices for A, as discussed in "Age-partitioning
  ## of skewness contributions from different luck types"
  Fbullet = Qbullet = Gbullet = matrix (0, mzA, mzA)

  ## Sbullet updates survival
  Sbullet = diag (colSums(A))

  ## Fbullet updates number of kids.
  for (j in 1:mT) {
    for (i in j:(mT-1)) {
      for (z in 1:bigmz) {
        if (Fdist == "Poisson") {
          Fbullet[(i-1)*bigmz + z, (j-1)*bigmz + z] =
            dpois (i-j, lambda=sum(F[,z]))
        } else if (Fdist == "Bernoulli") {
          Fbullet[(i-1)*bigmz + z, (j-1)*bigmz + z] =
            dbinom (i-j, prob=sum(F[,z]), size=1)
        } else {
          cat ("Did not recognize Fdist choice.\n")
        }
      }
    }
  }
  ## Make mT absorbing.  If you have #kids index < mT, the probability
  ## of going to mT while in stage z = 1 - all other kid transition
  ## probabilities with stage z.

  for (j in 1:(mT-1)) {
    for (z in 1:bigmz) {
      Fbullet[(mT-1)*bigmz + z, (j-1)*bigmz + z] =
        ##      1 - sum(Fbullet[1:((mT-1)*bigmz), (j-1)*bigmz + z])
        1 - sum(Fbullet[(0:(mT-2))*bigmz + z, (j-1)*bigmz + z])
    }
  }
  ## If you have #kids index = mT, you stay there.
  for (z in 1:bigmz) 
    Fbullet[(mT-1)*bigmz + z, (mT-1)*bigmz + z] = 1

  ## Qbullet updates env.
  smallQbullet = matrix (0, bigmz, bigmz)
  for (i in 1:numEnv) {
    for (j in 1:numEnv) {
      smallQbullet[(i-1)*mz + 1:mz, (j-1)*mz + 1:mz] = diag(mz)*Q[i,j]
    }
  }
  ## Qbullet is the same for each number of kids
  for (k in 1:mT) 
    Qbullet[(k-1)*bigmz + 1:bigmz, (k-1)*bigmz + 1:bigmz] = smallQbullet

  ## Gbullet updates stage
  smallGbullet = matrix (0, bigmz, bigmz)
  for (q in 1:numEnv) {
    S = diag (colSums (Plist[[q]]))
    Sinv = diag (1 / diag(S))
    G = Plist[[q]] %*% Sinv
    smallGbullet[(q-1)*mz + 1:mz, (q-1)*mz + 1:mz] = G
  }
  for (k in 1:mT) 
    Gbullet[(k-1)*bigmz + 1:bigmz, (k-1)*bigmz + 1:bigmz] = smallGbullet

  ## sanity check
  if (debugging)
    cat ("Checking the bullet matrices for all #kids:",
         range((Qbullet %*% Gbullet %*% Sbullet %*% Fbullet - A)),
         "should = 0.\n")


  ####################################################################
  ## Now make the additionally extended space matrix that allows us to
  ## stop partway through a time step.  Our new prefix is es to
  ## indicate this additional extension.  ("es" = "extended space")
  ####################################################################
  
  esmzA = 4*mzA

  ##esbigF = esA = matrix (0, esmzA, esmzA)
  esA = matrix (0, esmzA, esmzA)
  esA[mzA + 1:mzA, 1:mzA] = Fbullet
  esA[2*mzA + 1:mzA, mzA + 1:mzA] = Sbullet
  esA[3*mzA + 1:mzA, 2*mzA + 1:mzA] = Gbullet
  esA[1:mzA, 3*mzA + 1:mzA] = Qbullet

  ##################################################################
  ## What is the distribution of #kids at death when reproduction
  ## comes before survival and growth?  If you're unfamiliar with
  ## this calculation, you can find it derived in section 3.2.1,
  ## "Mortality: age and size at death" of "Data-driven modelling of
  ## structured populations," by Ellner, Childs, and Rees, Springer,
  ## 2016.
  ##################################################################

  ## Create a survival probability vector 
  surv = apply (esA, 2, sum);  die = 1-surv; 

  ## And the fundamental matrix of esA
  fundesA <- solve(diag(ncol(esA))-esA)

  ## Make omega
  omega = matrix(0,nrow(fundesA),ncol(fundesA)); 

  for(j in 1:ncol(fundesA))
    omega[,j] = die*fundesA[,j]

  distAtBirth = matrix(0, bigmz, 4*mT)
  distAtBirth[,1] = m0  ## Everyone starts off with zero kids at the
  ## beginning of the time step

  ## Get distribution of states at death.  
  distAtDeath = omega %*% matrix(distAtBirth, ncol=1)  # state vector
  ## marginalize over stages to get just the distribution of #kids at
  ## death
  distAtDeath = array (distAtDeath, dim=c(bigmz, mT, 4)) # state
                                       # matrix
  distKidsAtDeath = apply (distAtDeath, 2, sum)

  ## What threshold number of kids represents the 99th %ile of the LRO
  ## distribution? 
  cumDistKidsAtDeath = cumsum (distKidsAtDeath)
  R90 = which (cumDistKidsAtDeath > 0.9)[1]
  R99 = which (cumDistKidsAtDeath > 0.99)[1]
  if (debugging) {
    cat ("The 90th %ile of the LRO distribution occurs at", R90,
         "kids.\n")
    cat ("The 99th %ile of the LRO distribution occurs at", R99,
         "kids.\n")
  }
  RCutoff = which (cumDistKidsAtDeath > percentileCutoff)[1]
  if (RCutoff > maxKids)
    stop ("RCutoff = ", RCutoff, "but maxKids = ", maxKids, "\n")

  probLifespanCondR = matrix (0, RCutoff+2, mA)
  probThresholdOrMore = matrix (0, RCutoff+2, esmzA)
  probSurvAgeA = numeric (mA+1)
  probSuccessCondZ = matrix (0, RCutoff+1, esmzA)
  probSuccess = numeric (RCutoff+1)
  jointProbLifespanR = probLifespanCondR = matrix (0, RCutoff+1, mA)
  normalSurvProb = numeric (RCutoff+2)

  AaM0 = c(m0, rep(0, (mT-1)*bigmz))
  mzAZero = rep(0, mzA)
  extendedInit = c(AaM0, rep(mzAZero, 3))

  ## The case of R = 0 (kidsIndexThreshold = 1) #########
  probThresholdOrMore[1,] = 1
  esAa = diag (esmzA)
  esA4 = esA %*% esA %*% esA %*% esA
  for (a in 1:(mA+1)) {
    cat ("a = ", a, "\n")
    ## Probability of surviving to age a (eq. 18)
    probSurvAgeA[a] = colSums (esAa) %*% extendedInit
    if (a >= 2)
      ## Probability of having lifespan = a, conditional on R = 0
      ## (eq. 19)
      probLifespanCondR[1,a-1] =
        probSurvAgeA[a-1] - probSurvAgeA[a]
    ## Keep track of survival probabilities for unconditional kernel.
    ## we need to hit the last matrix 4 times to advance one time step
    esAa = esA4 %*% esAa
  }
  ## survival probability without conditioning on having R > 0
  normalSurvProb = probSurvAgeA

  ## The case of R > 0
  ##for (kidsIndexThreshold in 2:(RCutoff+2)) {

  ## k is the actual number of kids for the threshold.  We start with
  ## the threshold for success at LRO = 1 kid.
  for (k in 1:(RCutoff+1)) {
    cat ("#kids threshold = ", k, "\n")

    ## Calculate transition matrix esA conditional on success (eq. 16)
    cutoff = k*bigmz
    transientStates = c(1:cutoff, mzA + 1:cutoff, 2*mzA + 1:cutoff,
                        3*mzA + 1:cutoff)
    out = makeCondKernel (esA, transientStates)
    ## probability that you met or exceeded the threshold number of
    ## kids 
    probThresholdOrMore[k+1,] = out$q2Extended

    ## probability that you had precisely k kids (eq. 15)
    probSuccessCondZ[k,] =
      probThresholdOrMore[k,] - probThresholdOrMore[k+1,]
    probSuccess[k] = probSuccessCondZ[k,] %*% extendedInit

    if (debugging)
      cat ("Pr(success) = ", probSuccess[k], "\n")

    ## calculate extended transition matrix conditional on success
    ## (eq. 16) 
    condEsA = matrix (0, esmzA, esmzA)
    ## For any starting state with #kids > threshold, probSuccessCondZ
    ## is zero and I think condEsA should be zero too.  Only do the
    ## following for z s.t. probSuccessCondZ > 0.
    for (j in transientStates)
      condEsA[,j] = esA[,j] * probSuccessCondZ[k,] /
        probSuccessCondZ[k,j]

    ## Useful later
    condEsA4 = condEsA %*% condEsA %*% condEsA %*% condEsA

    ## Initial distribution conditional on having LRO = exactly
    ## kidsIndexThreshold-1 (eq. 17)
    initCondSuccess =
      probSuccessCondZ[k,] * extendedInit
    initCondSuccess = initCondSuccess / sum(initCondSuccess)

    ## What is the lifespan dist. given that you had at least
    ## numKidsThreshold kids?
    condEsAa = diag(esmzA)
    for (a in 1:(mA+1)) {
      cat ("a = ", a, "\n")
      ## eq. 18
      probSurvAgeA[a] = colSums (condEsAa) %*% initCondSuccess
      ## Keep track of survival probabilities for unconditional kernel.
      ## we need to hit the last matrix 4 times to advance one time step
      condEsAa = condEsA4 %*% condEsAa
    }

    ## eq. 19
    probLifespanCondR[k,] = -diff (probSurvAgeA)
  }  ## end loop over k

  ## Sanity check: passes if maxAge is large enough
  if (debugging)
    cat ("sum_x Pr(lifespan = x | R) is ",
         apply (probLifespanCondR, 1, sum), "should = 1s.\n")

  meanLifespanCondR = sdLifespanCondR = CVLifespanCondR =
    numeric (RCutoff+1)
  
  for (k in 1:(RCutoff+1)) {
    meanLifespanCondR[k] = sum((1:mA)*probLifespanCondR[k,])
    varLifespanCondR = sum((1:mA)^2*probLifespanCondR[k,]) -
      meanLifespanCondR[k]^2
    sdLifespanCondR[k] = sqrt(varLifespanCondR)
    CVLifespanCondR[k] = sdLifespanCondR[k] / meanLifespanCondR[k]
  }

  ## Now let's get Pr(R | Lifespan) (except that I've decided I don't
  ## want that after all, but let's keep it here, just in case I want
  ## it later.)

  ## Passed sanity check: I have compared the output below to
  ## calculating Pr(lifespan) from an age-structured model and they
  ## are identical.
  distLifespan = apply (jointProbLifespanR, 2, sum)

  probRCondLifespan = matrix (0, mA, RCutoff+1)
  for (a in 1:mA)
    probRCondLifespan[a,] = jointProbLifespanR[,a] / distLifespan[a]
  
  return (list(probLifespanCondR=probLifespanCondR,
               distKidsAtDeath=distKidsAtDeath,
               sdLifespanCondR=sdLifespanCondR,
               CVLifespanCondR=CVLifespanCondR,
               maxKidsIndex=RCutoff+1,
               normalSurvProb=normalSurvProb)) }


######## Code graveyard --- useful but ultimately unused ##########

distLifespanCondR = function (Plist, Flist, Q,
                              m0, maxAge, maxKids,
                              B=NULL, Fdist="Poisson") 
{
  require (Matrix)
  debugging = TRUE
  
  mT = maxKids + 1
  mA = maxAge + 1
  
  mz = dim(Plist[[1]])[1]
  numEnv = dim(Q)[1]
  bigmz = numEnv*mz

  ## Define megamatrices M and F
  F = M = matrix (0, bigmz, bigmz)
  for (i in 1:numEnv) {
    for (j in 1:numEnv) {
      M[(i-1)*mz + 1:mz, (j-1)*mz + 1:mz] = Plist[[j]]*Q[i,j]
      F[(i-1)*mz + 1:mz, (j-1)*mz + 1:mz] = Flist[[j]]*Q[i,j]
    }
  }

  ## expected number of offspring in a clutch
  ## b = colSums (F)

  if (is.null(B)) 
    ## B[i,j] is the probability that a class-j individual has i-1 kids.
    ## We assume Poisson-distributed number of offspring.
    ## The columns of B should sum to 1 and they do.
    B = mk_B (maxKids, F)
  
  ## Construct A, the transition matrix for a (number of kids) x stage x
  ## env. model.   
  out = make_AxT (B, M, mT)
  A = out$A
  mzA = bigmz*mT
  K = out$K

  #####################################################################
  ## making the bullet matrices
  #####################################################################

  ## Make "bullet" matrices for A.
  Fbullet = Qbullet = Gbullet = matrix (0, mzA, mzA)

  ## Sbullet updates survival
  Sbullet = diag (colSums(A))

  ## Fbullet updates number of kids.
  for (j in 1:mT) {
    for (i in j:(mT-1)) {
      for (z in 1:bigmz) {
        if (Fdist == "Poisson") {
          Fbullet[(i-1)*bigmz + z, (j-1)*bigmz + z] =
            dpois (i-j, lambda=sum(F[,z]))
        } else if (Fdist == "Bernoulli") {
          Fbullet[(i-1)*bigmz + z, (j-1)*bigmz + z] =
            dbinom (i-j, prob=sum(F[,z]), size=1)
        } else {
          cat ("Did not recognize Fdist choice.\n")
        }
      }
    }
  }
  ## Make mT absorbing.  If you have #kids index < mT, the probability
  ## of going to mT while in stage z = 1 - all other kid transition
  ## probabilities with stage z.

  for (j in 1:(mT-1)) {
    for (z in 1:bigmz) {
      Fbullet[(mT-1)*bigmz + z, (j-1)*bigmz + z] =
        ##      1 - sum(Fbullet[1:((mT-1)*bigmz), (j-1)*bigmz + z])
        1 - sum(Fbullet[(0:(mT-2))*bigmz + z, (j-1)*bigmz + z])
    }
  }
  ## If you have #kids index = mT, you stay there.
  for (z in 1:bigmz) 
    Fbullet[(mT-1)*bigmz + z, (mT-1)*bigmz + z] = 1

  ## Qbullet updates env.
  smallQbullet = matrix (0, bigmz, bigmz)
  for (i in 1:numEnv) {
    for (j in 1:numEnv) {
      smallQbullet[(i-1)*mz + 1:mz, (j-1)*mz + 1:mz] = diag(mz)*Q[i,j]
    }
  }
  ## Qbullet is the same for each number of kids
  for (k in 1:mT) 
    Qbullet[(k-1)*bigmz + 1:bigmz, (k-1)*bigmz + 1:bigmz] = smallQbullet

  ## Gbullet updates stage
  smallGbullet = matrix (0, bigmz, bigmz)
  for (q in 1:numEnv) {
    S = diag (colSums (Plist[[q]]))
    Sinv = diag (1 / diag(S))
    G = Plist[[q]] %*% Sinv
    smallGbullet[(q-1)*mz + 1:mz, (q-1)*mz + 1:mz] = G
  }
  for (k in 1:mT) 
    Gbullet[(k-1)*bigmz + 1:bigmz, (k-1)*bigmz + 1:bigmz] = smallGbullet

  ## sanity check
  if (debugging)
    cat ("Checking the bullet matrices for all #kids:",
         range((Qbullet %*% Gbullet %*% Sbullet %*% Fbullet - A)),
         "should = 0.\n")

  ## This line is time-consuming for large matrices
  Pbullet = Gbullet %*% Sbullet %*% Fbullet

  ####################################################################
  ## Now make the additionally extended space matrix that allows us to
  ## stop partway through a time step.  Our new prefix is es to
  ## indicate this additional extension.  ("es" = "extended space")
  ####################################################################

  esmzA = 4*mzA

  ##esbigF = esA = matrix (0, esmzA, esmzA)
  esA = matrix (0, esmzA, esmzA)
  esA[mzA + 1:mzA, 1:mzA] = Fbullet
  esA[2*mzA + 1:mzA, mzA + 1:mzA] = Sbullet
  esA[3*mzA + 1:mzA, 2*mzA + 1:mzA] = Gbullet
  esA[1:mzA, 3*mzA + 1:mzA] = Qbullet

  ## Now take the size x kids extended state transition matrix and
  ## expand it to include age as a state variable.

  bigesmzA = esmzA*mA
  bigEsA = matrix (0, bigesmzA, bigesmzA)
  for (a in 1:maxAge) {
    ## the reproduction, survival, and growth updates keep us within
    ## the same age.
    bigEsA[(a-1)*esmzA + 1:esmzA, (a-1)*esmzA + 1:(3*mzA)] =
      esA[1:esmzA, 1:(3*mzA)]
    ## the environment update takes us to the next age.
    bigEsA[a*esmzA + 1:esmzA, (a-1)*esmzA + 3*mzA + 1:mzA] =
      esA[1:esmzA, 3*mzA + 1:mzA]
  }
  ## Final age class is age maxAge and older
  bigEsA[maxAge*esmzA + 1:esmzA, maxAge*esmzA + 1:esmzA] = esA

##################################################################
  ## What is the joint distribution of age and #kids at death?
##################################################################

  ## Create a survival probability vector 
  surv = apply (bigEsA, 2, sum);  die = 1-surv; 

  cat ("Calculating the fundamental matrix...\n")
  ## And the fundamental matrix of A
  fundBigEsA <- solve(diag(ncol(bigEsA)) - bigEsA)

  ## Make omega
  omega = matrix (0, nrow(fundBigEsA), ncol(fundBigEsA)); 

  cat ("Calculating omega...\n")
  for(j in 1:ncol(fundBigEsA))
    omega[,j] = die*fundBigEsA[,j]

  distAtBirth = matrix(0, bigmz, 4*mT*mA)
  distAtBirth[,1] = m0  ## Everyone starts off with zero kids at age
  ## 0 at the beginning of the time step.

  ## Get distribution of states at death
  distAtDeathVec = omega %*% matrix(distAtBirth, ncol=1)  # state vector
  ##distAtDeath = array (distAtDeathVec, dim=c(bigmz, mT, mA, 4))
  distAtDeath = array (distAtDeathVec, dim=c(bigmz, mT, 4, mA))
  ## distKidsAtDeath is good.
  distKidsAtDeath = apply (distAtDeath, 2, sum)
  ## distAgeAtDeath is complete nonsense, presumably because I ordered
  ## the dimensions incorrectly for distAtDeath.
  distAgeAtDeath = apply (distAtDeath, 4, sum)
  distAgeKidsAtDeath = apply (distAtDeath, c(2, 4), sum)

  ## Sanity check: passes
  if (debugging) {
    cutoff = bigmz
    transientStates = c(1:cutoff, mzA + 1:cutoff, 2*mzA + 1:cutoff,
                        3*mzA + 1:cutoff)
    out = makeCondKernel (esA, transientStates)
    esQ2Extended = out$q2Extended
    AaM0 = c(m0, rep(0, (mT-1)*bigmz))
    mzAZero = rep(0, mzA)
    cat ("The probability of having at least one kid:", 
         1 - distKidsAtDeath[1], "should = ",
         esQ2Extended %*% c(AaM0, mzAZero, mzAZero, mzAZero), "\n")
  }

  ## Sanity check: slightly off?  Eviction problem?
  meanAgeAtDeath = sum((1:mA)*distAgeAtDeath)
  N = solve (diag(bigmz) - M)
  meanAgeAtDeath2 = colSums(N) %*% m0
  cat ("Checking mean lifespan:", meanAgeAtDeath, "should = ",
       meanAgeAtDeath2, "\n")

  ## Pr(Lifespan | R) = P(Lifespan, R) / P(R)
  probLifespanCondR = matrix (0, mT, mA)
  for (k in 1:mT)
    probLifespanCondR[k,] = distAgeKidsAtDeath[k,] / distKidsAtDeath[k]

  ## Pr(R | Lifespan) = P(Lifespan, R) / P(lifespan)
  probRCondLifespan = matrix (0, mA, mT)
  for (a in 1:mA)
    probRCondLifespan[a,] = distAgeKidsAtDeath[,a] / distAgeAtDeath[a]

  ## Calculate std. dev. of lifespan and CV of lifespan conditional on
  ## R.
  sdLifespanCondR = CVLifespanCondR = numeric (mT)
  for (k in 1:mT) {
    meanLifespanCondR = sum((1:mA)*probLifespanCondR[k,])
    varLifespanCondR = sum((1:mA)^2*probLifespanCondR[k,]) -
      meanLifespanCondR^2
    sdLifespanCondR[k] = sqrt(varLifespanCondR)
    CVLifespanCondR[k] = sdLifespanCondR[k] / meanLifespanCondR
  }

  return (list(probLifespanCondR=probLifespanCondR,
               distKidsAtDeath=distKidsAtDeath,
               distAgeAtDeath=distAgeAtDeath,
               sdLifespanCondR=sdLifespanCondR,
               CVLifespanCondR=CVLifespanCondR))
}


#########################################################################
## Post-reproduction census partition for var. and skewness with no
## env. var.
##
#########################################################################

getVarSkewnessPartitionsNoEnvVarPostBreeding = function (P, F, c0, maxAge=100,
            esR1=NULL, esR2=NULL,
            esR3=NULL,
            bsR1=NULL, bsR2=NULL,
            bsR3=NULL, debugging=FALSE) {
  ## Input parameters:
  ## P is the survival/growth transition matrix (called U in COMPADRE)
  ## F is the matrix of fertility transitions
  ## c0 is the birth state distribution (also called mixing distribution)
  
  mz = dim(P)[1]
  
  ## Make survival, growth, and fecundity bullet matrices
  Sbullet = diag (colSums (P))
  
  Gbullet = P;  
  for(ic in 1:ncol(Gbullet)){
    if(Sbullet[ic,ic]>0) Gbullet[,ic]=Gbullet[,ic]/Sbullet[ic,ic]
  }       
  cat(range(Gbullet%*%Sbullet-P), "should equal 0", "\n"); 
  
  ## #kids isn't part of the state definition, so Fbullet = I
  Fbullet = diag (mz)
  
  ## create matrices for extended state space, which I will denote with
  ## the prefix "es".
  
  esmz = 3*mz
  esF = esP = matrix (0, esmz, esmz)
  
  esP[mz + 1:mz, 1:mz] = Sbullet
  esP[2*mz + 1:mz, mz + 1:mz] = Gbullet
  esP[1:mz, 2*mz + 1:mz] = Fbullet
    
  esF[1:mz, 2*mz + 1:mz] = F

  ################################################################
  ## Extended state model that includes ur-stage alpha_z.  Used for
  ## calculating contributions from birth stage.  The "bs" before
  ## variable names stands for "birth state."
  ################################################################

  bsmz = 1 + mz
  ## initial state --- everybody starts off in the ur-state.
  bsC0 = rep(0, bsmz)
  bsC0[1] = 1

  bsF = bsP = matrix (0, bsmz, bsmz)
  ## From ur-state to birth state
  bsP[1 + (1:mz),1] = c0
  ## From normal state to normal state
  bsP[1 + 1:mz, 1 + 1:mz] = P

  ## You only start reproducing once you're past the ur-state.  Note
  ## that all we need are the column sums of bsF.
  bsF[1, 1 + 1:mz] = colSums (F)

  ## set up reward matrices ###############################

  ## If the reward matrices aren't specified, we assume
  ## Poisson-distributed clutch sizes
  if (is.null(esR1)) {
    esR1 = esR2 = esR3 = matrix (0, esmz+1, esmz+1)
    bsR1 = bsR2 = bsR3 = matrix (0, bsmz+1, bsmz+1)

    ## First moment of clutch size (Poisson)
    for (j in 1:esmz) 
      esR1[,j] = sum(esF[,j])
    for (j in 1:bsmz) 
      bsR1[,j] = sum(bsF[,j])

    ## Second moment of clutch size  
    esR2 = esR1 + esR1^2
    bsR2 = bsR1 + bsR1^2
    ## Third moment of clutch size
    esR3 = esR1+ 3*esR1*esR1 + esR1*esR1*esR1
    bsR3 = bsR1+ 3*bsR1*bsR1 + bsR1*bsR1*bsR1
  }
  
  ## Get moments conditional on init. state #########################

  out = calcMoments (esP, esR1, esR2, esR3)
  esRho1Vec = out$rho1Vec
  esMu2Vec = out$mu2Vec
  esSkewnessVec = out$skewnessVec

  out = calcMoments (bsP, bsR1, bsR2, bsR3)
  bsRho1Vec = out$rho1Vec
  bsMu2Vec = out$mu2Vec
  bsMu3Vec = out$mu3Vec
  bsSkewnessVec = out$skewnessVec

  ## Start calculating luck terms ###################################

  survTrajecSkewness = growthTrajecSkewness = 
    fecSkewness = numeric (maxAge)
  fecUpdateSkewness = survUpdateSkewness =
    growthUpdateSkewness =  numeric (maxAge)
  survUpdateVar = growthUpdateVar = fecUpdateVar =
    numeric (maxAge) 
  survTrajecVar = growthTrajecVar = fecVar =
    numeric (maxAge)
  surv = numeric (maxAge)

  ## for ur-state var.
  ## expectation wrt z of Var(success | z)
  expZVarCondZ = bsMu2Vec %*% bsC0
  ## variance wrt z of Exp(success | z)
  varZExpCondZ = sum(bsC0*bsRho1Vec^2) - (sum(bsC0*bsRho1Vec)^2)
  ## Law of total variance
  urVar = expZVarCondZ + varZExpCondZ
  
  justBirthStateVar = bsMu2Vec %*% bsP %*% bsC0
  birthStateVar = urVar - justBirthStateVar

  ## for ur-state skewness
  expZMu3CondZ = bsMu3Vec %*% bsC0
  ## 3rd moment wrt z of Exp(success | z)
  rho3ZExpCondZ = sum(bsC0*(bsRho1Vec)^3)
  ## 2nd moment wrt z of Exp(success | z)
  rho2ZExpCondZ = sum(bsC0*(bsRho1Vec)^2)
  ## 1st moment wrt z of Exp(success | z)
  expZExpCondZ = sum(bsC0*bsRho1Vec)
  ## 3rd central moment wrt z of Exp(success | z)
  mu3ZExpCondZ = rho3ZExpCondZ -
    3*expZExpCondZ*rho2ZExpCondZ +
    2*expZExpCondZ^3
  ## covariance wrt z of Exp(success | z) and Var(success | z)
  covZExpVarCondZ = sum(bsC0*bsRho1Vec*bsMu2Vec) -
    sum(bsC0*bsRho1Vec)*sum(bsC0*bsMu2Vec)
  ## Law of total cumulance
  urMu3 = expZMu3CondZ + mu3ZExpCondZ +
    3*covZExpVarCondZ
  urSkewness = urMu3 / urVar^1.5
  
  justBirthStateSkewness = bsSkewnessVec %*% bsP %*% bsC0
  birthStateSkewness = urSkewness - justBirthStateSkewness
  
  ## Starting the age loop
  PaC0 = c0  ## P^a c_0
  mzZero = rep(0, mz)

  ## Sanity check: passes
  if (debugging) {
    N = solve(diag(mz) - P)
    rho1Vec = colSums (F %*% N)
    cat (esRho1Vec %*% c(c0, mzZero, mzZero),
         "should = ", rho1Vec %*% c0, "\n")
  }

  survUpdateSkewness[1] = esSkewnessVec %*% c(mzZero, Sbullet %*% PaC0, mzZero)
  growthUpdateSkewness[1] = esSkewnessVec %*% 
    c(mzZero, mzZero, P %*% PaC0)
  fecUpdateSkewness[1] = esSkewnessVec %*% 
    c(P %*% PaC0, mzZero, mzZero)

  survUpdateVar[1] = esMu2Vec %*% c(mzZero, Sbullet %*% PaC0, mzZero)
  growthUpdateVar[1] = esMu2Vec %*% c(mzZero, mzZero, P %*% PaC0)
  fecUpdateVar[1] = esMu2Vec %*% c(P %*% PaC0, mzZero, mzZero)

  surv[1] = sum (PaC0)
  ## a is actually age + 1, since the first year of life is age 0
  for (a in 2:maxAge) {
    PaC0 = P %*% PaC0

    survUpdateSkewness[a] = esSkewnessVec %*% c(mzZero, Sbullet %*% PaC0, mzZero)
    growthUpdateSkewness[a] = esSkewnessVec %*% 
      c(mzZero, mzZero, P %*% PaC0)
    fecUpdateSkewness[a] = esSkewnessVec %*% 
      c(P %*% PaC0, mzZero, mzZero)

    survUpdateVar[a] = esMu2Vec %*% c(mzZero, Sbullet %*% PaC0, mzZero)
    growthUpdateVar[a] = esMu2Vec %*% c(mzZero, mzZero, P %*% PaC0)
    fecUpdateVar[a] = esMu2Vec %*% c(P %*% PaC0, mzZero, mzZero)

    ## fecundity skewness and var.
    fecSkewness[a-1] = growthUpdateSkewness[a-1] - fecUpdateSkewness[a-1]
    fecVar[a-1] = growthUpdateVar[a-1] - fecUpdateVar[a-1]
    
    ## survival trajectory skewness and var.
    survTrajecSkewness[a] = fecUpdateSkewness[a-1] -
      survUpdateSkewness[a]
    survTrajecVar[a] = fecUpdateVar[a-1] - survUpdateVar[a]
    
    ## growth trajectory skewness and var.
    growthTrajecSkewness[a-1] = survUpdateSkewness[a-1] -
      growthUpdateSkewness[a-1] 
    growthTrajecVar[a-1] = survUpdateVar[a-1] - growthUpdateVar[a-1]

    surv[a] = sum(PaC0)
  }

  ## By what age are 99% of individuals dead?
  lifespan = which(surv < 0.01)[1]

  ## And get the first value of survSkewness and survVar
  survTrajecSkewness[1] = justBirthStateSkewness - survUpdateSkewness[1]
  survTrajecVar[1] = justBirthStateVar - survUpdateVar[1]

  ## By what age are 99% of individuals dead?
  lifespan = which(surv < 0.01)[1]

  totSurvTrajecSkewness = sum(survTrajecSkewness)
  totGrowthTrajecSkewness = sum(growthTrajecSkewness)
  totFecSkewness = sum(fecSkewness)
  totSkewness = bsSkewnessVec %*% bsC0
  ## Sanity check
  if (debugging) 
    cat (totSurvTrajecSkewness + totGrowthTrajecSkewness +
         totFecSkewness + birthStateSkewness, "should = ",
         totSkewness, "\n")

  totSurvTrajecVar = sum(survTrajecVar)
  totGrowthTrajecVar = sum(growthTrajecVar)
  totFecVar = sum(fecVar)
  totVar = bsMu2Vec %*% bsC0
  ## Sanity check
  if (debugging)
    cat (totSurvTrajecVar + totGrowthTrajecVar +
         totFecVar + birthStateVar, "should = ",
         totVar, "\n")

  return (out=list(birthStateVar=birthStateVar,
                   survTrajecVar=survTrajecVar,
                   growthTrajecVar=growthTrajecVar,
                   fecVar=fecVar,
                   totVar=totVar,
                   birthStateSkewness=birthStateSkewness,
                   survTrajecSkewness=survTrajecSkewness,
                   growthTrajecSkewness=growthTrajecSkewness,
                   fecSkewness=fecSkewness,
                   totSkewness=totSkewness,
                   lifespan=lifespan))
}

##
## If you don't supply a B matrix, this will assume
## Poisson-distributed clutch sizes

getVarSuccessEnvVar = function (numKidsThreshold, Plist, Flist, Q,
                                m0, maxAge=100, maxKids=30,
                                B=NULL, Fdist="Poisson")
{
  require (Matrix)
  debugging = TRUE
  
  mT = maxKids + 1
  kidsIndexThreshold = numKidsThreshold + 1
  
  mz = dim(Plist[[1]])[1]
  numEnv = dim(Q)[1]
  bigmz = numEnv*mz

  ## Define megamatrices M and F
  F = M = matrix (0, bigmz, bigmz)
  for (i in 1:numEnv) {
    for (j in 1:numEnv) {
      M[(i-1)*mz + 1:mz, (j-1)*mz + 1:mz] = Plist[[j]]*Q[i,j]
      F[(i-1)*mz + 1:mz, (j-1)*mz + 1:mz] = Flist[[j]]*Q[i,j]
    }
  }

  ## expected number of offspring in a clutch
  ## b = colSums (F)

  if (is.null(B)) 
    ## B[i,j] is the probability that a class-j individual has i-1 kids.
    ## We assume Poisson-distributed number of offspring.
    ## The columns of B should sum to 1 and they do.
    B = mk_B (maxKids, F)
  
  ## Construct A, the transition matrix for a (number of kids) x stage x
  ## env. model.   
  out = make_AxT (B, M, mT)
  A = out$A
  mzA = bigmz*mT
  K = out$K

  ## What *is* the distribution of kids at death? #########################

  ## In A, survival and growth come before reproduction, and we have
  ## assumed reproduction comes first.  Using A to calculate #kids
  ## at death will give the wrong answer.  Use esA instead.
  
  #####################################################################
  ## making the bullet matrices
  #####################################################################

  ## Make "bullet" matrices for A.
  Fbullet = Qbullet = Gbullet = matrix (0, mzA, mzA)

  ## Sbullet updates survival
  Sbullet = diag (colSums(A))

  ## Fbullet updates number of kids.  Remember to condition on survival.
  for (j in 1:mT) {  ## loop over current #kids
    for (i in j:(mT-1)) {  ## loop over future #kids
      for (z in 1:bigmz) {  ## loop over stage/env.
        if (Fdist == "Poisson") {
          Fbullet[(i-1)*bigmz + z, (j-1)*bigmz + z] =
            dpois (i-j, lambda=sum(F[,z]))
        } else if (Fdist == "Bernoulli") {
          Fbullet[(i-1)*bigmz + z, (j-1)*bigmz + z] =
            dbinom (i-j, prob=sum(F[,z]), size=1)
        } else {
          cat ("Did not recognize Fdist choice.\n")
        }
      }
    }
  }
  ## Make mT absorbing.  If you have #kids index < mT, the probability
  ## of going to mT while in stage z = 1 - all other kid transition
  ## probabilities with stage z.

  for (j in 1:(mT-1)) {
    for (z in 1:bigmz) {
      Fbullet[(mT-1)*bigmz + z, (j-1)*bigmz + z] =
        ##      1 - sum(Fbullet[1:((mT-1)*bigmz), (j-1)*bigmz + z])
        1 - sum(Fbullet[(0:(mT-2))*bigmz + z, (j-1)*bigmz + z])
    }
  }
  ## If you have #kids index = mT, you stay there.
  for (z in 1:bigmz) 
    Fbullet[(mT-1)*bigmz + z, (mT-1)*bigmz + z] = 1

  ## Qbullet updates env.
  smallQbullet = matrix (0, bigmz, bigmz)
  for (i in 1:numEnv) {
    for (j in 1:numEnv) {
      smallQbullet[(i-1)*mz + 1:mz, (j-1)*mz + 1:mz] = diag(mz)*Q[i,j]
    }
  }
  ## Qbullet is the same for each number of kids
  for (k in 1:mT) 
    Qbullet[(k-1)*bigmz + 1:bigmz, (k-1)*bigmz + 1:bigmz] = smallQbullet

  ## Gbullet updates stage
  smallGbullet = matrix (0, bigmz, bigmz)
  for (q in 1:numEnv) {
    S = diag (colSums (Plist[[q]]))
    Sinv = diag (1 / diag(S))
    G = Plist[[q]] %*% Sinv
    smallGbullet[(q-1)*mz + 1:mz, (q-1)*mz + 1:mz] = G
  }
  for (k in 1:mT) 
    Gbullet[(k-1)*bigmz + 1:bigmz, (k-1)*bigmz + 1:bigmz] = smallGbullet

  ## sanity check
  if (debugging)
    cat ("Checking the bullet matrices for all #kids:",
         range((Qbullet %*% Gbullet %*% Sbullet %*% Fbullet - A)),
         "should = 0.\n")

  ## This line is time-consuming for large matrices
  Pbullet = Gbullet %*% Sbullet %*% Fbullet

  ####################################################################
  ## Now make the additionally extended space matrix that allows us to
  ## stop partway through a time step.  Our new prefix is es to
  ## indicate this additional extension.  ("es" = "extended space")
  ####################################################################

  esmzA = 4*mzA

  ##esbigF = esA = matrix (0, esmzA, esmzA)
  esA = matrix (0, esmzA, esmzA)
  esA[mzA + 1:mzA, 1:mzA] = Fbullet
  esA[2*mzA + 1:mzA, mzA + 1:mzA] = Sbullet
  esA[3*mzA + 1:mzA, 2*mzA + 1:mzA] = Gbullet
  esA[1:mzA, 3*mzA + 1:mzA] = Qbullet

  ##################################################################
  ## What is the distribution of #kids at death when reproduction
  ## comes before survival and growth?
  ##################################################################

  ## Create a survival probability vector 
  surv = apply (esA, 2, sum);  die = 1-surv; 

  ## And the fundamental matrix of esA
  fundesA <- solve(diag(ncol(esA))-esA)

  ## Make omega
  omega = matrix(0,nrow(fundesA),ncol(fundesA)); 

  for(j in 1:ncol(fundesA))
    omega[,j] = die*fundesA[,j]

  distAtBirth = matrix(0, bigmz, 4*mT)
  distAtBirth[,1] = m0  ## Everyone starts off with zero kids at the
  ## beginning of the time step

  ## Get distribution of states at death
  distAtDeath = omega %*% matrix(distAtBirth, ncol=1)  # state vector
  ## distAtDeath = matrix(distAtDeath, nrow=bigmz) # state matrix
  ## distKidsAtDeath = apply(distAtDeath, 2, sum)[1:mT]
  distAtDeath = array (distAtDeath, dim=c(bigmz, mT, 4)) # state
                                       # matrix
  distKidsAtDeath = apply (distAtDeath, 2, sum)

  ## What threshold number of kids represents the 99th %ile of the LRO
  ## distribution? 
  cumDistKidsAtDeath = cumsum (distKidsAtDeath)
  R90 = which (cumDistKidsAtDeath > 0.9)[1]
  R99 = which (cumDistKidsAtDeath > 0.99)[1]
  if (debugging) {
    cat ("The 90th %ile of the LRO distribution occurs at", R90,
         "kids.\n")
    cat ("The 99th %ile of the LRO distribution occurs at", R99,
         "kids.\n")
  }


  #####################################################################
  ## Get probability of success conditional on starting state and from
  ## that, the variance of success.
  #####################################################################
  
  ## Define transient states: kids index < kidsIndexThreshold
  cutoff = (kidsIndexThreshold-1)*bigmz
  transientStates = c(1:cutoff, mzA + 1:cutoff, 2*mzA + 1:cutoff,
                      3*mzA + 1:cutoff)
  out = makeCondKernel (esA, transientStates)
  esQ2Extended = out$q2Extended
  varSuccessCondZ = esQ2Extended * (1 - esQ2Extended)

  ## Sanity check
  if (debugging & (Fdist == "Poisson") & (numKidsThreshold == 1)) {
    fbar = colSums(F)
    ## chance of one or more offspring for Poisson-dist. clutch size
    pb = 1 - exp(-fbar)
    P0 = matrix (NA, bigmz, bigmz)
    for (j in 1:bigmz)
      P0[,j] = M[,j]*(1 - pb[j]); ## Table 3.3 in The Book
    N0 = solve (diag(bigmz) - P0); B = pb%*%N0; ## Table 3.3
    AaM0 = c(m0, rep(0, (mT-1)*bigmz))
    mzAZero = rep(0, mzA)
    ## Does this work?  Remember that survival and growth come after
    ## fecundity for esA and its derived quantity, esQ2Extended.  Do
    ## they do so for M as well?  It seems to work for the
    ## age-structured callospermus code, though not for the
    ## stage-structured version.
    cat ("Prob. of having at least one kid = ",
         esQ2Extended %*% c(AaM0, mzAZero, mzAZero, mzAZero),
         "should = ", B %*% m0,
         "should = ", 1 - distKidsAtDeath[1], "\n") 
  }

  
  ##################################################################
  ## Start calculating trajectory lucks
  ##################################################################

  surv = survTrajecVar = growthTrajecVar = envTrajecVar =
    fecVar = numeric(maxAge)
  fecUpdateVar = survUpdateVar = growthUpdateVar =
    envUpdateVar = numeric (maxAge)

  ## The initial state m0 with zero kids
  AaM0 = c(m0, rep(0, (mT-1)*bigmz))

  mzAZero = rep(0, mzA)

  fecUpdateVar[1] = varSuccessCondZ %*%
    c(mzAZero, Fbullet %*% AaM0, mzAZero, mzAZero)

  survUpdateVar[1] = varSuccessCondZ %*%
    c(mzAZero, mzAZero, Sbullet %*% Fbullet %*% AaM0, mzAZero)

  growthUpdateVar[1] = varSuccessCondZ %*%
    c(mzAZero, mzAZero, mzAZero,  Pbullet %*% AaM0)

  envUpdateVar[1] = varSuccessCondZ %*%
    c(A %*% AaM0, mzAZero, mzAZero, mzAZero)

  surv[1] = sum (AaM0)

  for (a in 2:maxAge) {
    AaM0 = A %*% AaM0

    fecUpdateVar[a] = varSuccessCondZ %*%
      c(mzAZero, Fbullet %*% AaM0, mzAZero, mzAZero)

    survUpdateVar[a] = varSuccessCondZ %*%
      c(mzAZero, mzAZero, Sbullet %*% Fbullet %*% AaM0, mzAZero)

    growthUpdateVar[a] = varSuccessCondZ %*%
      c(mzAZero, mzAZero, mzAZero,  Pbullet %*% AaM0)

    envUpdateVar[a] = varSuccessCondZ %*%
      c(A %*% AaM0, mzAZero, mzAZero, mzAZero)

    ## Note: need to calculate fecVar[1] separately.
    ## fecundity because env. update happens at the end of the
    ## previous time step
    fecVar[a] = envUpdateVar[a-1] - fecUpdateVar[a]

    ## survival trajectory 
    survTrajecVar[a-1] = fecUpdateVar[a-1] -
      survUpdateVar[a-1]

    ## growth trajectory 
    growthTrajecVar[a-1] = survUpdateVar[a-1] -
      growthUpdateVar[a-1]

    ## environment trajectory
    envTrajecVar[a-1] = growthUpdateVar[a-1] -
      envUpdateVar[a-1]

    surv[a] = sum (AaM0)
  }

  ## What about birth env. luck?
  ## Extended state model that includes ur-stage alpha_z and
  ## ur-environment alpha_q.  Used for calculating contributions from
  ## birth stage and birth environment.  The "bs" before variable names
  ## stands for "birth state."

  ## Note: I have compared the action of bsA on the ur-state to esA on
  ## its initial state, up through the end of the first full time step,
  ## and they match.  Also, bsA %*% bsA * bsM0 should be the same as the
  ## initial condition for esA, and it is:
  ##
  ##    (bsA %*% bsA %*% bsM0)[1 + mz + 1:mzA] = AaM0,
  ##
  ## where AaM0 = c(m0, rep(0, (mT-1)*bigmz)).  I think that bsA works.

  bsmzA = 1 + mz + esmzA
  ## initial state
  bsM0 = rep(0, bsmzA)
  bsM0[1] = 1

  bsF = bsA = matrix (0, bsmzA, bsmzA)
  ## From ur-stage to birth stage and ur-environment
  bsA[1 + (1:mz),1] = c0
  ## From birth stage and ur-environment to m0
  for (q in 1:numEnv) {
    bsA[1 + q*mz + 1:mz, 1 + 1:mz] = diag (u0[q], mz)
  }
  ## From normal stage x env. state to normal stage x env. state
  bsA[1 + mz + 1:esmzA, 1 + mz + 1:esmzA] = esA

  ## Define transient states: kids index < kidsIndexThreshold
  cutoff = (kidsIndexThreshold-1)*bigmz
  transientStates = c(1:(1 + mz + cutoff),
                      mzA + 1:(1 + mz + cutoff),
                      2*mzA + 1:(1 + mz + cutoff),
                      3*mzA + 1:(1 + mz + cutoff))
  out = makeCondKernel (bsA, transientStates)
  bsQ2Extended = out$q2Extended
  bsVarSuccessCondZ = bsQ2Extended * (1 - bsQ2Extended)

  bsM0 = rep (0, bsmzA)
  bsM0[1] = 1
  expZVarCondZ = bsVarSuccessCondZ %*% bsM0
  varZExpCondZ = sum(bsM0*bsQ2Extended)^2 - sum(bsM0*bsQ2Extended)^2
  urVar = expZVarCondZ + varZExpCondZ

  justBirthStageVar = bsVarSuccessCondZ %*% bsA %*% bsM0
  birthStageEnvVar = bsVarSuccessCondZ %*% bsA %*% (bsA %*% bsM0)

  birthStateVar = urVar - justBirthStageVar
  birthEnvVar = justBirthStageVar - birthStageEnvVar
  fecVar[1] = birthStageEnvVar - fecUpdateVar[1]

  totEnvTrajecVar = sum(envTrajecVar)
  totSurvTrajecVar = sum(survTrajecVar)
  totGrowthTrajecVar = sum(growthTrajecVar)
  totFecVar = sum(fecVar)

  totVar = totEnvTrajecVar + totSurvTrajecVar +
    totGrowthTrajecVar + totFecVar + birthStateVar + birthEnvVar

  ## By what age are 99% of individuals dead?
  lifespan = which(surv < 0.01)[1]

  ## mean lifespan
  N = solve(diag(bigmz) - M)
  meanLifespan = colSums(N) %*% m0

  ## Sanity check
  if (debugging) {
    AaM0 = c(m0, rep(0, (mT-1)*bigmz))
    cat (varSuccessCondZ %*% c(AaM0, mzAZero, mzAZero, mzAZero), "should = ",
  ##       expZVarCondZ, "should = ",
         totEnvTrajecVar + totSurvTrajecVar +
         totGrowthTrajecVar + totFecVar, "\n")
  }
  
  if (debugging) {
    dev.new ()
    par (cex.lab=1.4, cex.axis=1.3, lwd=2,bty="l",mgp=c(2,1,0), mar=c(4,4,1,1))
    ymax = max(survTrajecVar, growthTrajecVar, envTrajecVar, fecVar)
    ymin = min(survTrajecVar, growthTrajecVar, envTrajecVar, fecVar)
    plot (0:25, survTrajecVar[1:26], type="l", xlab="Age",
          ylab="Contrib. to Var(success)", ylim=c(ymin, ymax))
    lines (0:25, growthTrajecVar[1:26], type="l",
           col="orange", cex=1.2)
    lines (0:25, envTrajecVar[1:26], type="l",
           col="green", cex=1.2)
    lines (0:25, fecVar[1:26], type="l", col="blue")
    lines (rep(lifespan, 2), c(0, 0.4*ymax), col="black",
           lty=2, lwd=2)
    lines (0:25, rep(0, 26), lty=2, lwd=1)

    legend (x="topright",
            legend=c("Surv. trajectory", "Growth trajectory", "Env trajectory",
                     "Fecundity", "90% of individ. are dead"),
            inset=0.05, cex=1.2,
            col=c("black", "orange", "green", "blue", "black"),
            lty=c(1,1,1,1,2), lwd=2, bty="n")
  }

  return (out=list(survTrajecVar=survTrajecVar,
                   growthTrajecVar=growthTrajecVar,
                   envTrajecVar=envTrajecVar,
                   fecVar=fecVar,
                   totSurvTrajecVar=totSurvTrajecVar,
                   totGrowthTrajecVar=totGrowthTrajecVar,
                   totEnvTrajecVar=totEnvTrajecVar,
                   totFecVar=totFecVar,
##                   birthStateVar=birthStateVar,
##                   birthEnvVar=birthEnvVar,
                   totVar=totVar,
                   lifespan=lifespan,
                   meanLifespan=meanLifespan))
}

## This version loops over #kids threshold values until the 99th %ile
## of LRO.
##
## If you don't supply a B matrix, this will assume
## Poisson-distributed clutch sizes
##
## percentileCutoff = the percentile of the LRO distribution to which
## you want the #kids success threshold to loop.  E.g.  If you want
## the success threshold to eventually be as high as 99%, set
## percentileCutoff = 0.99.  Note that for some life histories, over
## 90% of individuals have zero kids.
loopVarSuccessEnvVar = function (Plist, Flist, Q,
                                 m0, maxAge=100, maxKids=30,
                                 percentileCutoff=0.99,
                                 B=NULL, Fdist="Poisson")
{
  require (Matrix)
  debugging = TRUE
  
  mT = maxKids + 1
  
  mz = dim(Plist[[1]])[1]
  numEnv = dim(Q)[1]
  bigmz = numEnv*mz

  ## Define megamatrices M and F
  F = M = matrix (0, bigmz, bigmz)
  for (i in 1:numEnv) {
    for (j in 1:numEnv) {
      M[(i-1)*mz + 1:mz, (j-1)*mz + 1:mz] = Plist[[j]]*Q[i,j]
      F[(i-1)*mz + 1:mz, (j-1)*mz + 1:mz] = Flist[[j]]*Q[i,j]
    }
  }

  ## expected number of offspring in a clutch
  ## b = colSums (F)

  if (is.null(B)) 
    ## B[i,j] is the probability that a class-j individual has i-1 kids.
    ## We assume Poisson-distributed number of offspring.
    ## The columns of B should sum to 1 and they do.
    B = mk_B (maxKids, F)
  
  ## Construct A, the transition matrix for a (number of kids) x stage x
  ## env. model.   
  out = make_AxT (B, M, mT)
  A = out$A
  mzA = bigmz*mT
  K = out$K

  #####################################################################
  ## making the bullet matrices
  #####################################################################

  ## Make "bullet" matrices for A.
  Fbullet = Qbullet = Gbullet = matrix (0, mzA, mzA)

  ## Sbullet updates survival
  Sbullet = diag (colSums(A))

  ## Fbullet updates number of kids.
  for (j in 1:mT) {
    for (i in j:(mT-1)) {
      for (z in 1:bigmz) {
        if (Fdist == "Poisson") {
          Fbullet[(i-1)*bigmz + z, (j-1)*bigmz + z] =
            dpois (i-j, lambda=sum(F[,z]))
        } else if (Fdist == "Bernoulli") {
          Fbullet[(i-1)*bigmz + z, (j-1)*bigmz + z] =
            dbinom (i-j, prob=sum(F[,z]), size=1)
        } else {
          cat ("Did not recognize Fdist choice.\n")
        }
      }
    }
  }
  ## Make mT absorbing.  If you have #kids index < mT, the probability
  ## of going to mT while in stage z = 1 - all other kid transition
  ## probabilities with stage z.

  for (j in 1:(mT-1)) {
    for (z in 1:bigmz) {
      Fbullet[(mT-1)*bigmz + z, (j-1)*bigmz + z] =
        ##      1 - sum(Fbullet[1:((mT-1)*bigmz), (j-1)*bigmz + z])
        1 - sum(Fbullet[(0:(mT-2))*bigmz + z, (j-1)*bigmz + z])
    }
  }
  ## If you have #kids index = mT, you stay there.
  for (z in 1:bigmz) 
    Fbullet[(mT-1)*bigmz + z, (mT-1)*bigmz + z] = 1

  ## Qbullet updates env.
  smallQbullet = matrix (0, bigmz, bigmz)
  for (i in 1:numEnv) {
    for (j in 1:numEnv) {
      smallQbullet[(i-1)*mz + 1:mz, (j-1)*mz + 1:mz] = diag(mz)*Q[i,j]
    }
  }
  ## Qbullet is the same for each number of kids
  for (k in 1:mT) 
    Qbullet[(k-1)*bigmz + 1:bigmz, (k-1)*bigmz + 1:bigmz] = smallQbullet

  ## Gbullet updates stage
  smallGbullet = matrix (0, bigmz, bigmz)
  for (q in 1:numEnv) {
    S = diag (colSums (Plist[[q]]))
    Sinv = diag (1 / diag(S))
    G = Plist[[q]] %*% Sinv
    smallGbullet[(q-1)*mz + 1:mz, (q-1)*mz + 1:mz] = G
  }
  for (k in 1:mT) 
    Gbullet[(k-1)*bigmz + 1:bigmz, (k-1)*bigmz + 1:bigmz] = smallGbullet

  ## sanity check
  if (debugging)
    cat ("Checking the bullet matrices for all #kids:",
         range((Qbullet %*% Gbullet %*% Sbullet %*% Fbullet - A)),
         "should = 0.\n")

  ## This line is time-consuming for large matrices
  Pbullet = Gbullet %*% Sbullet %*% Fbullet

  ####################################################################
  ## Now make the additionally extended space matrix that allows us to
  ## stop partway through a time step.  Our new prefix is es to
  ## indicate this additional extension.  ("es" = "extended space")
  ####################################################################

  esmzA = 4*mzA

  ##esbigF = esA = matrix (0, esmzA, esmzA)
  esA = matrix (0, esmzA, esmzA)
  esA[mzA + 1:mzA, 1:mzA] = Fbullet
  esA[2*mzA + 1:mzA, mzA + 1:mzA] = Sbullet
  esA[3*mzA + 1:mzA, 2*mzA + 1:mzA] = Gbullet
  esA[1:mzA, 3*mzA + 1:mzA] = Qbullet

  ##################################################################
  ## What is the distribution of #kids at death when reproduction
  ## comes before survival and growth?
  ##################################################################

  ## Create a survival probability vector 
  surv = apply (esA, 2, sum);  die = 1-surv; 

  ## And the fundamental matrix of esA
  fundesA <- solve(diag(ncol(esA))-esA)

  ## Make omega
  omega = matrix(0,nrow(fundesA),ncol(fundesA)); 

  for(j in 1:ncol(fundesA))
    omega[,j] = die*fundesA[,j]

  distAtBirth = matrix(0, bigmz, 4*mT)
  distAtBirth[,1] = m0  ## Everyone starts off with zero kids at the
  ## beginning of the time step

  ## Get distribution of states at death
  distAtDeath = omega %*% matrix(distAtBirth, ncol=1)  # state vector
  ## distAtDeath = matrix(distAtDeath, nrow=bigmz) # state matrix
  ## distKidsAtDeath = apply(distAtDeath, 2, sum)[1:mT]
  distAtDeath = array (distAtDeath, dim=c(bigmz, mT, 4)) # state
                                       # matrix
  distKidsAtDeath = apply (distAtDeath, 2, sum)

  ## What threshold number of kids represents the 99th %ile of the LRO
  ## distribution? 
  cumDistKidsAtDeath = cumsum (distKidsAtDeath)
  R90 = which (cumDistKidsAtDeath > 0.9)[1]
  R99 = which (cumDistKidsAtDeath > 0.99)[1]
  if (debugging) {
    cat ("The 90th %ile of the LRO distribution occurs at", R90,
         "kids.\n")
    cat ("The 99th %ile of the LRO distribution occurs at", R99,
         "kids.\n")
  }
  RCutoff = which (cumDistKidsAtDeath > percentileCutoff)[1]

  surv = survTrajecVar = growthTrajecVar = envTrajecVar =
    fecVar = numeric(maxAge)
  fecUpdateVar = survUpdateVar = growthUpdateVar =
    envUpdateVar = numeric (maxAge)

  totSurvTrajecVar = proporSurvTrajecVar = numeric (RCutoff)
  totGrowthTrajecVar = proporGrowthTrajecVar = numeric (RCutoff)
  totEnvTrajecVar = proporEnvTrajecVar = numeric (RCutoff)
  totFecVar = proporFecVar = numeric (RCutoff)
  birthStateVar = birthEnvVar = numeric (RCutoff)

  for (numKidsThreshold in 1:RCutoff) {
    cat ("#kids threshold = ", numKidsThreshold, "\n")
    kidsIndexThreshold = numKidsThreshold + 1
    
    #####################################################################
    ## Get probability of success conditional on starting state and from
    ## that, the variance of success.
    #####################################################################
    
    ## Define transient states: kids index < kidsIndexThreshold
    cutoff = (kidsIndexThreshold-1)*bigmz
    transientStates = c(1:cutoff, mzA + 1:cutoff, 2*mzA + 1:cutoff,
                        3*mzA + 1:cutoff)
    out = makeCondKernel (esA, transientStates)
    esQ2Extended = out$q2Extended
    varSuccessCondZ = esQ2Extended * (1 - esQ2Extended)

    ## Sanity check
    if (debugging & (Fdist == "Poisson") & (numKidsThreshold == 1)) {
      fbar = colSums(F)
      ## chance of one or more offspring for Poisson-dist. clutch size
      pb = 1 - exp(-fbar)
      P0 = matrix (NA, bigmz, bigmz)
      for (j in 1:bigmz)
        P0[,j] = M[,j]*(1 - pb[j]); ## Table 3.3 in The Book
      N0 = solve (diag(bigmz) - P0); B = pb%*%N0; ## Table 3.3
      AaM0 = c(m0, rep(0, (mT-1)*bigmz))
      mzAZero = rep(0, mzA)
      ## Does this work?  Remember that survival and growth come after
      ## fecundity for esA and its derived quantity, esQ2Extended.  Do
      ## they do so for M as well?  It seems to work for the
      ## age-structured callospermus code, though not for the
      ## stage-structured version.
      cat ("Prob. of having at least one kid = ",
           esQ2Extended %*% c(AaM0, mzAZero, mzAZero, mzAZero),
           "should = ", B %*% m0,
           "should = ", 1 - distKidsAtDeath[1], "\n") 
    }

    ##################################################################
    ## Start calculating trajectory lucks
    ##################################################################

    ## The initial state m0 with zero kids
    AaM0 = c(m0, rep(0, (mT-1)*bigmz))

    mzAZero = rep(0, mzA)

    fecUpdateVar[1] = varSuccessCondZ %*%
      c(mzAZero, Fbullet %*% AaM0, mzAZero, mzAZero)

    survUpdateVar[1] = varSuccessCondZ %*%
      c(mzAZero, mzAZero, Sbullet %*% Fbullet %*% AaM0, mzAZero)

    growthUpdateVar[1] = varSuccessCondZ %*%
      c(mzAZero, mzAZero, mzAZero,  Pbullet %*% AaM0)

    envUpdateVar[1] = varSuccessCondZ %*%
      c(A %*% AaM0, mzAZero, mzAZero, mzAZero)

    surv[1] = sum (AaM0)

    for (a in 2:maxAge) {
      cat ("a = ", a, "\n")
      AaM0 = A %*% AaM0

      fecUpdateVar[a] = varSuccessCondZ %*%
        c(mzAZero, Fbullet %*% AaM0, mzAZero, mzAZero)

      survUpdateVar[a] = varSuccessCondZ %*%
        c(mzAZero, mzAZero, Sbullet %*% Fbullet %*% AaM0, mzAZero)

      growthUpdateVar[a] = varSuccessCondZ %*%
        c(mzAZero, mzAZero, mzAZero,  Pbullet %*% AaM0)

      envUpdateVar[a] = varSuccessCondZ %*%
        c(A %*% AaM0, mzAZero, mzAZero, mzAZero)

      ## Note: need to calculate fecVar[1] separately.
      ## fecundity because env. update happens at the end of the
      ## previous time step
      fecVar[a] = envUpdateVar[a-1] - fecUpdateVar[a]

      ## survival trajectory 
      survTrajecVar[a-1] = fecUpdateVar[a-1] -
        survUpdateVar[a-1]

      ## growth trajectory 
      growthTrajecVar[a-1] = survUpdateVar[a-1] -
        growthUpdateVar[a-1]

      ## environment trajectory
      envTrajecVar[a-1] = growthUpdateVar[a-1] -
        envUpdateVar[a-1]

      surv[a] = sum (AaM0)
    }

    ## What about birth env. luck?
    ## Extended state model that includes ur-stage alpha_z and
    ## ur-environment alpha_q.  Used for calculating contributions from
    ## birth stage and birth environment.  The "bs" before variable names
    ## stands for "birth state."

    ## Note: I have compared the action of bsA on the ur-state to esA on
    ## its initial state, up through the end of the first full time step,
    ## and they match.  Also, bsA %*% bsA * bsM0 should be the same as the
    ## initial condition for esA, and it is:
    ##
    ##    (bsA %*% bsA %*% bsM0)[1 + mz + 1:mzA] = AaM0,
    ##
    ## where AaM0 = c(m0, rep(0, (mT-1)*bigmz)).  I think that bsA works.

    bsmzA = 1 + mz + esmzA
    ## initial state
    bsM0 = rep(0, bsmzA)
    bsM0[1] = 1

    bsF = bsA = matrix (0, bsmzA, bsmzA)
    ## From ur-stage to birth stage and ur-environment
    bsA[1 + (1:mz),1] = c0
    ## From birth stage and ur-environment to m0
    for (q in 1:numEnv) {
      bsA[1 + q*mz + 1:mz, 1 + 1:mz] = diag (u0[q], mz)
    }
    ## From normal stage x env. state to normal stage x env. state
    bsA[1 + mz + 1:esmzA, 1 + mz + 1:esmzA] = esA

    ## Define transient states: kids index < kidsIndexThreshold
    cutoff = (kidsIndexThreshold-1)*bigmz
    transientStates = c(1:(1 + mz + cutoff),
                        1 + mz + mzA + 1:cutoff,
                        1 + mz + 2*mzA + 1:cutoff,
                        1 + mz + 3*mzA + 1:cutoff)
    out = makeCondKernel (bsA, transientStates)
    bsQ2Extended = out$q2Extended
    bsVarSuccessCondZ = bsQ2Extended * (1 - bsQ2Extended)

    bsM0 = rep (0, bsmzA)
    bsM0[1] = 1
    expZVarCondZ = bsVarSuccessCondZ %*% bsM0
    varZExpCondZ = sum(bsM0*bsQ2Extended)^2 - sum(bsM0*bsQ2Extended)^2
    urVar = expZVarCondZ + varZExpCondZ

    justBirthStageVar = bsVarSuccessCondZ %*% bsA %*% bsM0
    birthStageEnvVar = bsVarSuccessCondZ %*% bsA %*% (bsA %*% bsM0)

    birthStateVar[numKidsThreshold] = urVar - justBirthStageVar
    birthEnvVar[numKidsThreshold] = justBirthStageVar - birthStageEnvVar
    fecVar[1] = birthStageEnvVar - fecUpdateVar[1]

    totEnvTrajecVar[numKidsThreshold] = sum(envTrajecVar)
    totSurvTrajecVar[numKidsThreshold] = sum(survTrajecVar)
    totGrowthTrajecVar[numKidsThreshold] = sum(growthTrajecVar)
    totFecVar[numKidsThreshold] = sum(fecVar)

    totVar = totEnvTrajecVar[numKidsThreshold] +
      totSurvTrajecVar[numKidsThreshold] +
      totGrowthTrajecVar[numKidsThreshold] +
      totFecVar[numKidsThreshold] +
      birthStateVar[numKidsThreshold] +
      birthEnvVar[numKidsThreshold]

    proporSurvTrajecVar[numKidsThreshold] =
      totSurvTrajecVar[numKidsThreshold] / totVar
    proporGrowthTrajecVar[numKidsThreshold] =
      totGrowthTrajecVar[numKidsThreshold] / totVar
    proporEnvTrajecVar[numKidsThreshold] =
      totEnvTrajecVar[numKidsThreshold] / totVar
    proporFecVar[numKidsThreshold] =
      totFecVar[numKidsThreshold] / totVar

    ## By what age are 99% of individuals dead?
    lifespan = which(surv < 0.01)[1]

    ## mean lifespan
    N = solve(diag(bigmz) - M)
    meanLifespan = colSums(N) %*% m0
    
  } ## end loop over numKidsThreshold
  
  return (out=list(proporSurvTrajecVar=proporSurvTrajecVar,
                   proporGrowthTrajecVar=proporGrowthTrajecVar,
                   proporEnvTrajecVar=proporEnvTrajecVar,
                   proporFecVar=proporFecVar,
                   totSurvTrajecVar=totSurvTrajecVar,
                   totGrowthTrajecVar=totGrowthTrajecVar,
                   totEnvTrajecVar=totEnvTrajecVar,
                   totFecVar=totFecVar,
##                   birthStateVar=birthStateVar,
##                   birthEnvVar=birthEnvVar,
                   totVar=totVar,
                   lifespan=lifespan,
                   meanLifespan=meanLifespan))
}

loopFanovaProbSuccess = function (Plist, Flist, Q,
                                  m0, maxKids=30,
                                  percentileCutoff=0.99,
                                  baseline="failure",
                                  treatment="success",
                                  B=NULL, Fdist="Poisson")
{
  require (Matrix)
  debugging = TRUE
  
  mT = maxKids + 1
  
  mz = dim(Plist[[1]])[1]
  numEnv = dim(Q)[1]
  bigmz = numEnv*mz

  ## Define megamatrices M and F
  F = M = matrix (0, bigmz, bigmz)
  for (i in 1:numEnv) {
    for (j in 1:numEnv) {
      M[(i-1)*mz + 1:mz, (j-1)*mz + 1:mz] = Plist[[j]]*Q[i,j]
      F[(i-1)*mz + 1:mz, (j-1)*mz + 1:mz] = Flist[[j]]*Q[i,j]
    }
  }

  ## expected number of offspring in a clutch
  ## b = colSums (F)

  if (is.null(B)) 
    ## B[i,j] is the probability that a class-j individual has i-1 kids.
    ## We assume Poisson-distributed number of offspring.
    ## The columns of B should sum to 1 and they do.
    B = mk_B (maxKids, F)

  ## Construct A, the transition matrix for a (number of kids) x stage x
  ## env. model.   
  out = make_AxT (B, M, mT)
  A = out$A
  mzA = bigmz*mT
  K = out$K

  #####################################################################
  ## making the bullet matrices
  #####################################################################

  ## Make "bullet" matrices for A.
  Fbullet = Qbullet = Gbullet = matrix (0, mzA, mzA)

  ## Sbullet updates survival
  Sbullet = diag (colSums(A))

  ## Fbullet updates number of kids.
  for (j in 1:mT) {
    for (i in j:(mT-1)) {
      for (z in 1:bigmz) {
        if (Fdist == "Poisson") {
          Fbullet[(i-1)*bigmz + z, (j-1)*bigmz + z] =
            dpois (i-j, lambda=sum(F[,z]))
        } else if (Fdist == "Bernoulli") {
          Fbullet[(i-1)*bigmz + z, (j-1)*bigmz + z] =
            dbinom (i-j, prob=sum(F[,z]), size=1)
        } else {
          cat ("Did not recognize Fdist choice.\n")
        }
      }
    }
  }
  ## Make mT absorbing.  If you have #kids index < mT, the probability
  ## of going to mT while in stage z = 1 - all other kid transition
  ## probabilities with stage z.

  for (j in 1:(mT-1)) {
    for (z in 1:bigmz) {
      Fbullet[(mT-1)*bigmz + z, (j-1)*bigmz + z] =
        ##      1 - sum(Fbullet[1:((mT-1)*bigmz), (j-1)*bigmz + z])
        1 - sum(Fbullet[(0:(mT-2))*bigmz + z, (j-1)*bigmz + z])
    }
  }
  ## If you have #kids index = mT, you stay there.
  for (z in 1:bigmz) 
    Fbullet[(mT-1)*bigmz + z, (mT-1)*bigmz + z] = 1

  ## Qbullet updates env.
  smallQbullet = matrix (0, bigmz, bigmz)
  for (i in 1:numEnv) {
    for (j in 1:numEnv) {
      smallQbullet[(i-1)*mz + 1:mz, (j-1)*mz + 1:mz] = diag(mz)*Q[i,j]
    }
  }
  ## Qbullet is the same for each number of kids
  for (k in 1:mT) 
    Qbullet[(k-1)*bigmz + 1:bigmz, (k-1)*bigmz + 1:bigmz] = smallQbullet

  ## Gbullet updates stage
  smallGbullet = matrix (0, bigmz, bigmz)
  for (q in 1:numEnv) {
    S = diag (colSums (Plist[[q]]))
    Sinv = diag (1 / diag(S))
    G = Plist[[q]] %*% Sinv
    smallGbullet[(q-1)*mz + 1:mz, (q-1)*mz + 1:mz] = G
  }
  for (k in 1:mT) 
    Gbullet[(k-1)*bigmz + 1:bigmz, (k-1)*bigmz + 1:bigmz] = smallGbullet

  ## sanity check
  if (debugging)
    cat ("Checking the bullet matrices for all #kids:",
         range((Qbullet %*% Gbullet %*% Sbullet %*% Fbullet - A)),
         "should = 0.\n")


  ####################################################################
  ## Now make the additionally extended space matrix that allows us to
  ## stop partway through a time step.  Our new prefix is es to
  ## indicate this additional extension.  ("es" = "extended space")
  ####################################################################
  
  esmzA = 4*mzA

  ##esbigF = esA = matrix (0, esmzA, esmzA)
  esA = matrix (0, esmzA, esmzA)
  esA[mzA + 1:mzA, 1:mzA] = Fbullet
  esA[2*mzA + 1:mzA, mzA + 1:mzA] = Sbullet
  esA[3*mzA + 1:mzA, 2*mzA + 1:mzA] = Gbullet
  esA[1:mzA, 3*mzA + 1:mzA] = Qbullet

  ##################################################################
  ## What is the distribution of #kids at death when reproduction
  ## comes before survival and growth?
  ##################################################################

  ## Create a survival probability vector 
  surv = apply (esA, 2, sum);  die = 1-surv; 

  ## And the fundamental matrix of esA
  fundesA <- solve(diag(ncol(esA))-esA)

  ## Make omega
  omega = matrix(0,nrow(fundesA),ncol(fundesA)); 

  for(j in 1:ncol(fundesA))
    omega[,j] = die*fundesA[,j]

  distAtBirth = matrix(0, bigmz, 4*mT)
  distAtBirth[,1] = m0  ## Everyone starts off with zero kids at the
  ## beginning of the time step

  ## Get distribution of states at death
  distAtDeath = omega %*% matrix(distAtBirth, ncol=1)  # state vector
  ## distAtDeath = matrix(distAtDeath, nrow=bigmz) # state matrix
  ## distKidsAtDeath = apply(distAtDeath, 2, sum)[1:mT]
  distAtDeath = array (distAtDeath, dim=c(bigmz, mT, 4)) # state
                                       # matrix
  distKidsAtDeath = apply (distAtDeath, 2, sum)

  ## What threshold number of kids represents the 99th %ile of the LRO
  ## distribution? 
  cumDistKidsAtDeath = cumsum (distKidsAtDeath)
  R90 = which (cumDistKidsAtDeath > 0.9)[1]
  R99 = which (cumDistKidsAtDeath > 0.99)[1]
  if (debugging) {
    cat ("The 90th %ile of the LRO distribution occurs at", R90,
         "kids.\n")
    cat ("The 99th %ile of the LRO distribution occurs at", R99,
         "kids.\n")
  }
  RCutoff = which (cumDistKidsAtDeath > percentileCutoff)[1]

  epsilonSurv = proporSurvMain = numeric (RCutoff)
  epsilonGrowth = proporGrowthMain = numeric (RCutoff)
  epsilonEnv = proporEnvMain = numeric (RCutoff)
  epsilonFec = proporFecMain = numeric (RCutoff)
  epsilonFecSurv = proporFecSurvMain = numeric (RCutoff)
  epsilonFecGrowth = proporFecGrowthMain = numeric (RCutoff)
  epsilonFecEnv = proporFecEnvMain = numeric (RCutoff)
  remainder = proporRemainder = numeric (RCutoff)
  epsilonZero = proporEpsilonZero = numeric (RCutoff)

  FProbSuccess = SProbSuccess = GProbSuccess =
    QProbSuccess = allTreatmentProbSuccess = numeric (RCutoff)
  FSProbSuccess = FGProbSuccess = FQProbSuccess =
    numeric (RCutoff)

  for (numKidsThreshold in 1:RCutoff) {
    cat ("#kids threshold = ", numKidsThreshold, "\n")
    kidsIndexThreshold = numKidsThreshold + 1

    ## Get esA conditional on failure and from that deduce the
    ## "bullet" matrices (called *circ instead of *bullet to
    ## distinguish them from the bullet matrices for the unconditional
    ## esA).
    cutoff = (kidsIndexThreshold-1)*bigmz
    transientStates = c(1:cutoff, mzA + 1:cutoff, 2*mzA + 1:cutoff,
                        3*mzA + 1:cutoff)
    out = makeCondFailureKernel (esA, transientStates)
    condEsA = out$MCond

    Fcirc = Scirc = Gcirc = Qcirc = matrix (0, mzA, mzA)
    Fcirc[1:cutoff, 1:cutoff] =
      condEsA[cutoff + 1:cutoff, 1:cutoff]
    ## Survival, growth, and env. transitions depend only on z and q,
    ## not #kids, so we can extend the circ matrices to non-transient
    ## states (i.e. those with #kids >= the threshold) by just
    ## repeating their (z,q)-based entries.  E.g., diag(Sbullet) and
    ## diag(Sbullet) just repeat after every bigmz entries.
    Scirc[1:cutoff, 1:cutoff] =
      condEsA[2*cutoff + 1:cutoff, cutoff + 1:cutoff]
    Gcirc[1:cutoff, 1:cutoff] =
      condEsA[3*cutoff + 1:cutoff, 2*cutoff + 1:cutoff]
    Qcirc[1:cutoff, 1:cutoff] =
      condEsA[1:cutoff, 3*cutoff + 1:cutoff]
    
    for (k in kidsIndexThreshold:mT) {
      Scirc[(k-1)*bigmz + 1:bigmz, (k-1)*bigmz + 1:bigmz] =
        condEsA[2*bigmz + 1:bigmz, bigmz + 1:bigmz]
      Gcirc[(k-1)*bigmz + 1:bigmz, (k-1)*bigmz + 1:bigmz] =
        condEsA[3*bigmz + 1:bigmz, 2*bigmz + 1:bigmz]
      Qcirc[(k-1)*bigmz + 1:bigmz, (k-1)*bigmz + 1:bigmz] =
        condEsA[1:bigmz, 3*bigmz + 1:bigmz]
    }

    ## Get esA conditional on success and from that deduce the
    ## "bullet" matrices (called *star instead of *bullet to
    ## distinguish them from the bullet matrices for the unconditional
    ## esA).
    cutoff = (kidsIndexThreshold-1)*bigmz
    transientStates = c(1:cutoff, mzA + 1:cutoff, 2*mzA + 1:cutoff,
                        3*mzA + 1:cutoff)
    out = makeCondKernel (esA, transientStates)
    condEsA = out$MCond
    ## Sanity check: passes
    if (debugging) {
      out = makeCondKernel (condEsA, transientStates)
      esQ2Extended = out$q2Extended
      AaM0 = c(m0, rep(0, (mT-1)*bigmz))
      mzAZero = rep(0, mzA)
      cat("Probability of success for the kernel conditional on success is",
          esQ2Extended %*% c(AaM0, rep(mzAZero, 3)), "\n")
    }

    Fstar = condEsA[mzA + 1:mzA, 1:mzA]
    Sstar = condEsA[2*mzA + 1:mzA, mzA + 1:mzA]
    Gstar = condEsA[3*mzA + 1:mzA, 2*mzA + 1:mzA]
    Qstar = condEsA[1:mzA, 3*mzA + 1:mzA]

    ## What is baseline and what is treatment?

    if (baseline == "failure") {
      ## baseline is guaranteed failure, treatment = guaranteed success
      baselineF = Fcirc
      baselineS = Scirc
      baselineG = Gcirc
      baselineQ = Qcirc
    }

    if (treatment == "success") {
      treatmentF = Fstar
      treatmentS = Sstar
      treatmentG = Gstar
      treatmentQ = Qstar
    } else if (treatment == "normal") {
      treatmentF = Fbullet
      treatmentS = Sbullet
      treatmentG = Gbullet
      treatmentQ = Qbullet
    }
      
    ## Transient states for all the following
    transientStates = c(1:cutoff, mzA + 1:cutoff, 2*mzA + 1:cutoff,
                        3*mzA + 1:cutoff)
    ## Initial state
    AaM0 = c(m0, rep(0, (mT-1)*bigmz))
    mzAZero = rep(0, mzA)

    ## Get epsilonZero (all baseline)

    esCondA = matrix (0, esmzA, esmzA)
    esCondA[mzA + 1:mzA, 1:mzA] = baselineF
    esCondA[2*mzA + 1:mzA, mzA + 1:mzA] = baselineS
    esCondA[3*mzA + 1:mzA, 2*mzA + 1:mzA] = baselineG
    esCondA[1:mzA, 3*mzA + 1:mzA] = baselineQ

    out = makeCondKernel (esCondA, transientStates)
    esQ2Extended = out$q2Extended

    AaM0 = c(m0, rep(0, (mT-1)*bigmz))
    mzAZero = rep(0, mzA)
    epsilonZero[numKidsThreshold] = esQ2Extended %*%
      c(AaM0, rep(mzAZero, 3))

    ## Sanity check: passes
    if (debugging) {
      cat("Probability of success for the kernel conditional on failure is",
          epsilonZero[numKidsThreshold], "\n")
    }

    ## Get main effect of F #################

    esCondA = matrix (0, esmzA, esmzA)
    esCondA[mzA + 1:mzA, 1:mzA] = treatmentF
    esCondA[2*mzA + 1:mzA, mzA + 1:mzA] = baselineS
    esCondA[3*mzA + 1:mzA, 2*mzA + 1:mzA] = baselineG
    esCondA[1:mzA, 3*mzA + 1:mzA] = baselineQ

    out = makeCondKernel (esCondA, transientStates)
    esQ2Extended = out$q2Extended

    FProbSuccess[numKidsThreshold] = esQ2Extended %*%
      c(AaM0, rep(mzAZero, 3))

    epsilonFec[numKidsThreshold] =
      (FProbSuccess - epsilonZero)[numKidsThreshold] 

    ## Get main effect of S #################

    esCondA = matrix (0, esmzA, esmzA)
    esCondA[mzA + 1:mzA, 1:mzA] = baselineF
    esCondA[2*mzA + 1:mzA, mzA + 1:mzA] = treatmentS
    esCondA[3*mzA + 1:mzA, 2*mzA + 1:mzA] = baselineG
    esCondA[1:mzA, 3*mzA + 1:mzA] = baselineQ

    ## If circ is the baseline and star is the treatment, it appears
    ## that the fundamental matrix for U (transitions among the
    ## transient states) is singular BUT a2 (prob. of being successful
    ## in the next time step) is of course zero, so SProbSuccess is
    ## zero --- as it must be if we're using Fcirc.
    if (prod(baselineS == Scirc) == 1) {
      SProbSuccess[numKidsThreshold] = 0
    } else {
      out = makeCondKernel (esCondA, transientStates)
      esQ2Extended = out$q2Extended
      SProbSuccess[numKidsThreshold] = esQ2Extended %*%
        c(AaM0, rep(mzAZero, 3))
    }

    epsilonSurv[numKidsThreshold] =
      (SProbSuccess - epsilonZero)[numKidsThreshold] 

    ## Get main effect of G #################

    esCondA = matrix (0, esmzA, esmzA)
    esCondA[mzA + 1:mzA, 1:mzA] = baselineF
    esCondA[2*mzA + 1:mzA, mzA + 1:mzA] = baselineS
    esCondA[3*mzA + 1:mzA, 2*mzA + 1:mzA] = treatmentG
    esCondA[1:mzA, 3*mzA + 1:mzA] = baselineQ
    
    out = makeCondKernel (esCondA, transientStates)
    esQ2Extended = out$q2Extended

    GProbSuccess[numKidsThreshold] = esQ2Extended %*%
      c(AaM0, rep(mzAZero, 3))

    epsilonGrowth[numKidsThreshold] =
      (GProbSuccess - epsilonZero)[numKidsThreshold]

    ## Get main effect of Q #################

    esCondA = matrix (0, esmzA, esmzA)
    esCondA[mzA + 1:mzA, 1:mzA] = baselineF
    esCondA[2*mzA + 1:mzA, mzA + 1:mzA] = baselineS
    esCondA[3*mzA + 1:mzA, 2*mzA + 1:mzA] = baselineG
    esCondA[1:mzA, 3*mzA + 1:mzA] = treatmentQ
    
    out = makeCondKernel (esCondA, transientStates)
    esQ2Extended = out$q2Extended

    QProbSuccess[numKidsThreshold] = esQ2Extended %*%
      c(AaM0, rep(mzAZero, 3))

    epsilonEnv[numKidsThreshold] =
      (QProbSuccess - epsilonZero)[numKidsThreshold]

    ## Get interaction effects with F #################

    ## F and S
    esCondA = matrix (0, esmzA, esmzA)
    esCondA[mzA + 1:mzA, 1:mzA] = treatmentF
    esCondA[2*mzA + 1:mzA, mzA + 1:mzA] = treatmentS
    esCondA[3*mzA + 1:mzA, 2*mzA + 1:mzA] = baselineG
    esCondA[1:mzA, 3*mzA + 1:mzA] = baselineQ

    out = makeCondKernel (esCondA, transientStates)
    esQ2Extended = out$q2Extended

    FSProbSuccess[numKidsThreshold] = esQ2Extended %*%
      c(AaM0, rep(mzAZero, 3))

    epsilonFecSurv[numKidsThreshold] =
      (FSProbSuccess -
       (epsilonFec + epsilonZero))[numKidsThreshold] 

    ## F and G
    esCondA = matrix (0, esmzA, esmzA)
    esCondA[mzA + 1:mzA, 1:mzA] = treatmentF
    esCondA[2*mzA + 1:mzA, mzA + 1:mzA] = baselineS
    esCondA[3*mzA + 1:mzA, 2*mzA + 1:mzA] = treatmentG
    esCondA[1:mzA, 3*mzA + 1:mzA] = baselineQ

    out = makeCondKernel (esCondA, transientStates)
    esQ2Extended = out$q2Extended

    FGProbSuccess[numKidsThreshold] = esQ2Extended %*%
      c(AaM0, rep(mzAZero, 3))

    epsilonFecGrowth[numKidsThreshold] =
      (FGProbSuccess -
       (epsilonGrowth + epsilonZero))[numKidsThreshold] 

    ## F and Q
    esCondA = matrix (0, esmzA, esmzA)
    esCondA[mzA + 1:mzA, 1:mzA] = treatmentF
    esCondA[2*mzA + 1:mzA, mzA + 1:mzA] = baselineS
    esCondA[3*mzA + 1:mzA, 2*mzA + 1:mzA] = baselineG
    esCondA[1:mzA, 3*mzA + 1:mzA] = treatmentQ

    out = makeCondKernel (esCondA, transientStates)
    esQ2Extended = out$q2Extended

    FQProbSuccess[numKidsThreshold] = esQ2Extended %*%
      c(AaM0, rep(mzAZero, 3))

    epsilonFecEnv[numKidsThreshold] =
      (FQProbSuccess -
       (epsilonEnv + epsilonZero))[numKidsThreshold] 

    ## Get prob(success) with all treatment matrices
    esCondA = matrix (0, esmzA, esmzA)
    esCondA[mzA + 1:mzA, 1:mzA] = treatmentF
    esCondA[2*mzA + 1:mzA, mzA + 1:mzA] = treatmentS
    esCondA[3*mzA + 1:mzA, 2*mzA + 1:mzA] = treatmentG
    esCondA[1:mzA, 3*mzA + 1:mzA] = treatmentQ
    
    out = makeCondKernel (esCondA, transientStates)
    esQ2Extended = out$q2Extended

    allTreatmentProbSuccess[numKidsThreshold] = esQ2Extended %*%
      c(AaM0, rep(mzAZero, 3))
    
    ## Get sum of remaining terms ######################

    remainder[numKidsThreshold] =
      (allTreatmentProbSuccess -
       (epsilonFecSurv + epsilonFecGrowth + epsilonFecEnv +
        epsilonFec + epsilonSurv + epsilonGrowth + epsilonEnv +
        epsilonZero))[numKidsThreshold]
  }  ## end loop over numKidsThreshold

  return (list(epsilonZero=epsilonZero,
               epsilonFec=epsilonFec,
               epsilonSurv=epsilonSurv,
               epsilonGrowth=epsilonGrowth,
               epsilonEnv=epsilonEnv,
               epsilonFecSurv=epsilonFecSurv,
               epsilonFecGrowth=epsilonFecGrowth,
               epsilonFecEnv=epsilonFecEnv,
               remainder=remainder))
}
```
