## Supplementary material for "To prosper, live long: understanding the sources of reproductive skew and extreme reproductive success in structured populations": Associated R code: kestrelLifespanCondLRO.R

```
#########################################################################
##
## kestrelLifespanCondLRO.R: Calculates the probability of having
## lifespan L given LRO R.
##
#########################################################################

rm(list=ls(all=TRUE)); graphics.off();
source ("partitioningUtilities.R")
source ("genericPartitionVarSkewness5.R")
source ("megamatrixFunctions.R")

load ("Falco_naumanni.Rdata")

maxKids = 16
mT = maxKids + 1
maxAge = 40
percentileCutoff = 0.99

## if TRUE, highSurvival increases the survival of all stages by 20%
highSurvival = FALSE

Plist = falco_Umats
Flist = falco_Fmats

if (highSurvival) {
  numEnv = length(Plist)
  for (q in 1:numEnv)
    Plist[[q]][2,] = Plist[[q]][2,]*1.2
  maxAge = 60
}

## Assume all year types are equally likely
numEnv = length(Plist)
Q = matrix (1/numEnv, numEnv, numEnv)
u0 = rep (1/numEnv, numEnv)

## Assume all individuals are born into stage 1
mz = dim(Plist[[1]])[1]
c0 = rep (0, mz)
c0[1] = 1

## Set up cross-classified birth distribution
bigmz = numEnv*mz
m0 = matrix (outer (c0, as.vector(u0)), bigmz, 1)

out = distLifespanCondR2 (Plist, Flist, Q, m0, maxKids,
                         maxAge, percentileCutoff)

distKidsAtDeath = out$distKidsAtDeath
probLifespanCondR = out$probLifespanCondR
sdLifespanCondR = out$sdLifespanCondR
CVLifespanCondR = out$CVLifespanCondR
maxKidsIndex = out$maxKidsIndex
normalSurvProb = out$normalSurvProb

## Subtract 1 to change numKids index into actual number of kids
R50 = min(which(cumsum(distKidsAtDeath) >= 0.5)) - 1
R90 = min(which(cumsum(distKidsAtDeath) >= 0.9)) - 1
R95 = min(which(cumsum(distKidsAtDeath) >= 0.95)) - 1
R99 = min(which(cumsum(distKidsAtDeath) >= 0.99)) - 1

probLifespan = -diff(normalSurvProb)

## Subtract 1 to change age index into age
L50 = min(which(cumsum(probLifespan) >= 0.5)) - 1
L75 = min(which(cumsum(probLifespan) >= 0.75)) - 1
L90 = min(which(cumsum(probLifespan) >= 0.9)) - 1
L95 = min(which(cumsum(probLifespan) >= 0.95)) - 1
L99 = min(which(cumsum(probLifespan) >= 0.99)) - 1

## At what age should we cut off the plots?
##LCutoff = min(which(probLifespanCondR[1,] < 0.001))
LCutoff = L99

if (highSurvival) {
  save (probLifespan, distKidsAtDeath, probLifespanCondR,
        CVLifespanCondR, sdLifespanCondR,
        L50, L75, L90, L95, L99, R50, R90, R95, R99,
        file="kestrelLifespanCondLROHighSurv.Rdata")
} else {
  save (probLifespan, distKidsAtDeath, probLifespanCondR,
        CVLifespanCondR, sdLifespanCondR,
        L50, L75, L90, L95, L99, R50, R90, R95, R99,
        file="kestrelLifespanCondLRO.Rdata")
}
```
