## Supplementary material for "To prosper, live long: understanding the sources of reproductive skew and extreme reproductive success in structured populations": Associated R code: KittiwakeMatrices.R

```
####################################################################
#
# Kittiwake model used by Steiner et al., 2010
# Neutral model for females with cross-classification by age and repr. state
# Parameters come from tables S1, S2, S3, and S4.  
####################################################################

## Ages 1 and 2 stage transition matrix
M1 = diag(5)
M2 = diag(5)
## Stage-specific survival from age 1 to age 2, age 2 to age 3
s1 = rep(1, 5)
s2 = rep(1, 5)

## Age 3 stage transition matrix
M3 = diag (5)
M3[1:5,1] = c(0.785, 0, 0.183, 0.028, 0.004)
## Stage-specific survival from age 3 to age 4
s3 = c(1, 1, 0.694, 1, 0.153)

## Age 4 stage transition matrix
M4 = matrix (0, 5, 5)
M4[1:5,1] = c(0.496, 0, 0.348, 0.129, 0.027)
M4[2,2] = 1
M4[2:5,3] = c(0.378, 0.295, 0.291, 0.036)
M4[2:5,4] = c(0.185, 0.543, 0.206, 0.066)
M4[3:4,5] = c(0.001, 0.999)

## Stage-specific survival for ages 4 and up.  Survival probability
## for immatures was not given.  I set it to 1 to match s3.
#s = c(1, 0.78, 0.78, 0.83, 0.81)
## logit(pars) = c(0.78, 0.78, 0.83, 0.81)
## I.e. pars = "quality" and survival is a logit transform of quality
pars = c(log(0.78/(1 - 0.78)), log(0.78/(1 - 0.78)),
    log(0.83/(1 - 0.83)), log(0.81/(1 - 0.81)))

## Ages 5 and up stage transition matrix
M5 = matrix (0, 5, 5)
M5[1:5,1] = c(0.332, 0, 0.417, 0.210, 0.041)
M5[2:5,2] = c(0.321, 0.404, 0.225, 0.05)
M5[2:5,3] = c(0.17, 0.47, 0.25, 0.11)
M5[2:5,4] = c(0.059, 0.38, 0.397, 0.164)
M5[2:5,5] = c(0.035, 0.326, 0.378, 0.261)

## Age x stage megamatrices
makeP = function (pars) {
  s = c(1, exp(pars)/(1 + exp(pars)))
  P = matrix (0, 25, 25)
  P[5*1 + 1:5, 5*0 + 1:5] = M1 * matrix (rep(s1, 5), 5, 5, byrow=TRUE)
  P[5*2 + 1:5, 5*1 + 1:5] = M2 * matrix (rep(s2, 5), 5, 5, byrow=TRUE)
  P[5*3 + 1:5, 5*2 + 1:5] = M3 * matrix (rep(s3, 5), 5, 5, byrow=TRUE)
  P[5*4 + 1:5, 5*3 + 1:5] = M4 * matrix (rep(s, 5), 5, 5, byrow=TRUE)
  P[5*4 + 1:5, 5*4 + 1:5] = M5 * matrix (rep(s, 5), 5, 5, byrow=TRUE)

  return (P)
}

## Create fecundity matrix.  F[i,j] = the expected number of kids of stage i
## produced by an individual of stage j.

F = matrix (0, 25, 25)
# Stage 5 (F2) individuals produce 2 or 3 fledglings.  In the absence
# of other information, I'm assuming they do each with prob. 1/2.
b = c(0, 0, 0, 1, 2.5)  # mean num. offspring at each stage.  
F[1,] = rep(b, 5)

######################################################################
#  Function to make the B matrix B[i,j] is the probability that a
#  class-j individual has i-1 kids.  Steiner et al. say that a stage 4
#  individual has 1 chick and a stage 5 individual has either 2 or 3,
#  which we assume happens with equal probability.  Recall that in the
#  Age x Stage model, age is the slow variable.
#
#  M = maximum number of kids = 3
#############################################################

mk_B = function (M=3) {
  B = matrix (0, M+1, 25)
  for (i in 1:5) {  # loop over ages
    B[1:(M+1),(i-1)*5 + 1:3] = rep(0, M) # stages 1--3 have no kids
    B[1,(i-1)*5 + 1:3] = 1 # stages 1--3 have no kids
    B[,(i-1)*5 + 4] = c(0, 1, 0, 0) # stage 4 has 1 kid
    B[,(i-1)*5 + 5] = c(0, 0, 0.5, 0.5) # stage 5 has 2 or 3 kids
  }
  return (B)
}

#P=makeP(pars);
#B=mk_B();
#out=make_AxT(B,P,mT=100)

makeTraitVecs = function (numTraits) {

  ## get CV of quality based on survival variation in Fig. 1, Cam et
  ## al. 2002.
  q75 = log(0.9/0.1);   #75th percentile on inverse-logit scale ("quality")
  q25 = log(0.7/0.3)    #25th percentile on inverse-logit scale ("quality")
  IQR=q75-q25;
  sigmaSurv=IQR/1.35;
  muSurv = (q75+q25)/2;
  CVSurv = sigmaSurv/muSurv;

  ## get std. dev. of quality for the quality values for stages 2--5
  ## aged 5+ (and age 4, by the look of it).  (Survival is guaranteed
  ## for stage 1.)
  sdQuality = CVSurv * pars  ## sigma_h for stages 2--5, aged 5+

  ## Let quality range over the middle 95% of its (assumed normal)
  ## distribution
  svec = seq(-1.96, 1.96, length.out=numTraits)
  
  ## vector of associated probabilities
  qualityProb = dnorm(svec,mean=0,sd=1); 
  qualityProb = qualityProb/sum(qualityProb); 
 
  ## matrix of survival quality values
  qualityVals = matrix (0, numTraits, 4)
  for (x in 1:numTraits) 
    qualityVals[x,] = pars + svec[x]*sdQuality

  ## Variation in breeding probabilities #######################

  ## estimate CV of breeding prob. from Cam et al. 2002, fig. 1
  q25 = 0.94
  q75 = 0.98
  
  IQR=q75-q25;
  sigma=IQR/1.35;
  mu = (q75+q25)/2;
  CVBreedingProb = sigma/mu;

  ## If quality h = mu_h + s*sigma_h, then the breeding probability
  ## scaling factor zeta is 1 + ((h - mu_h)/sigma_h) * CVBreedingProb

  breedingProbFactorVals = numeric(numTraits)
  for (x in 1:numTraits)
    breedingProbFactorVals[x] = 1 + svec[x]*CVBreedingProb

  return (out = list(qualityVals=qualityVals,
                     qualityProb=qualityProb,
                     breedingProbFactorVals=breedingProbFactorVals))
}
```
