## Supplementary material for "To prosper, live long: understanding the sources of reproductive skew and extreme reproductive success in structured populations": Associated R code: kittiwakeVarMu3Partition.R

```
#################################################################
##
## kittiwakePartitionVarSuccess: Partitions the (binomial) variance of
## success, where success = meeting or exceeding some threshold LRO.
##
## Original model is an age x stage model, with ages 1, 2, 3, 4, 5+
## and stages
##
## 1: immature
## 2: non-breeder
## 3: failed breeder
## 4: produced 1 chick
## 5: produced 2 or 3 chicks (with equal probability)
##
## The original model had matrices for males and females.  We're using
## the female matrices.
##
## BUG?  Are we sure that the current approach to calculating trait
## contribution is correct?  It seems more reliable than the bstsA
## method, but man would I like them to agree.
##
## This code is almost identical to kittiwakeVarSkewnessPartition.R,
## but does the partitioning of the 3rd central moment of LRO, not the
## skewness.  See that code for detailed comments.
##
###################################################################

rm(list=ls(all=TRUE)); graphics.off(); 
require(Matrix); require(statmod);

source ("kittiwake/KittiwakeMatrices.R")
source("Utilities.R") 
source("megamatrixFunctions.R");
source ("partitioningUtilities.R")
source ("genericPartitionVarSkewness6.R")

lineTypes = c(1,2,3,4)
names(lineTypes) = c("surv", "env", "growth", "fec")

colors = c("red", "green", "orange", "blue")
names(colors) = c("surv", "env", "growth", "fec")

debugging = TRUE

## How finely are we going to slice up the trait distribution?
numTraits = 50
##numTraits = 10

## Maximum lifespan
maxAge = 140

## Maximum clutch size
maxClutch = 3

## Maximum #kids in a lifetime
maxKids = 20
mT = maxKids + 1

## number of stages within one age
mz = 5
## number of age classes
numAge = 5
## number of states (#age classes x #stage classes)
bigmz = numAge*mz

graphics.off(); 

## birth age x stage distribution
c0 = rep(0, bigmz); c0[1] = 1

## Mean number of offspring at each stage
##b = rep(c(0,0,0,1,1.5), 5)

## set up trait distribution
out = makeTraitVecs (numTraits)
qualityVals = out$qualityVals
qualityProb = out$qualityProb
breedingProbFactorVals = out$breedingProbFactorVals

## Allocate some memory
survTrajecVarCondX = growthTrajecVarCondX = 
  fecVarCondX = matrix (0, numTraits, maxAge)
survTrajecMu3CondX = growthTrajecMu3CondX = 
  fecMu3CondX = matrix (0, numTraits, maxAge)
reproArrivalMode = lifespan = numeric (numTraits)

totSurvTrajecVarCondX = totGrowthTrajecVarCondX =
  totFecVarCondX = numeric (numTraits)

totSurvTrajecMu3CondX = totGrowthTrajecMu3CondX =
  totFecMu3CondX = numeric (numTraits)

expRCondX = totVarCondX = totMu3CondX = totMu3CondX =
  numeric (numTraits)
Plist = list ()

esbigmz = 3*bigmz
bsbigmz = 1 + bigmz

esF = esR1 = esR2 = esR3 = matrix (0, esbigmz+1, esbigmz+1)
bsF = bsR1 = bsR2 = bsR3 = matrix (0, bsbigmz+1, bsbigmz+1)

## define reward matrices.  (F doesn't depend on x.)
esF[bigmz + 1:bigmz, 1:bigmz] = F
bsF[1, 1 + 1:bigmz] = colSums (F)

for (j in 1:esbigmz) 
  esR1[,j] = sum(esF[,j])

esMultichickStages = seq (5, 25, 5)
esR2 = esR1^2
esR2[,esMultichickStages] = 13/2
esR3 = esR1^3
esR3[,esMultichickStages] = 35/2

for (j in 1:bsbigmz) 
  bsR1[,j] = sum(bsF[,j])
bsMultichickStages = 1 + seq(5, 25, 5)
bsR2 = bsR1^2
bsR2[,bsMultichickStages] = 13/2
bsR3 = bsR1^3
bsR3[,bsMultichickStages] = 35/2

## Loop over traits
for (x in 1:numTraits) {
  cat ("x = ", x, "\n")
  newpars = qualityVals[x,]
  breedingProbFactor = breedingProbFactorVals[x]

  ## Modify stage transition matrices (M1 for age 1, M2 for age 2,
  ## etc.)  We have decided to let the 5+ year olds be the ones where
  ## traits vary.
  newM3 = M3; newM4 = M4; newM5 = M5; 
  
  newM5[4:5,] = newM5[4:5,] * breedingProbFactor
  for (j in 1:5) {
    b = sum(newM5[4:5,j]); norm = sum(newM5[1:3,j]); bfac = (1-b)/norm; 
    newM5[1:3,j]=newM5[1:3,j]*bfac; 
  }

  ## Make transition matrix P
  s = c(1, exp(newpars)/(1 + exp(newpars)))
  P = matrix (0, 25, 25)
  P[5*1 + 1:5, 5*0 + 1:5] = M1 * matrix (rep(s1, 5), 5, 5, byrow=TRUE)
  P[5*2 + 1:5, 5*1 + 1:5] = M2 * matrix (rep(s2, 5), 5, 5, byrow=TRUE)
  P[5*3 + 1:5, 5*2 + 1:5] = newM3 * matrix (rep(s3, 5), 5, 5, byrow=TRUE)
  P[5*4 + 1:5, 5*3 + 1:5] = newM4 * matrix (rep(s, 5), 5, 5, byrow=TRUE)
  P[5*4 + 1:5, 5*4 + 1:5] = newM5 * matrix (rep(s, 5), 5, 5,
                                            byrow=TRUE)

  Plist[[x]] = P
  
  out = getVarMu3PartitionsNoEnvVar (P, F, c0, maxAge,
                                          esR1, esR2, esR3,
                                          bsR1, bsR2, bsR3)

  survTrajecMu3CondX[x,] = out$survTrajecMu3
  growthTrajecMu3CondX[x,] = out$growthTrajecMu3
  fecMu3CondX[x,] = out$fecMu3
  totMu3CondX[x] = out$totMu3
  lifespan[x] = out$lifespan

  survTrajecVarCondX[x,] = out$survTrajecVar
  growthTrajecVarCondX[x,] = out$growthTrajecVar
  fecVarCondX[x,] = out$fecVar
  totVarCondX[x] = out$totVar
  totVarCondX2 = 
    sum(survTrajecVarCondX[x,] + growthTrajecVarCondX[x,] +
        fecVarCondX[x,])


  ## Get expected LRO while you're at it.
  N = solve (diag(bigmz) - P)
  expRCondX[x] = (colSums (F %*% N)) %*% c0

  ## And get modal time to reproductive state
  absorbingStates = (0:4)*mz + 4
  absorbingStates = c(absorbingStates, (0:4)*mz + 5)

  out = getModalTimeToHitState (absorbingStates, Plist, Flist,
                                Q=NULL, c0, maxAge)
  reproArrivalMode[x] = out$hitTimeMode

  ## Sanity check on variance.  Passes, although as x approaches its
  ## highest values, totVarCondX2 disagrees a little with totVarCondX
  ## (obtained from bsP in getVarMu3PartitionsNoEnvVar ()) and
  ## expZVarRCondXZ + varZExpRCondXZ, which always agree.
  if (debugging) {
    expRCondXZ = colSums (F %*% N)
    sigbsq = c(0, 0, 0, 0, 0.25)
    pb = c(0, 0, 0, 1, 1) # prob. of breeding
    b = c(0, 0, 0, 1, 2.5)  # mean num. offspring at each stage.  
    rbarPib = expRCondXZ %*% P     # \bar{r} \pi_b
    r2 = (sigbsq + (pb*b)^2 + 2*pb*b*rbarPib) %*% N
    varRCondXZ = r2 - expRCondXZ^2
    expZVarRCondXZ = varRCondXZ %*% c0
    varZExpRCondXZ = sum(c0*expRCondXZ^2) - sum(c0*expRCondXZ)^2
    cat ("Var. check:", totVarCondX[x], "should = ",
         expZVarRCondXZ + varZExpRCondXZ, 
         "should = ", totVarCondX2, "\n")
  }
}

totSurvTrajecMu3CondX = apply (survTrajecMu3CondX, 1, sum)
totGrowthTrajecMu3CondX = apply (growthTrajecMu3CondX, 1, sum)
totFecMu3CondX = apply (fecMu3CondX, 1, sum)

totSurvTrajecVarCondX = apply (survTrajecVarCondX, 1, sum)
totGrowthTrajecVarCondX = apply (growthTrajecVarCondX, 1, sum)
totFecVarCondX = apply (fecVarCondX, 1, sum)

traitAve = function (x) {
  return (sum(qualityProb*x))
}

## Average over traits
fecMu3 = apply (fecMu3CondX, 2, traitAve)
survTrajecMu3 = apply (survTrajecMu3CondX, 2, traitAve)
growthTrajecMu3 = apply (growthTrajecMu3CondX, 2, traitAve)

fecVar = apply (fecVarCondX, 2, traitAve)
survTrajecVar = apply (survTrajecVarCondX, 2, traitAve)
growthTrajecVar = apply (growthTrajecVarCondX, 2, traitAve)

totSurvTrajecMu3 = sum (survTrajecMu3)
totGrowthTrajecMu3 = sum (growthTrajecMu3)
totFecMu3 = sum (fecMu3)

totSurvTrajecVar = sum (survTrajecVar)
totGrowthTrajecVar = sum (growthTrajecVar)
totFecVar = sum (fecVar)


aveLifespan = traitAve (lifespan)
aveReproArrivalMode = traitAve (reproArrivalMode)


######################################################################
## To get total skewness or variance, we need to create transition
## matrices and reward matrices for a model in which state is
## classified by trait x stage.  Usually, we need trait x stage x
## env., so in keeping with, e.g., the fulmar code, I'll use the
## prefix "ts" to suggest "triple state" as a way to distinguish these
## matrices.
##
## And since we need to start off in the ur-state, we need to include
## pre-birth states (the ur-trait, ur-stage), so the full prefix is
## bsts...
##
## A guide to index ranges: 1 = the ur state.  2--(1 + numTraits):
## trait is assigned but nothing else.
## (numTraits+2)--(numTraits+1 + numTraits*bigmz): trait and z are
## assigned.  (numTraits+2)--(numTraits+1+bigmz): trait 1, z.  Etc.
##
#######################################################################

tsbigmz = numTraits * bigmz
bstsbigmz = 1 + numTraits + tsbigmz

tsF = matrix (0, bstsbigmz, bstsbigmz)
tsP = as.matrix (bdiag (Plist))
for (x in 1:numTraits)
  tsF[(x-1)*bigmz + 1:bigmz, (x-1)*bigmz + 1:bigmz] = F

bstsF = bstsP = matrix (0, bstsbigmz, bstsbigmz)
## From ur-stage to just traits
bstsP[1 + (1:numTraits),1] = qualityProb
## From just traits to traits and stage
for (x in 1:numTraits)
  bstsP[1 + numTraits + (x-1)*bigmz + 1:bigmz, 1 + x] = c0
## From normal stage state to normal stage state
bstsP[1 + numTraits + 1:tsbigmz,
      1 + numTraits + 1:tsbigmz] = tsP

## You only start reproducing once you're past the ur-stages.  I guess
## reproduction should go to the ur-stage state.  Not that it matters
## much, since all we need are the column sums of bsF.
bstsF[1, 1 + numTraits + 1:tsbigmz] =
  rep (colSums (F), numTraits)

## set up bstsPplus, the transition matrix with absorbing state (dead)
bstsPplus = matrix (0, bstsbigmz+1, bstsbigmz+1)
bstsPplus[1:bstsbigmz, 1:bstsbigmz] = bstsP
bstsPplus[bstsbigmz+1, 1:bstsbigmz] = 1 - colSums(bstsP)
bstsPplus[bstsbigmz+1, bstsbigmz+1] = 1

## Set up bstsZ, which cleaves off dead states
bstsZ = cbind (diag(bstsbigmz), rep(0, bstsbigmz))

## Set up the vector 1_s, where s = bigmz + 1
bstsOneS = rep(0, bstsbigmz+1)
bstsOneS[bstsbigmz+1] = 1

## set up reward matrices
bstsR1 = bstsR2 = bstsR3 = matrix (0, bstsbigmz+1, bstsbigmz+1)
bstsR1Stripped = bstsR2Stripped = matrix (0, bstsbigmz, bstsbigmz)

foo = colSums(bstsF)
for (j in 1:bstsbigmz)
  bstsR1[,j] = foo[j]

bstsMultichickStages = 1 + numTraits + seq(5, numTraits*25, 5)
bstsR2 = bstsR1^2
bstsR2[,bstsMultichickStages] = 13/2
bstsR3 = bstsR1^3
bstsR3[,bstsMultichickStages] = 35/2

bstsR1Stripped = bstsR1[1:bstsbigmz,1:bstsbigmz]
##bstsR2Stripped = bstsR1Stripped + bstsR1Stripped^2
bstsR2Stripped = bstsR2[1:bstsbigmz,1:bstsbigmz]

## Ex(R)
bstse = rep (1, bstsbigmz+1)
bstsN = solve (diag(bstsbigmz) - bstsP)
bstsRho1Vec = t(bstsN) %*% bstsZ %*% t(bstsPplus * bstsR1) %*% bstse

## Ex(R^2 | x)
bstsRho2Vec = t(bstsN) %*% (bstsZ %*% t(bstsPplus * bstsR2) %*% bstse +
                        2*t(bstsP * bstsR1Stripped) %*% bstsRho1Vec)

## Ex(R^3 | x)
bstsRho3Vec = t(bstsN) %*% (bstsZ %*% t(bstsPplus * bstsR3) %*% bstse +
                        3*t(bstsP*bstsR2Stripped) %*% bstsRho1Vec +
                        3*t(bstsP*bstsR1Stripped) %*% bstsRho2Vec)

bstsRho1Vec = t(bstsRho1Vec)
bstsRho2Vec = t(bstsRho2Vec)
bstsRho3Vec = t(bstsRho3Vec)

bstsMu2Vec = bstsRho2Vec - bstsRho1Vec^2
bstsMu3Vec = bstsRho3Vec - 3*bstsRho1Vec*bstsRho2Vec + 2*bstsRho1Vec^3

urState = rep(0, bstsbigmz)
urState[1] = 1
totMu3 = bstsMu3Vec %*% urState
totVar = bstsMu2Vec %*% urState

traitMu3 = totMu3 - traitAve (totMu3CondX)
## Note: 2018 paper says that traits make up 38.9% of the variance.
## (Here it's 22.4%.)  But I'm not sure we were handling trait
## variation the same way.
traitVar = totVar - traitAve (totVarCondX)

## Use the law of total variance to get the full variance another
## way.  

## Variance
expXVarRCondX = traitAve (totVarCondX)
varXExpRCondX = sum(qualityProb*expRCondX^2) - traitAve (expRCondX)^2
totVar2 = expXVarRCondX + varXExpRCondX

## sanity check: passes!
cat ("Checking total variance:", totVar, "should = ", totVar2, "\n")


par(cex.lab=1.4, lwd=2)

## Average over traits and include contribution to skewness from
## traits
ymax = max(survTrajecMu3, growthTrajecMu3,
           fecMu3)
ymin = min(survTrajecMu3, growthTrajecMu3,
           fecMu3) 
plot(0:40, survTrajecMu3[1:41], col=colors["surv"],
     lty=lineTypes["surv"],
     xlab="Age", ylab="Contributions to Mu3(R)", type="l",
     ylim=c(ymin, ymax))
lines (0:40, growthTrajecMu3[1:41], col=colors["growth"],
      lty=lineTypes["growth"])
lines (rep(aveLifespan, 2), c(0, 0.35*ymax), col="black",
       lty=1, lwd=2)
lines (rep(aveReproArrivalMode, 2), c(0, ymax), col="purple",
       lty=1, lwd=2)
lines (0:40, fecMu3[1:41], col=colors["fec"],
      lty=lineTypes["fec"])

ydiff = ymax - ymin

legend (x=17, y=0.94*ymax, legend=c("99% of individ. are dead",
                                     "Mode(age of 1st repro.)"),
        col=c("black", "purple"),
        lty=1, bty="n", lwd=2, cex=1.2)

legend (x=17, y=0.81*ymax,
        legend=c(paste("Surv trajec. luck: ",
                       round(100*totSurvTrajecMu3/totMu3, digits=1),
                       "%", sep=""),
                 paste("Growth trajec. luck: ",
                       round(100*totGrowthTrajecMu3/totMu3, digits=1),
                       "%", sep=""),
                 paste("Fecundity luck: ",
                       round(100*totFecMu3/totMu3, digits=1),
                       "%", sep="")),
        col=colors[-2], lty=lineTypes[-2], bty="n", lwd=2, cex=1.2)

text (x=26, y=ymin+0.54*ydiff, col="black", cex=1.2,
      paste ("Fixed traits: ",
             round (100*traitMu3/totMu3, digits=1),
             "%", sep=""))

dev.copy2pdf (file="kittiwakeMu3Partition.pdf")
```
