## Supplementary material for "To prosper, live long: understanding the sources of reproductive skew and extreme reproductive success in structured populations": Associated R code: makeP.R

```
#################################################################
##
## makeP: makes transition matrices for Lomatium bradshawii for the
## two Rose Prairie locations.  There are 7 years for each location.
##
##################################################################

location = 1  ## choose location 1 or 2
numEnv = 7
mz = 5
bigmz = mz*numEnv

if (location == 1) {
  d = read.table ("RosePrairie1.dat", header=FALSE)
} else if (location == 2) {
  d = read.table ("RosePrairie2.dat", header=FALSE)
}

Flist = Plist = list (numEnv)
for (q in 1:numEnv) {
  foo = matrix (d[,q], 5, 5)
  Plist[[q]] = Flist[[q]] = matrix (0, 5, 5)
  Flist[[q]][1,4:5] = foo[1,4:5]
  Plist[[q]][2:5,] = foo[2:5,]
  Plist[[q]][1,1:3] = foo[1,1:3]
}
```
