## Supplementary material for "To prosper, live long: understanding the sources of reproductive skew and extreme reproductive success in structured populations": Associated R code: lomatiumSkewnessPartition5.R

```
#################################################################
##
## lomatiumThirdMomentAgePartition5: uses
## genericPartitionVarSkewness2.R, our now-canonical partitioning
## code.
##
#################################################################

rm(list=ls(all=TRUE)); graphics.off(); 
require(Matrix); require(statmod);

demogDir = "./Lomatium/"
if(.Platform$OS.type=="windows") {
  demogDir = "c:/repos/fanova/envVar/Lomatium"
  codeDir = "c:/repos/fanova/envVar"
}

## Make P and Q matrices
setwd (demogDir)
source ("makeP.R")
setwd ("../")

source ("Utilities.R")
source ("partitioningUtilities.R")
source ("genericPartitionVarSkewness6.R")
source ("megamatrixFunctions.R")

lineTypes = c(1,2,3,4)
names(lineTypes) = c("surv", "env", "growth", "fec")

colors = c("red", "green", "orange", "blue")
names(colors) = c("surv", "env", "growth", "fec")

onlyEnv1 = FALSE
stickyEnv = FALSE ## positively autocorrelated env.

### really maxAge+1, because we let age start at 0 
maxAge = 100

surv = envTrajec = survTrajec = growthTrajec = rep (0, maxAge)

## birth size distribution
c0 = rep (0, mz); c0[1] = 1

if (onlyEnv1) {
  Q = matrix (0, numEnv, numEnv)
  Q[1,1] = 1
  u0 = c(1, rep(0, numEnv-1))
} else {
  ## Specify probability of fire (good) versus bad years
  ## And draw randomly from each subset of the data
  Q = matrix (0, numEnv, numEnv)
  pf=0.333; qf = 1-pf; 
  into = c(pf/4,pf/4,qf/3,pf/4,pf/4,qf/3,qf/3);
  for(j in 1:7) Q[,j]=into;
  u0 = domEig(Q)$w
}

if (stickyEnv) {s = 0.8; Q = diag(s, numEnv) + (1 - s)*Q}

## The initial cross-classified distribution
m0 = matrix (outer (c0, as.vector(u0)), bigmz, 1)

out = getVarSkewnessPartitionsEnvVar (Plist, Flist, Q, m0, maxAge)

birthEnvSkewness = out$birthEnvSkewness
survTrajecSkewness = out$survTrajecSkewness
growthTrajecSkewness = out$growthTrajecSkewness
envTrajecSkewness = out$envTrajecSkewness
fecSkewness = out$fecSkewnes
totSkewness = out$totSkewness
lifespan = out$lifespan

totSurvTrajecSkewness = sum(survTrajecSkewness)
totGrowthTrajecSkewness = sum(growthTrajecSkewness)
totEnvTrajecSkewness = sum(envTrajecSkewness)
totFecSkewness = sum(fecSkewness)

birthEnvVar = out$birthEnvVar
survTrajecVar = out$survTrajecVar
growthTrajecVar = out$growthTrajecVar
envTrajecVar = out$envTrajecVar
fecVar = out$fecVar
totVar = out$totVar

totSurvTrajecVar = sum(survTrajecVar)
totGrowthTrajecVar = sum(growthTrajecVar)
totEnvTrajecVar = sum(envTrajecVar)
totFecVar = sum(fecVar)

out = getModalTimeToFirstRepro (Plist, Flist, Q, m0)
reproArrivalMode = out$reproArrivalMode

## Plot skewness

par (cex.lab=1.4, lwd=2)
ymax = max(survTrajecSkewness, growthTrajecSkewness,
           envTrajecSkewness)
ymin = min(survTrajecSkewness, growthTrajecSkewness, envTrajecSkewness)
plot (0:15, survTrajecSkewness[1:16], col=colors["surv"],
      lty=lineTypes["surv"],
      xlab="Age", ylab="Contributions to Skewness(R)", type="l",
      ylim=c(ymin, ymax))
lines (0:15, growthTrajecSkewness[1:16], col=colors["growth"],
      lty=lineTypes["growth"])
lines (0:15, envTrajecSkewness[1:16], col=colors["env"],
      lty=lineTypes["env"])
lines (0:15, fecSkewness[1:16], col=colors["fec"],
      lty=lineTypes["fec"])
lines (rep(lifespan, 2), c(0, 0.35*ymax), col="black",
       lty=1, lwd=2)
lines (rep(reproArrivalMode, 2), c(0, ymax), col="purple",
       lty=1, lwd=2)

xloc = 0.3*15

legend (x=xloc, y=0.98*ymax, legend=c("99% of individ. are dead",
                                     "Mode(age of 1st repro.)"),
        col=c("black", "purple"),
        lty=1, bty="n", lwd=2, cex=1.2)

legend (x=xloc, y=0.78*ymax,
        legend=c(paste("Surv. trajec. luck: ",
                       round(100*totSurvTrajecSkewness/totSkewness, digits=1),
                       "%", sep=""),
                 paste("Env. trajec. luck: ",
                       round(100*totEnvTrajecSkewness/totSkewness, digits=1),
                       "%", sep=""),
                 paste("Growth trajec. luck: ",
                       round(100*totGrowthTrajecSkewness/totSkewness, digits=1),
                       "%", sep=""),
                 paste("Fecundity luck: ",
                       round(100*totFecSkewness/totSkewness, digits=1),
                       "%", sep="")),
        col=colors, lty=lineTypes, bty="n", lwd=2, cex=1.2)


text (x=8.5, y=0.41*ymax, cex=1.2,
      paste("Birth env.: ",
            round(100*birthEnvSkewness/totSkewness, digits=1),
            "%", sep=""), col="black")

dev.copy2pdf (file="lomatiumSkewnessPartition.pdf")

## And plot variance
dev.new ()
par (cex.lab=1.4, lwd=2)
ymax = max(survTrajecVar, growthTrajecVar,
           envTrajecVar)
ymin = min(survTrajecVar, growthTrajecVar, envTrajecVar)
plot (0:15, survTrajecVar[1:16], col=colors["surv"],
      lty=lineTypes["surv"],
      xlab="Age", ylab="Contributions to Var(R)", type="l",
      ylim=c(ymin, ymax))
lines (0:15, growthTrajecVar[1:16], col=colors["growth"],
      lty=lineTypes["growth"])
lines (0:15, envTrajecVar[1:16], col=colors["env"],
      lty=lineTypes["env"])
lines (0:15, fecVar[1:16], col=colors["fec"],
      lty=lineTypes["fec"])
lines (rep(lifespan, 2), c(0, 0.35*ymax), col="black",
       lty=1, lwd=2)
lines (rep(reproArrivalMode, 2), c(0, ymax), col="purple",
       lty=1, lwd=2)

xloc = 0.45*15

legend (x = xloc, y=0.98*ymax, legend=c("99% of individ. are dead",
                                     "Mode(age of 1st repro.)"),
        col=c("black", "purple"),
        lty=1, bty="n", lwd=2, cex=1.2)

legend (x=xloc, y=0.8*ymax,
        legend=c(paste("Surv. trajec. luck: ",
                       round(100*totSurvTrajecVar/totVar, digits=1),
                       "%", sep=""),
                 paste("Env. trajec. luck: ",
                       round(100*totEnvTrajecVar/totVar, digits=1),
                       "%", sep=""),
                 paste("Growth trajec. luck: ",
                       round(100*totGrowthTrajecVar/totVar, digits=1),
                       "%", sep=""),
                 paste("Fecundity luck: ",
                       round(100*totFecVar/totVar, digits=1),
                       "%", sep="")),
        col=colors, lty=lineTypes, bty="n", lwd=2, cex=1.2)

text (x=10.6, y=0.46*ymax, cex=1.2,
      paste("Birth env.: ",
            round(100*birthEnvVar/totVar, digits=1),
            "%", sep=""), col="black")


dev.copy2pdf (file="lomatiumVarPartition.pdf")
```
