## Supplementary material for "To prosper, live long: understanding the sources of reproductive skew and extreme reproductive success in structured populations": Associated R code: lomatiumVarMu3Partition.R

```
#################################################################
##
## lomatiumThirdMomentAgePartition5: uses
## genericPartitionVarSkewness2.R, our now-canonical partitioning
## code.
##
## BUG:  lomatiumAgePartition2.R in ../fullDistLuck uses something
## different for Q.
##
#################################################################

rm(list=ls(all=TRUE)); graphics.off(); 
require(Matrix); require(statmod);

demogDir = "./Lomatium/"
if(.Platform$OS.type=="windows") {
  demogDir = "c:/repos/fanova/envVar/Lomatium"
  codeDir = "c:/repos/fanova/envVar"
}

## Make P and Q matrices
setwd (demogDir)
source ("makeP.R")
setwd ("../")

source ("Utilities.R")
source ("partitioningUtilities.R")
source ("genericPartitionVarSkewness6.R")
source ("megamatrixFunctions.R")

lineTypes = c(1,2,3,4)
names(lineTypes) = c("surv", "env", "growth", "fec")

colors = c("red", "green", "orange", "blue")
names(colors) = c("surv", "env", "growth", "fec")

### really maxAge+1, because we let age start at 0 
maxAge = 100

surv = envTrajec = survTrajec = growthTrajec = rep (0, maxAge)

## birth size distribution
c0 = rep (0, mz); c0[1] = 1

## Specify probability of fire (good) versus bad years
## And draw randomly from each subset of the data
Q = matrix (0, numEnv, numEnv)
pf=0.333; qf = 1-pf; 
into = c(pf/4,pf/4,qf/3,pf/4,pf/4,qf/3,qf/3);
for(j in 1:7) Q[,j]=into;
u0 = domEig(Q)$w

## The initial cross-classified distribution
m0 = matrix (outer (c0, as.vector(u0)), bigmz, 1)

out = getVarMu3PartitionsEnvVar (Plist, Flist, Q, m0, maxAge)

birthEnvMu3 = out$birthEnvMu3
survTrajecMu3 = out$survTrajecMu3
growthTrajecMu3 = out$growthTrajecMu3
envTrajecMu3 = out$envTrajecMu3
fecMu3 = out$fecMu3
totMu3 = out$totMu3
lifespan = out$lifespan

totSurvTrajecMu3 = sum(survTrajecMu3)
totGrowthTrajecMu3 = sum(growthTrajecMu3)
totEnvTrajecMu3 = sum(envTrajecMu3)
totFecMu3 = sum(fecMu3)

birthEnvVar = out$birthEnvVar
survTrajecVar = out$survTrajecVar
growthTrajecVar = out$growthTrajecVar
envTrajecVar = out$envTrajecVar
fecVar = out$fecVar
totVar = out$totVar

totSurvTrajecVar = sum(survTrajecVar)
totGrowthTrajecVar = sum(growthTrajecVar)
totEnvTrajecVar = sum(envTrajecVar)
totFecVar = sum(fecVar)

out = getModalTimeToFirstRepro (Plist, Flist, Q, m0)
reproArrivalMode = out$reproArrivalMode

## Plot skewness

par (cex.lab=1.4, lwd=2)
ymax = max(survTrajecMu3, growthTrajecMu3,
           envTrajecMu3)
ymin = min(survTrajecMu3, growthTrajecMu3, envTrajecMu3)
plot (0:15, survTrajecMu3[1:16], col=colors["surv"],
      lty=lineTypes["surv"],
      xlab="Age", ylab="Contributions to Mu3(R)", type="l",
      ylim=c(ymin, ymax))
lines (0:15, growthTrajecMu3[1:16], col=colors["growth"],
      lty=lineTypes["growth"])
lines (0:15, envTrajecMu3[1:16], col=colors["env"],
      lty=lineTypes["env"])
lines (0:15, fecMu3[1:16], col=colors["fec"],
      lty=lineTypes["fec"])
lines (rep(lifespan, 2), c(0, 0.3*ymax), col="black",
       lty=1, lwd=2)
lines (rep(reproArrivalMode, 2), c(0, ymax), col="purple",
       lty=1, lwd=2)

xloc = 0.4*15

legend (x=xloc, y=0.98*ymax, legend=c("99% of individ. are dead",
                                     "Mode(age of 1st repro.)"),
        col=c("black", "purple"),
        lty=1, bty="n", lwd=2, cex=1.2)

legend (x=xloc, y=0.79*ymax,
        legend=c(paste("Surv. trajec. luck: ",
                       round(100*totSurvTrajecMu3/totMu3, digits=1),
                       "%", sep=""),
                 paste("Env. trajec. luck: ",
                       round(100*totEnvTrajecMu3/totMu3, digits=1),
                       "%", sep=""),
                 paste("Growth trajec. luck: ",
                       round(100*totGrowthTrajecMu3/totMu3, digits=1),
                       "%", sep=""),
                 paste("Fecundity luck: ",
                       round(100*totFecMu3/totMu3, digits=1),
                       "%", sep="")),
        col=colors, lty=lineTypes, bty="n", lwd=2, cex=1.2)

text (x=10, y=0.44*ymax, cex=1.2,
      paste("Birth env.: ",
            round(100*birthEnvMu3/totMu3, digits=1),
            "%", sep=""), col="black")

dev.copy2pdf (file="lomatiumMu3Partition.pdf")

if (FALSE) {
## And plot variance
dev.new ()
par (cex.lab=1.4, lwd=2)
ymax = max(survTrajecVar, growthTrajecVar,
           envTrajecVar)
ymin = min(survTrajecVar, growthTrajecVar, envTrajecVar)
plot (0:15, survTrajecVar[1:16], col="black",
      xlab="Age", ylab="Var contributions", type="l",
      ylim=c(ymin, ymax))
lines (0:15, growthTrajecVar[1:16], col="orange")
lines (0:15, envTrajecVar[1:16], col="green")
lines (0:15, fecVar[1:16], col="blue")
lines (rep(lifespan, 2), c(0, 0.3*ymax), col="black",
       lty=2, lwd=2)
if (FALSE) {
  legend (x="topright", legend=c("Surv. trajec. variance",
                                 "Growth trajec. variance",
                                 "Env. trajec. variance",
                                 "Fec. variance",
                                 "90% of individ. are dead"),
          col=c("black", "orange", "green", "blue", "black"),
          lty=c(1,1,1,1,1,2), bty="n", lwd=2)
} else {
  legend (x=7, y=0.014, legend=c("99% of individ. are dead"),
          col=c("black", "orange", "green", "blue", "black"),
          lty=2, bty="n", lwd=2, cex=1.2)
}

xloc = 0.75*15

text (x=xloc, y=0.67*ymax, cex=1.2,
      paste("Surv trajec.: ",
            round(100*totSurvTrajecVar/totVar, digits=1),
            "%", sep=""), col="black")

text (x=xloc, y=0.6*ymax, cex=1.2,
      paste("Env. trajec.: ",
            round(100*totEnvTrajecVar/totVar, digits=1),
            "%", sep=""), col="green")

text (x=xloc, y=0.53*ymax, cex=1.2,
      paste("Growth trajec.: ",
            round(100*totGrowthTrajecVar/totVar, digits=1),
            "%", sep=""), col="orange")

text (x=xloc, y=0.46*ymax, cex=1.2,
      paste("Fecundity: ",
            round(100*totFecVar/totVar, digits=1),
            "%", sep=""), col="blue")

text (x=xloc, y=0.39*ymax, cex=1.2,
      paste("Birth env.: ",
            round(100*birthEnvVar/totVar, digits=1),
            "%", sep=""), col="black")


##dev.copy2pdf (file="lomatiumVarPartition.pdf")
}
```
