## Supplementary material for "To prosper, live long: understanding the sources of reproductive skew and extreme reproductive success in structured populations": Associated R code: megamatrixFunctions.R

```
######################################################################
#  Function to make the B matrix B[i,j] is the probability that a
#  class-j individual has i-1 kids.  We assume Poisson-distributed
#  number of offspring.
#
#  maxKids = maximum number of kids = 20 by default
#############################################################

mk_B = function (maxKids=20, F) {
  bigmz = ncol(F)
  B = matrix (0, maxKids+1, bigmz)
  for (z in 1:bigmz) 
    B[,z] = dpois (0:maxKids, lambda=sum(F[,z]))

  return (B)
}

######################################################################
# Function to take B and M matrices, and compute the transition 
# probabilities from size-class i and j total kids, to all size classes
# and l total kids. This returns a vector of zeros if (l-j) is < 0
# or above the assumed maxnumber of kids per year,  ncol(B) - 1. 
######################################################################
p_xT <- function(l, i, j, B, M) {
  bigmz <- ncol(M); maxKids <- nrow(B)-1;
  newKids <- (l-j); 
  if((newKids < 0) | (newKids > maxKids)) {
    return(rep(0, bigmz))
  }else{
    return(M[,i]*B[newKids+1,i])
  }
}

##############################################################################
# Function to make the 2D iteration matrix A for a size-kids model
# based on the M and B matrices summarizing a size-structured
# IPM. Apart from B and M the only input is mT, dimension for T 
# (so range of T is 0 to mT-1). The 4-D iteration array K is also returned. 
#
# Iteration matrix is modifited so individuals who get to the maximum
# values of T in the matrix stay there, but continue to grow/shrink/die
#############################################################################

make_AxT <- function(B, M, mT) {
  bigmz=ncol(M); Kvals=array(0,c(bigmz,mT,bigmz,mT));  
  for(z in 1:bigmz){ # initial size
    for(k in 1:mT){ # initial T 
      for(kp in 1:(mT-1)){ # final T
        Kvals[,kp,z,k]=p_xT(kp,z,k,B,M)
      }
      for (zp in 1:bigmz)
        ## make last kids class absorbing
        Kvals[zp,mT,z,k] = M[zp,z] - sum(Kvals[zp,,z,k])
    }
  }
  ## make kids-class mT absorbing: stay there with prob=1
  Kvals[1:bigmz,1:mT,1:bigmz,mT] <- 0; 
  Kvals[1:bigmz,mT,1:bigmz,mT] <- M; 
  A <- Kvals; dim(A) <- c(bigmz*mT,bigmz*mT); 

  return(list(A=A,K=Kvals)) 
}  
```
