## Supplementary material for "To prosper, live long: understanding the sources of reproductive skew and extreme reproductive success in structured populations": Associated R code: partitioningUtilities.R

```
#############################################################################
##
## partitioningUtilities: Utility functions for partitioning variance
## and skewness, with the full kernel and with a conditional kernel.
##
#############################################################################

## Calculate probability of success conditional on initial state and
## kernel conditional becoming successful before death.  If you are
## unfamiliar with this calculation, you can find it detailed in
## Snyder and Ellner, Am. Nat. 188(2), E28--E45, 2016 (see sections
## "Probability of becoming lucky" and "Comparing life trajectories")
## or in Kemeny, J. G., and J. L. Snell. 1960. Finite Markov chains. Van
## Nostrand, New York.

makeCondKernel = function (M, transientStates) {

  ## How many states are there all told?
  numStates = dim(M)[1]
  ## Create U, the transition matrix for the transient states
  U = M[transientStates, transientStates]
  ## How many transient states are there?
  numTransient = length(transientStates)
  ## calculate a2, the probability of hitting an absorbing state in
  ## the next time step
  a2 = apply (M, 2, sum)[transientStates] - apply (U, 2, sum)
  ## calculate q2, the probability of being in Z2 before you die given
  ## that you are now in state i.
  NU = solve (diag(numTransient) - U)
  q2 = a2 %*% NU
  q2Extended = rep (1, numStates); q2Extended[transientStates] = q2

  ## Transition matrix conditional on reproducing before death.
  MCond = matrix (0, numStates, numStates)
  for (j in 1:numStates)
    MCond[,j] = M[,j] * q2Extended / q2Extended[j]
    
  return (out=list (q2Extended=q2Extended, MCond=MCond))
}


## Calculate 2nd and 3rd (non-central) moments (mu2 and mu3) and
## skewness of given transition matrix and associated fecundity
## matrix.  Inputs: reward matrices for first, second, third moments.
## These calculations come from van Daalen, S. F., and
## H. Caswell. 2017. Lifetime reproductive output: individual
## stochasticity, variance, and sensitivity analysis. Theoretical
## Ecology 10:355 – 374.

calcMoments = function (M, R1, R2, R3) {

  numStates = dim(M)[1]

  ## set up Mplus, the transition matrix with absorbing state (dead)
  Mplus = matrix (0, numStates+1, numStates+1)
  Mplus[1:numStates, 1:numStates] = M
  Mplus[numStates+1, 1:numStates] = 1 - colSums(M)
  Mplus[numStates+1, numStates+1] = 1

  ## Set up Z, which cleaves off dead states
  Z = cbind (diag(numStates), rep(0, numStates))

  ## Set up the vector 1_s, where s = bigmz + 1
  oneS = rep(0, numStates+1)
  oneS[numStates+1] = 1

  ## R1Stripped has the absorbed state cleaved off
  R1Stripped = R1[1:numStates, 1:numStates]
  R2Stripped = R2[1:numStates, 1:numStates]

  ## Get zero-centered moments rho1, rho2, rho3

  ## Ex(R)
  e = rep (1, numStates+1)
  N = solve (diag(numStates) - M)
  rho1Vec = t(N) %*% Z %*% t(Mplus * R1) %*% e

  ## Ex(R^2 | x)
  rho2Vec = t(N) %*% (Z %*% t(Mplus * R2) %*% e +
                        2*t(M * R1Stripped) %*% rho1Vec)

  ## Ex(R^3 | x)
  rho3Vec = t(N) %*% (Z %*% t(Mplus * R3) %*% e +
                    3*t(M*R2Stripped) %*% rho1Vec +
                    3*t(M*R1Stripped) %*% rho2Vec)

  ## non-central moments
  rho1Vec = t(rho1Vec)
  rho2Vec = t(rho2Vec)
  rho3Vec = t(rho3Vec)

  ## central moments
  mu2Vec = rho2Vec - rho1Vec^2
  mu3Vec = rho3Vec - 3*rho1Vec*rho2Vec + 2*rho1Vec^3

  ## skewness. NOTE, skewness=0 whenever the variance=0. 
  skewnessVec = rep(0,length(mu3Vec)); 
  e = which(mu2Vec>0); 
  skewnessVec[e] = mu3Vec[e]/mu2Vec[e]^1.5

  return (out=list(rho1Vec=rho1Vec, mu2Vec=mu2Vec, mu3Vec=mu3Vec,
                   skewnessVec=skewnessVec))
}

########################################################################
#### Code graveyard.  Useful but ultimately unused #####################
########################################################################

## Calculate probability of not being successful conditional on
## initial state and kernel conditional on not becoming successful
## before death
makeCondFailureKernel = function (M, transientStates) {

  ## How many states are there all told?
  numStates = dim(M)[1]
  ## Create U, the transition matrix for the transient states
  U = M[transientStates, transientStates]
  ## How many transient states are there?
  numTransient = length(transientStates)
  if (FALSE) {
    ## calculate a1, the probability of dying in the next time step
    a1 = (1 - apply (M, 2, sum))[transientStates]
    ## calculate q1, the probability of dying before hitting the
    ## "successful" absorbing state given that you are now in state i.
    NU = solve (diag(numTransient) - U)
    q1 = a1 %*% NU
  }
  ## calculate a2, the probability of hitting an absorbing state in
  ## the next time step
  a2 = apply (M, 2, sum)[transientStates] - apply (U, 2, sum)
  ## calculate q2, the probability of being in Z2 before you die given
  ## that you are now in state i.
  NU = solve (diag(numTransient) - U)
  q2 = a2 %*% NU
  q1 = 1 - q2
  
  ## Transition matrix conditional on not reproducing before death.
  MCond = matrix (0, numTransient, numTransient)
  for (j in 1:numTransient)
    MCond[,j] = M[transientStates,transientStates[j]] * q1 / q1[j]

  return (out=list (q1=q1, MCond=MCond))
}
```
