## Supplementary material for "To prosper, live long: understanding the sources of reproductive skew and extreme reproductive success in structured populations": Associated R code: plotSculpinLifespanCondLRO3.R

```
#########################################################################
##
## sculpinLifespanCondR.Rdata and
## sculpinLifespanCondRNoEnvVarInFec.Rdata generated by
## sculpinLifespanCondLRO.R
##
#########################################################################

rm(list=ls(all=TRUE)); graphics.off();
load ("SculpinFalsePoint.Rdata")

noEnvVarInFecundity = FALSE

if (noEnvVarInFecundity == TRUE) {
  load ("sculpinLifespanCondRNoEnvVarInFec.Rdata")
} else {
  load ("sculpinLifespanCondR.Rdata")
}

##maxKids = 15
##mT = maxKids + 1
##maxAge = 50
##percentileCutoff = 0.96

##maxKidsIndex = R99
##maxKidsIndex = dim(probLifespanCondR)[1]
##LCutoff = dim(probLifespanCondR)[2]
RCutoff = R99
maxKidsIndex = RCutoff + 1
LCutoff = L99

centralPanel = matrix(1, 7, 6)
horizPanel = matrix (2, 1, 6)
mat = rbind(centralPanel, horizPanel)
vertPanel = matrix(c(rep(3, 7), 0), 8, 1)
mat = cbind (vertPanel, mat)
layout (mat=mat)
##par(mgp=c(2,1,0), cex.axis=1.3, cex.lab=1.4);
par(cex.axis=1.3, cex.lab=1.4);
##par (mar=c(2,6,3,1))
par (mar=c(2,6,2,1))

image (0:RCutoff, 1:LCutoff,
       probLifespanCondR[1:maxKidsIndex,1:LCutoff], axes=TRUE,
       xlab="", ylab="", ylim=c(1, LCutoff), xlim=c(0, maxKidsIndex-1) )
contour (0:RCutoff, 1:LCutoff,
         probLifespanCondR[1:maxKidsIndex,1:LCutoff], add=T)

##axis (side=1, 0:maxKidsIndex)
##axis (side=2, 0:LCutoff)
##axis (side=2, at=c(L50, L75, L90, L95),
##      label=c("50th %ile", "75th %ile", "90th %ile", "95th %ile"))
##mtext ("Lifespan", side=2, line=3.5, cex=1.3)

## add a skinny probability density for P(R) below the plot
par (mar=c(2,5,0,0))
plot (0:RCutoff, distKidsAtDeath[1:maxKidsIndex], axes=FALSE,
      type="l", lwd=2,
      xlab="", ylab="")
axis (side=1, at=c(R50, R90, R95),
      label=c("50th %ile", "90th %ile", "95th %ile"))
mtext ("Lifetime reproductive output", side=3, line=-2, cex=1.3)

## add a skinny probability density for P(L) to the left of the plot
par (mar=c(0, 0, 0, 2))
##par (mar=c(2,0,3,2))
plot (-probLifespan[1:LCutoff], 1:LCutoff, axes=FALSE, type="l", lwd=2,
      xlab="", ylab="", ylim=c(1, LCutoff))
axis (side=4, at=c(L50, L90, L95),
      label=c("50th %ile", "90th %ile", "95th %ile"))
mtext ("Lifespan", side=2, line=-11, cex=1.3)

if (noEnvVarInFecundity == TRUE) {
##  dev.copy2pdf (file="sculpinFalsePointLifespanCondLRONoEnvVarInFec.pdf")
} else {
##  dev.copy2pdf (file="sculpinFalsePointLifespanCondLRO.pdf")
}
```
