## Supplementary material for "To prosper, live long: understanding the sources of reproductive skew and extreme reproductive success in structured populations": Associated R code: plotUmboniumLifespanCondLRO2.R

```
#############################################################################
##
## umboniumLifespanCondLRO.Rdata is generated by umboniumLifespanCondLRO.R
##
#############################################################################

rm(list=ls(all=TRUE)); graphics.off();
load ("umboniumLifespanCondLRO.Rdata")
load ("Umbonium.Rdata")

##maxKids = 15
##mT = maxKids + 1
##maxAge = 50
percentileCutoff = 0.99

maxKidsIndex = R99
LCutoff = L99

centralPanel = matrix(1, 7, 6)
horizPanel = matrix (2, 1, 6)
mat = rbind(centralPanel, horizPanel)
vertPanel = matrix(c(rep(3, 7), 0), 8, 1)
mat = cbind (vertPanel, mat)
layout (mat=mat)
par(mgp=c(2,1,0), cex.axis=1.3, cex.lab=1.4);
par (mar=c(2,6,3,1))

image (0:(maxKidsIndex-1), 1:LCutoff,
       probLifespanCondR[1:maxKidsIndex,1:LCutoff], axes=TRUE,
       xlab="", ylab="", xlim=c(0, maxKidsIndex-1), ylim=c(1,LCutoff))
contour (0:(maxKidsIndex-1), 1:LCutoff,
         probLifespanCondR[1:maxKidsIndex,1:LCutoff], add=T)

par (mar=c(2,5,0,1))  
plot (0:(maxKidsIndex-1), distKidsAtDeath[1:maxKidsIndex], axes=FALSE, type="l", lwd=2,
      xlab="", ylab="")
axis (side=1, at=c(R50, R90, R95),
      label=c("50th %ile", "90th %ile", "95th %ile"))
mtext ("Lifetime reproductive output", side=3, line=-2, cex=1.3)

## add a skinny probability density for P(L) to the left of the plot
par (mar=c(0, 0, 0, 2))
plot (-probLifespan[1:LCutoff], 1:LCutoff, axes=FALSE, type="l", lwd=2,
      xlab="", ylab="")
axis (side=4, at=c(L50, L90, L95),
      label=c("50th %ile", "90th %ile", "95th %ile"))
mtext ("Lifespan", side=2, line=-11, cex=1.3)

##dev.copy2pdf (file="umboniumLifespanCondLRO.pdf")
```
