## Supplementary material for "To prosper, live long: understanding the sources of reproductive skew and extreme reproductive success in structured populations": Associated R code: README_r1.txt

Title: To prosper, live long: understanding the sources of
reproductive skew and extreme reproductive success in structured
populations

Authors: Robin E. Snyder (, Case Western Reserve
University) and Stephen P. Ellner (, Cornell
University)

Robin Snyder is largely responsible for the code.

We used R version 4.0.2 running on the Ubuntu linux operating system,
kernel version 5.15.0.

Data abstract: The paper explains how to partition reproductive skew
--- i.e. the skewness of lifetime reproductive output (LRO) --- into
contributions from fixed trait variation; four forms of ``demographic
luck'' (birth state, fecundity luck, survival trajectory luck, and
growth trajectory luck); and two kinds of ``environmental luck''
(birth environment, and environment trajectory). Each of these is
further partitioned into contributions at different ages.  The paper
also presents a method to calculate the distribution of lifespan
conditional on LRO value, as a means of exploring the degree to which
exceptional reproductive success is the result of an exceptionally
long life.  The code in this collection implements those calculations
and generates the figures in the paper, which are a collection of
results for case studies.

The code to produce each figure in the paper is listed in the figure
captions.  Here we repeat that information in somewhat different form.

### Utility code collections ############################################

Utilities.R: some routines for calculating the dominant
eigenvalue/vector and for visualizing matrices.

megamatrixFunctions.R: various functions for dealing with megamatrices
(i.e. transition matrices for situations where state is classified by
multiple variables, e.g. size and environment or size and environment
and number of offspring to date)

partitioningUtilities.R: Some utilities for partitioning, e.g. making
a kernel conditional on reaching a set of absorbing states.

genericPartitionVarSkewness6.R: contains the routines that do the bulk
of the calculations.  Called by all of the case study
figure-generating scripts.  Calls functions in the previous utility scripts.

### Scripts associated with particular figures #########################

********************************************************************
All the code in this section calculates the contributions to the
variance, third central moment (mu_3), or skewness of LRO, for various
case studies.
********************************************************************

snowgooseVarSkewnessPartition3.R: generates fig. 2 variance and
skewness subplots

snowgooseVarMu3Partition.R: generates fig. 2 mu_3 subplot

sculpinVarSkewnessPartition2.R: generates fig. 3 variance and
skewness subplots

sculpinVarMu3Partition.R: generates fig. 3 mu_3 subplot

fulmarVarSkewnessPartition.R: generates fig. 4 variance and
skewness subplots

fulmarVarMu3Partition.R: generates fig. 4 mu_3 subplot

kittiwakeVarSkewnessPartition.R: generates fig. 5 variance and
skewness subplots

kittiwakeVarMu3Partition.R: generates fig. 5 mu_3 subplot

--- SI figures: ----------

kestrelVarSkewnessPartition.R: generates fig. S2 variance and
skewness subplots

kestrelVarMu3Partition.R: generates fig. S2 mu_3 subplot

umboniumVarSkewnessPartition.R: generates fig. S3 variance and
skewness subplots

umboniumVarMu3Partition.R: generates fig. S3 mu_3 subplot

lomatiumkewnessPartition5.R: generates fig. S4 variance and
skewness subplots

lomatiumVarMu3Partition.R: generates fig. S4 mu_3 subplot

xenosaurusVarSkewnessPartition.R: generates fig. S5 variance and
skewness subplots

xenosaurusVarMu3Partition.R: generates fig. S5 mu_3 subplot


*************************************************************
The code in this section calculates the distribution of lifespan
conditional on LRO value, for different case studies.
*************************************************************

Fig. 6 scripts:

First run snowgooseLifespanCondLRO.R, umboniumLifespanCondLRO.R, and
sculpinLifespanCondLRO.R, which do the actual calculations and store
the results in snowgooseLifespanCondLRO.Rdata,
umboniumLifespanCondLRO.Rdata, and sculpinLifespanCondLRO.Rdata,
respectively.  These contain probLifespanCondR and distKidsAtDeath,
where probLifespanCondR[i,j] is the probability of having lifespan j-1
given that you had LRO equal to i-1 and distKidsAtDeath[j] is the
probability of having LRO equal to j-1.

Then run plotSnowgooseLifespanCondLRO3.R,
plotUmboniumLifespanCondLRO2.R, and plotSculpinLifespanCondLRO3.R to
create the subplots in fig. 6.

**********************************************************************
The code in this section calculates the 90% central interval for the
probability of lifespan conditional on LRO value, for different case
studies.
**********************************************************************

Fig 7 scripts:

First run umboniumLifespanCondLRO.R once with noEnvVarInFecundity set
to TRUE and once with noEnvVarInFecundity set to FALSE in order to
generate umboniumCVLifespanCondRNoEnvVarInFec.dat and
umboniumCVLifespanCondR.dat, respectively.  These contain
probLifespanCondR, probLifespan, and distKidsAtDeath, where
probLifespanCondR[i,j] is the probability of having lifespan j-1 given
that you had LRO equal to i-1, probLifespan[j] is the probability of
having lifespan j-1, and distKidsAtDeath[j] is the probability of
having LRO equal to j-1.

Then run umboniumLifespanCIVsLROPercentile.R, which uses these .dat
files to create umboniumCILifespan.dat and
umboniumCILifespanNoEnvVarInFec.dat.  Then run
compareUmboniumLifespanCIVsLROPercentile.R, which generates fig. 7.

----------------------------

Fig 8 scripts:

First run kestrelLifespanCondLRO.R once with highSurvival set to TRUE
and once with highSurvival set to FALSE in order to generate
kestrelLifespanCondLROHighSurv.Rdata and kestrelLifespanCondLRO.Rdata,
respectively.  These contain probLifespanCondR, probLifespan, and
distKidsAtDeath, where probLifespanCondR[i,j] is the probability of
having lifespan j-1 given that you had LRO equal to i-1,
probLifespan[j] is the probability of having lifespan j-1, and
distKidsAtDeath[j] is the probability of having LRO equal to j-1.

Then run kestrelLifespanCIVsLROPercentile.R, which uses
these .Rdata files to create kestrelCILifespan.dat and
kestrelCILifespanHighSurv.dat.  Then run
compareKestrelLifespanCIVsLROPercentile.R, which generates fig. 8.


### Model matrix values: #################################################

fulmarMatrices directory: matrix values for each combination of
behavioral group and sea ice conditions.  Farray[i,j,k,m] is the
expected number of stage k offspring produced by a stage m individual
in behavioral syndrome group i under sea ice condition j.
Parray[i,j,k,m] is the probability that a stage m individual in
behavioral syndrome group i will survive and transition to become a
stage k individual under sea ice condition j.  Behavioral group
membership does not change over the course of a life and all offspring
inherit the parent's behavioral type (not that this matters for the
questions being investigated here).

kittiwake/KittiwakeMatrices.R: kittiwake matrix values.  M1 is the
survival/growth transition matrix for age 1 individuals, M2 is the
same matrix for age 2 individuals, and so forth up through M5, which
is for those age 5 and older.  E.g. M1[i,j] is the probability that a
one-year-old stage j individual survives and becomes a two-year-old
stage i individual.

Lomatium directory: Lomatium matrix values for the Rose Prairie #1
data set and code to create the survival/growth and fecundity
matrices.  makeP.R makes Flist and Plist.  Flist[[k]][i,j] is the
expected number of offspring of stage i produced by an individual of
stage j in year k.  Plist[[k]][i,j] is the probability that a
stage j individual will survive and transition to become a stage i
individual in year k.

SnowGoose.Rdata, SculpinFalsePoint.Rdata, Umbonium.Rdata,
Falco_naumanni.Rdata, TropicalXenosaurus.Rdata: fecundity and
survival/growth transition matrices in multiple years/environment
types extracted from COMADRE for snow geese, sculpins, Umbonium,
lesser kestrels, and Xenosaurus.  In each case, Flist is a list of
fecundity matrices: Flist[[k]][i,j] is the expected number of
offspring of stage i produced by an individual of stage j in year or
environment k.  Ulist is a list of survival/growth transition
matrices: Ulist[[k]][i,j] is the probability that a stage j individual
will survive and transition to become a stage i individual in year or
environment k.
