## Supplementary material for "To prosper, live long: understanding the sources of reproductive skew and extreme reproductive success in structured populations": Associated R code: sculpinLifespanCondLRO.R

```
####################################################################
##
## Generates .Rdata files for plotSculpinLifespanCondLRO3.R
##
####################################################################

rm(list=ls(all=TRUE)); graphics.off();
source ("partitioningUtilities.R")
source ("genericPartitionVarSkewness6.R")
source ("megamatrixFunctions.R")

load ("SculpinFalsePoint.Rdata")
##load ("SculpinOceanBeach.Rdata")

maxAge = 9
##maxKids = 27
maxKids = 30

## Set noEnvVarInFecundity if you want fecundity in each stage to be
## the average of its values across environments.
noEnvVarInFecundity = TRUE
if (noEnvVarInFecundity)
  cat ("No env. variation in fecundity\n")

## Assume all year types are equally likely
numEnv = length(Ulist)
Q = matrix (1/numEnv, numEnv, numEnv)
u0 = rep (1/numEnv, numEnv)

## Assume all individuals are born into stage 1
mz = dim(Ulist[[1]])[1]
c0 = rep (0, mz)
c0[1] = 1

if (noEnvVarInFecundity) {
  Fave = matrix (0, mz, mz)
  for (q in 1:numEnv)
    Fave = Fave + Flist[[q]]
  Fave = Fave/numEnv
  for (q in 1:numEnv)
    Flist[[q]] = Fave
}

## Set up cross-classified birth distribution
bigmz = numEnv*mz
m0 = matrix (outer (c0, as.vector(u0)), bigmz, 1)

out = distLifespanCondR2 (Ulist, Flist, Q, m0, maxKids,
                          maxAge, percentileCutoff=0.99)


distKidsAtDeath = out$distKidsAtDeath
probLifespanCondR = out$probLifespanCondR
sdLifespanCondR = out$sdLifespanCondR
CVLifespanCondR = out$CVLifespanCondR
maxKidsIndex = out$maxKidsIndex
normalSurvProb = out$normalSurvProb

## Subtract 1 to change numKids index into actual number of kids
R50 = min(which(cumsum(distKidsAtDeath) >= 0.5)) - 1
R90 = min(which(cumsum(distKidsAtDeath) >= 0.9)) - 1
R95 = min(which(cumsum(distKidsAtDeath) >= 0.95)) - 1
R99 = min(which(cumsum(distKidsAtDeath) >= 0.99)) - 1

probLifespan = -diff(normalSurvProb)

## Subtract 1 to change age index into age
L50 = min(which(cumsum(probLifespan) >= 0.5)) - 1
L75 = min(which(cumsum(probLifespan) >= 0.75)) - 1
L90 = min(which(cumsum(probLifespan) >= 0.9)) - 1
L95 = min(which(cumsum(probLifespan) >= 0.95)) - 1
L99 = min(which(cumsum(probLifespan) >= 0.99)) - 1

## At what age should we cut off the plots?
##LCutoff = min(which(probLifespanCondR[1,] < 0.001))
LCutoff = L99

if (noEnvVarInFecundity) {
  save (probLifespan, distKidsAtDeath, probLifespanCondR,
        CVLifespanCondR, sdLifespanCondR,
        L50, L75, L90, L95, L99, R50, R90, R95, R99,
        file="sculpinLifespanCondRNoEnvVarInFec.Rdata")
} else {
  save (probLifespan, distKidsAtDeath, probLifespanCondR,
        CVLifespanCondR, sdLifespanCondR,
        L50, L75, L90, L95, L99, R50, R90, R95, R99,
        file="sculpinLifespanCondR.Rdata")
}
```
