## Supplementary material for "To prosper, live long: understanding the sources of reproductive skew and extreme reproductive success in structured populations": Associated R code: sculpinVarMu3Partition.R

```
rm(list=ls(all=TRUE)); graphics.off();
source ("partitioningUtilities.R")
source ("genericPartitionVarSkewness6.R")

load ("SculpinFalsePoint.Rdata")

lineTypes = c(1,2,3,4)
names(lineTypes) = c("surv", "env", "growth", "fec")

colors = c("red", "green", "orange", "blue")
names(colors) = c("surv", "env", "growth", "fec")

## Assume all year types are equally likely
numEnv = length(Ulist)
Q = matrix (1/numEnv, numEnv, numEnv)
u0 = rep (1/numEnv, numEnv)

## Assume all individuals are born into stage 1
mz = dim(Ulist[[1]])[1]
c0 = rep (0, mz)
c0[1] = 1

## Set up cross-classified birth distribution
bigmz = numEnv*mz
m0 = matrix (outer (c0, as.vector(u0)), bigmz, 1)

out = getVarMu3PartitionsEnvVar (Plist=Ulist, Flist, Q, m0)

birthEnvMu3 = out$birthEnvMu3
survTrajecMu3 = out$survTrajecMu3
growthTrajecMu3 = out$growthTrajecMu3
envTrajecMu3 = out$envTrajecMu3
fecMu3 = out$fecMu3
totMu3 = out$totMu3
lifespan = out$lifespan

totSurvTrajecMu3 = sum(survTrajecMu3)
totGrowthTrajecMu3 = sum(growthTrajecMu3)
totEnvTrajecMu3 = sum(envTrajecMu3)
totFecMu3 = sum(fecMu3)

out = getModalTimeToFirstRepro (Plist=Ulist, Flist, Q, m0)
reproArrivalMode = out$reproArrivalMode

birthEnvVar = out$birthEnvVar
survTrajecVar = out$survTrajecVar
growthTrajecVar = out$growthTrajecVar
envTrajecVar = out$envTrajecVar
fecVar = out$fecVar
totVar = out$totVar

totSurvTrajecVar = sum(survTrajecVar)
totGrowthTrajecVar = sum(growthTrajecVar)
totEnvTrajecVar = sum(envTrajecVar)
totFecVar = sum(fecVar)

## Plot skewness

par (cex.lab=1.4, lwd=2)

ymax = max(survTrajecMu3, growthTrajecMu3,
           envTrajecMu3)
ymin = min(survTrajecMu3, growthTrajecMu3, envTrajecMu3)
plot (0:10, survTrajecMu3[1:11], col=colors["surv"],
      lty=lineTypes["surv"],
      xlab="Age", ylab="Contributions to Mu3(R)", type="l",
      ylim=c(ymin, ymax))
lines (0:10, growthTrajecMu3[1:11], col=colors["growth"],
      lty=lineTypes["growth"])
lines (0:10, envTrajecMu3[1:11], col=colors["env"],
      lty=lineTypes["env"])
lines (0:10, fecMu3[1:11], col=colors["fec"],
      lty=lineTypes["fec"])
lines (rep(lifespan, 2), c(0, 0.35*ymax), col="black",
       lty=1, lwd=2)
lines (rep(reproArrivalMode, 2), c(0, ymax), col="purple",
       lty=1, lwd=2)

legend (x = 4, y=0.95*ymax, legend=c("99% of individ. are dead",
                                     "Mode(age of 1st repro.)"),
        col=c("black", "purple"),
        lty=1, bty="n", lwd=2, cex=1.2)

legend (x=4, y=0.8*ymax,
        legend=c(paste("Surv. trajec. luck: ",
                       round(100*totSurvTrajecMu3/totMu3, digits=1),
                       "%", sep=""),
                 paste("Env. trajec. luck: ",
                       round(100*totEnvTrajecMu3/totMu3, digits=1),
                       "%", sep=""),
                 paste("Growth trajec. luck: ",
                       round(100*totGrowthTrajecMu3/totMu3, digits=1),
                       "%", sep=""),
                 paste("Fecundity luck: ",
                       round(100*totFecMu3/totMu3, digits=1),
                       "%", sep="")),
        col=colors, lty=lineTypes, bty="n", lwd=2, cex=1.2)

xloc = 0.6*10

if (FALSE) {
  text (x=xloc, y=0.56*ymax,
        paste("Surv trajec.: ",
              round(100*totSurvTrajecMu3/totMu3, digits=1),
              "%", sep=""), col="black")

  text (x=xloc, y=0.5*ymax,
        paste("Env. trajec.: ",
              round(100*totEnvTrajecMu3/totMu3, digits=1),
              "%", sep=""), col="green")

  text (x=xloc, y=0.44*ymax,
        paste("Growth trajec.: ",
              round(100*totGrowthTrajecMu3/totMu3, digits=1),
              "%", sep=""), col="orange")

  text (x=xloc, y=0.32*ymax,
        paste("Fecundity: ",
              round(100*totFecMu3/totMu3, digits=1),
              "%", sep=""), col="blue")
}

text (x=xloc, y=0.48*ymax, cex=1.2,
        paste("Birth env. luck: ", 
              round(100*birthEnvMu3/totMu3, digits=1),
              "%", sep=""), col="black")

dev.copy2pdf (file="sculpinMu3Partition.pdf")

if (FALSE) {
## And plot variance
dev.new ()
par (cex.lab=1.4, lwd=2)
ymax = max(survTrajecVar, growthTrajecVar,
           envTrajecVar)
ymin = min(survTrajecVar, growthTrajecVar, envTrajecVar)
plot (0:10, survTrajecVar[1:11], col="black",
      xlab="Age", ylab="Var contributions", type="l",
      ylim=c(ymin, ymax))
lines (0:10, growthTrajecVar[1:11], col="orange")
lines (0:10, envTrajecVar[1:11], col="green")
lines (0:10, fecVar[1:11], col="blue")
lines (rep(lifespan, 2), c(0, 0.35*ymax), col="black",
       lty=2, lwd=2)
if (FALSE) {
  legend (x="topright", legend=c("Surv. trajec. variance",
                                 "Growth trajec. variance",
                                 "Env. trajec. variance",
                                 "Fec. variance",
                                 "99% of individ. are dead"),
          col=c("black", "orange", "green", "blue", "black"),
          lty=c(1,1,1,1,1,2), bty="n", lwd=2)
} else {
  legend (x = 4, y=5.5, legend="99% of individ. are dead",
          lty=2, bty="n", lwd=2)
}

xloc = 0.6*10

text (x=xloc, y=0.66*ymax,
      paste("Surv trajec.: ",
            round(100*totSurvTrajecVar/totVar, digits=1),
            "%", sep=""), col="black")

text (x=xloc, y=0.6*ymax,
      paste("Env. trajec.: ",
            round(100*totEnvTrajecVar/totVar, digits=1),
            "%", sep=""), col="green")

text (x=xloc, y=0.54*ymax,
      paste("Growth trajec.: ",
            round(100*totGrowthTrajecVar/totVar, digits=1),
            "%", sep=""), col="orange")

text (x=xloc, y=0.48*ymax,
      paste("Fecundity: ",
            round(100*totFecVar/totVar, digits=1),
            "%", sep=""), col="blue")

text (x=xloc, y=0.42*ymax,
      paste("Birth env.: ",
            round(100*birthEnvVar/totVar, digits=1),
            "%", sep=""), col="black")


##dev.copy2pdf (file="sculpinVarPartition.pdf")


}
```
