## Supplementary material for "To prosper, live long: understanding the sources of reproductive skew and extreme reproductive success in structured populations": Associated R code: snowgooseVarSkewnessPartition3.R

```
## includes modal age of 1st successful repro.

rm(list=ls(all=TRUE)); graphics.off();

source ("partitioningUtilities.R")
source ("genericPartitionVarSkewness6.R")

load ("SnowGoose.Rdata")

lineTypes = c(1,2,3,4)
names(lineTypes) = c("surv", "env", "growth", "fec")

colors = c("red", "green", "orange", "blue")
names(colors) = c("surv", "env", "growth", "fec")

## Assume all year types are equally likely
numEnv = length(Ulist)
Q = matrix (1/numEnv, numEnv, numEnv)
u0 = rep (1/numEnv, numEnv)

## Assume all individuals are born into stage 1
mz = dim(Ulist[[1]])[1]
c0 = rep (0, mz)
c0[1] = 1

## Set up cross-classified birth distribution
bigmz = numEnv*mz
m0 = matrix (outer (c0, as.vector(u0)), bigmz, 1)

out = getVarSkewnessPartitionsEnvVar (Plist=Ulist, Flist, Q, m0)

birthEnvSkewness = out$birthEnvSkewness
survTrajecSkewness = out$survTrajecSkewness
growthTrajecSkewness = out$growthTrajecSkewness
envTrajecSkewness = out$envTrajecSkewness
fecSkewness = out$fecSkewness
totSkewness = out$totSkewness
lifespan = out$lifespan

totSurvTrajecSkewness = sum(survTrajecSkewness)
totGrowthTrajecSkewness = sum(growthTrajecSkewness)
totEnvTrajecSkewness = sum(envTrajecSkewness)
totFecSkewness = sum(fecSkewness)

birthEnvVar = out$birthEnvVar
survTrajecVar = out$survTrajecVar
growthTrajecVar = out$growthTrajecVar
envTrajecVar = out$envTrajecVar
fecVar = out$fecVar
totVar = out$totVar

totSurvTrajecVar = sum(survTrajecVar)
totGrowthTrajecVar = sum(growthTrajecVar)
totEnvTrajecVar = sum(envTrajecVar)
totFecVar = sum(fecVar)

out = getModalTimeToFirstRepro (Plist=Ulist, Flist, Q, m0)
reproArrivalMode = out$reproArrivalMode

## Plot skewness

par (cex.lab=1.4, lwd=2)

ymax = max(survTrajecSkewness, growthTrajecSkewness,
           envTrajecSkewness, fecSkewness)
ymin = min(survTrajecSkewness, growthTrajecSkewness,
           envTrajecSkewness, fecSkewness)
plot (0:30, survTrajecSkewness[1:31], col=colors["surv"],
      lty=lineTypes["surv"],
      xlab="Age", ylab="Contributions to Skewness(R)", type="l",
      ylim=c(ymin, ymax))
lines (0:30, growthTrajecSkewness[1:31], col=colors["growth"],
      lty=lineTypes["growth"])
lines (0:30, envTrajecSkewness[1:31], col=colors["env"],
      lty=lineTypes["env"])
lines (0:30, fecSkewness[1:31], col=colors["fec"],
      lty=lineTypes["fec"])
lines (rep(lifespan, 2), c(0, 0.4*ymax), col="black",
       lty=1, lwd=2)
lines (rep(reproArrivalMode, 2), c(0, ymax), col="purple",
       lty=1, lwd=2)
if (FALSE) {
  legend (x="topright", legend=c("Surv. trajec. skewness",
                                 "Growth trajec. skewness",
                                 "Env. trajec. skewness",
                                 "Fec. skewness",
                                 "99% of individ. are dead"),
          col=c("red", "orange", "green", "blue", "black"),
          lty=c(1,1,1,1,1,2), bty="n", lwd=2)
} else {
  legend (x = 11, y=0.94*ymax, legend=c("99% of individ. are dead",
                                   "Mode(age of 1st repro.)"),
          col=c("black", "purple"),
          lty=1, bty="n", lwd=2, cex=1.2)

  legend (x=11, y=0.77*ymax,
          legend=c(paste("Surv trajec. luck: ",
                         round(100*totSurvTrajecSkewness/totSkewness, digits=1),
                         "%", sep=""),
                   paste("Env. trajec. luck: ",
                         round(100*totEnvTrajecSkewness/totSkewness, digits=1),
                         "%", sep=""),
                   paste("Growth trajec. luck: ",
                         round(100*totGrowthTrajecSkewness/totSkewness, digits=1),
                         "%", sep=""),
                   paste("Fecundity luck: ",
                         round(100*totFecSkewness/totSkewness, digits=1),
                         "%", sep="")),
          col=colors, lty=lineTypes, bty="n", lwd=2, cex=1.2)
}

xloc = 0.6*30

if (FALSE) {
  text (x=xloc, y=0.66*ymax, cex=1.2,
        paste("Surv. trajec. luck: ",
              round(100*totSurvTrajecSkewness/totSkewness, digits=1),
              "%", sep=""), col=colors["surv"])

  text (x=xloc, y=0.6*ymax, cex=1.2,
        paste("Env. trajec. luck: ",
              round(100*totEnvTrajecSkewness/totSkewness, digits=1),
              "%", sep=""), col=colors["env"])

  text (x=xloc, y=0.54*ymax, cex=1.2,
        paste("Growth trajec. luck: ",
              round(100*totGrowthTrajecSkewness/totSkewness, digits=1),
              "%", sep=""), col=colors["growth"])

  text (x=xloc, y=0.48*ymax, cex=1.2,
        paste("Fecundity luck: ",
              round(100*totFecSkewness/totSkewness, digits=1),
              "%", sep=""), col=colors["fec"])
}

text (x=xloc, y=0.42*ymax, cex=1.2,
      paste("Birth env. luck: ",
            round(100*birthEnvSkewness/totSkewness, digits=1),
            "%", sep=""), col="black")


dev.copy2pdf (file="snowgooseSkewnessPartition.pdf")

## And plot variance
dev.new ()
par (cex.lab=1.4, lwd=2)
ymax = max(survTrajecVar, growthTrajecVar,
           envTrajecVar)
ymin = min(survTrajecVar, growthTrajecVar, envTrajecVar)
plot (0:30, survTrajecVar[1:31], col=colors["surv"],
      lty=lineTypes["surv"],
      xlab="Age", ylab="Contributions to Var(R)", type="l",
      ylim=c(ymin, ymax))
lines (0:30, growthTrajecVar[1:31], col=colors["growth"],
      lty=lineTypes["growth"])
lines (0:30, envTrajecVar[1:31], col=colors["env"],
      lty=lineTypes["env"])
lines (0:30, fecVar[1:31], col=colors["fec"],
      lty=lineTypes["fec"])
lines (rep(lifespan, 2), c(0, 0.4*ymax), col="black",
       lty=1, lwd=2)
lines (rep(reproArrivalMode, 2), c(0, ymax), col="purple",
       lty=1, lwd=2)


if (FALSE) {
  legend (x="topright", legend=c("Surv. trajec. variance",
                                 "Growth trajec. variance",
                                 "Env. trajec. variance",
                                 "Fec. variance",
                                 "99% of individ. are dead"),
          col=c("red", "orange", "green", "blue", "black"),
          lty=c(1,1,1,1,1,2), bty="n", lwd=2)
} else {
  legend (x = 11, y=0.94*ymax, legend=c("99% of individ. are dead",
                                        "Mode(age of 1st repro.)"),
          col=c("black", "purple"),
          lty=1, bty="n", lwd=2, cex=1.2)

  legend (x=11, y=0.77*ymax,
          legend=c(paste("Surv. trajec. luck: ",
                         round(100*totSurvTrajecVar/totVar, digits=1),
                         "%", sep=""),
                   paste("Env. trajec. luck: ",
                         round(100*totEnvTrajecVar/totVar, digits=1),
                         "%", sep=""),
                   paste("Growth trajec. luck: ",
                         round(100*totGrowthTrajecVar/totVar, digits=1),
                         "%", sep=""),
                   paste("Fecundity luck: ",
                         round(100*totFecVar/totVar, digits=1),
                         "%", sep="")),
          col=colors, lty=lineTypes, bty="n", lwd=2, cex=1.2)
}

xloc = 0.6*30

if (FALSE) {
  text (x=xloc, y=0.66*ymax, cex=1.2,
        paste("Surv trajec.: ",
              round(100*totSurvTrajecVar/totVar, digits=1),
              "%", sep=""), col=colors["surv"])

  text (x=xloc, y=0.6*ymax, cex=1.2,
        paste("Env. trajec.: ",
              round(100*totEnvTrajecVar/totVar, digits=1),
              "%", sep=""), col=colors["env"])

  text (x=xloc, y=0.54*ymax, cex=1.2,
        paste("Growth trajec.: ",
              round(100*totGrowthTrajecVar/totVar, digits=1),
              "%", sep=""), col=colors["growth"])

  text (x=xloc, y=0.48*ymax, cex=1.2,
        paste("Fecundity: ",
              round(100*totFecVar/totVar, digits=1),
              "%", sep=""), col=colors["fec"])
}

text (x=xloc, y=0.44*ymax, cex=1.2,
      paste("Birth env. luck: ",
            round(100*birthEnvVar/totVar, digits=1),
            "%", sep=""), col="black")


dev.copy2pdf (file="snowgooseVarPartition.pdf")
```
