## Supplementary material for "To prosper, live long: understanding the sources of reproductive skew and extreme reproductive success in structured populations": Associated R code: umboniumLifespanCIVsLROPercentile.R

```
#########################################################################
##
## kestrelLifespanCIVsLROPercentile.R: Calculates the 90% confidence interval
## of lifespan conditional on LRO.  Both LRO and lifespan are
## expressed as percentiles so that we can compare across species.
##
## umboniumLifespanCondLRONoEnvVarInFec.Rdata and
## umboniumLifespanCondLRO.Rdata are generated by
## umboniumLifespanCondLRO.R. 
##
########################################################################

rm(list=ls(all=TRUE)); graphics.off();

noEnvVarInFecundity = FALSE

if (noEnvVarInFecundity) {
  load ("umboniumLifespanCondLRONoEnvVarInFec.Rdata")
} else {
  load ("umboniumLifespanCondLRO.Rdata")
}


cumDistAgeAtDeath = cumsum (probLifespan)

RpercentileVector = seq (0.5, 0.99, 0.01)
numPercentiles = length (RpercentileVector)
RCutoffs = bottom5Lifespan = bottom5LifespanPercentile =
  top5Lifespan = top5LifespanPercentile = mlePercentile =
    numeric (numPercentiles)

maxKidsIndex = dim(probLifespanCondR)[1]
LCutoff = dim(probLifespanCondR)[2]

for (ii in 1:numPercentiles) {
  RCutoffs[ii] = min(which(cumsum(distKidsAtDeath) >=
                           RpercentileVector[ii]))
  
  ## Capture the portion where probability is increasing
  mle = which.max (probLifespanCondR[RCutoffs[ii],])
  if (mle == 1) {
    bottom5Lifespan[ii] = 1
  } else {
    risingProb = probLifespanCondR[RCutoffs[ii], 1:mle]
    ## Approximately where does it cross 0.05?
    if (risingProb[1] > 0.05) { ## The lower CI is below age 0.
      bottom5Lifespan[ii] = 1
    } else {
      f1 = approxfun (1:mle, risingProb)
      f2 = function(x) f1(x) - 0.05
      bottom5Lifespan[ii] = uniroot (f2, interval=c(1,mle))$root
    }
  }

  ## What percentile lifespan does the MLE represent?
  mlePercentile[ii] = cumDistAgeAtDeath[mle]
  
  ## What percentile lifespan does this represent?
  foo = floor(bottom5Lifespan[ii])
  bottom5LifespanPercentile[ii] =
    cumDistAgeAtDeath[foo] +
    (bottom5Lifespan[ii] - foo) *
    (cumDistAgeAtDeath[foo+1] - cumDistAgeAtDeath[foo])

  ## Capture the portion where probability is falling
  if (mle == LCutoff) {
    top5Lifespan[ii] = LCutoff
  } else {
    fallingProb = probLifespanCondR[RCutoffs[ii], mle:LCutoff]
    ## Approximately where does it cross 0.05?
    f1 = approxfun (mle:LCutoff, fallingProb)
    f2 = function(x) f1(x) - 0.05
    if (fallingProb[length(fallingProb)] > 0.05) {
      ## didn't quite get to 0.05
      top5Lifespan[ii] = LCutoff
    } else {
      top5Lifespan[ii] = uniroot (f2, interval=c(mle, LCutoff))$root
    }
  }

  ## What percentile lifespan does this represent?
  foo = floor(top5Lifespan[ii])
  if (foo == LCutoff) {
    top5LifespanPercentile[ii] = cumDistAgeAtDeath[foo]
  } else {
    top5LifespanPercentile[ii] =
      cumDistAgeAtDeath[foo] +
      (top5Lifespan[ii] - foo) *
      (cumDistAgeAtDeath[foo+1] - cumDistAgeAtDeath[foo])
  }
}

d = data.frame (Rpercentiles=RpercentileVector,
                bottom5=bottom5LifespanPercentile,
                top5=top5LifespanPercentile,
                mle=mlePercentile)
if (noEnvVarInFecundity) {
  write.table (d, file="umboniumCILifespanNoEnvVarInFec.dat",
               row.names=FALSE)
} else {
  write.table (d, file="umboniumCILifespan.dat", row.names=FALSE)
}
```
