## Supplementary material for "To prosper, live long: understanding the sources of reproductive skew and extreme reproductive success in structured populations": Associated R code: umboniumLifespanCondLRO.R

```
#########################################################################
##
## This file outputs umboniumLifespanCondLRO.Rdata, which is used as
## input for plotUmboniumLifespanCondLRO2.R, which does the plotting.
##
#########################################################################

rm(list=ls(all=TRUE)); graphics.off();
source ("partitioningUtilities.R")
source ("genericPartitionVarSkewness6.R")
source ("megamatrixFunctions.R")

maxKids = 25
mT = maxKids + 1
maxAge = 19
percentileCutoff = 0.99

## Set noEnvVarInFecundity if you want fecundity in each stage to be
## the average of its values across environments.
noEnvVarInFecundity = FALSE

load ("Umbonium.Rdata")

## Assume all year types are equally likely
numEnv = length(Ulist)
Q = matrix (1/numEnv, numEnv, numEnv)
u0 = rep (1/numEnv, numEnv)

## Assume all individuals are born into stage 1
mz = dim(Ulist[[1]])[1]
c0 = rep (0, mz)
c0[1] = 1

if (noEnvVarInFecundity) {
  Fave = matrix (0, mz, mz)
  for (q in 1:numEnv)
    Fave = Fave + Flist[[q]]
  Fave = Fave/numEnv
  for (q in 1:numEnv)
    Flist[[q]] = Fave
}


## Set up cross-classified birth distribution
bigmz = numEnv*mz
m0 = matrix (outer (c0, as.vector(u0)), bigmz, 1)

out = distLifespanCondR2 (Plist=Ulist, Flist, Q, m0, maxKids,
                         maxAge, percentileCutoff)

distKidsAtDeath = out$distKidsAtDeath
probLifespanCondR = out$probLifespanCondR
sdLifespanCondR = out$sdLifespanCondR
CVLifespanCondR = out$CVLifespanCondR
maxKidsIndex = out$maxKidsIndex
normalSurvProb = out$normalSurvProb


d = data.frame (percentile=cumsum(distKidsAtDeath[1:maxKidsIndex]),
                cv=CVLifespanCondR)

if (noEnvVarInFecundity) {
  write.table (d, file="umboniumCVLifespanCondRNoEnvVarInFec.dat",
               row.names=FALSE)
} else {
  write.table (d, file="umboniumCVLifespanCondR.dat", row.names=FALSE)
}

R50 = min(which(cumsum(distKidsAtDeath) >= 0.5))
R90 = min(which(cumsum(distKidsAtDeath) >= 0.9))
R95 = min(which(cumsum(distKidsAtDeath) >= 0.95))
R99 = min(which(cumsum(distKidsAtDeath) >= 0.99))

probLifespan = -diff(normalSurvProb)

L50 = min(which(cumsum(probLifespan) >= 0.5))
L75 = min(which(cumsum(probLifespan) >= 0.75))
L90 = min(which(cumsum(probLifespan) >= 0.9))
L95 = min(which(cumsum(probLifespan) >= 0.95))
L99 = min(which(cumsum(probLifespan) >= 0.99))

## At what age should we cut off the plots?
##LCutoff = min(which(probLifespanCondR[1,] < 0.001))
LCutoff = L99

if (noEnvVarInFecundity) {
  save (probLifespan, distKidsAtDeath, probLifespanCondR,
        CVLifespanCondR, sdLifespanCondR,
        L50, L75, L90, L95, L99, R50, R90, R95, R99,
        file="umboniumLifespanCondLRONoEnvVarInFec.Rdata")
} else {
  save (probLifespan, distKidsAtDeath, probLifespanCondR,
        CVLifespanCondR, sdLifespanCondR,
        L50, L75, L90, L95, L99, R50, R90, R95, R99,
        file="umboniumLifespanCondLRO.Rdata")
}
```
