## Supplementary material for "To prosper, live long: understanding the sources of reproductive skew and extreme reproductive success in structured populations": Associated R code: Utilities.R

```
#####################################################
# Utilities from the IPMbook code repo 
#####################################################        
domEig=function(A,tol=1e-8,verbose=FALSE) {
	qmax=10*tol; lam=1; x=rep(1,nrow(A));   
	while(qmax>tol) {
		x1=A%*%x;
		qmax=sum(abs(x1-lam*x));  
		lam=sum(x1); 
		x=x1/lam; 
        if(verbose) cat(lam,qmax,"\n"); 
	} 
    x1=A%*%x; qmax=sum(abs(x1-lam*x)); 
    lam=sum(x1); x=x1/lam; 
    if(verbose) cat(lam,qmax,"\n"); 
	return(list(lambda=lam,w=x/sum(x)))
}  	        

matrix.image=function(A, x=NULL, y=NULL, col=rainbow(100,start=0.67,end=0),
             bw=FALSE, do.contour=FALSE, do.legend=TRUE,...) {
 if(do.legend) layout(mat=cbind(matrix(1,5,5),rep(2,5)));
 par(mar=c(6,5,3,2)); 
 if(is.null(x)) x=1:ncol(A);
 if(is.null(y)) y=1:nrow(A); 
 nx=length(x); ny=length(y); 
 x1=c(1.5*x[1]-0.5*x[2],1.5*x[nx]-0.5*x[nx-1]); 
 y1=c(1.5*y[1]-0.5*y[2],1.5*y[ny]-0.5*y[ny-1]); 
 if(bw) col=grey( (200:50)/200 ); 
 image(list(x=x,y=y,z=t(A)),xlim=x1,ylim=rev(y1),col=col,cex.axis=1.5,cex.lab=1.5,bty="u",...);
 abline(v=range(x1)); abline(h=min(y1)-mean(diff(y1))); abline(h=max(y1)+mean(diff(y1))); 
 if(do.contour) contour(x,y,t(A),nlevels=5,labcex=1.2,add=TRUE);   
 
 if(do.legend) {
    l.y=seq(min(A),max(A),length=100);  
    par(mar=c(6,2,3,1))
    image(list(x=1:2,y=l.y,z=rbind(l.y,l.y)),col=col,bty="o",xaxt="n",yaxt="n"); 
    axis(side=2,cex.axis=1.5,at=pretty(seq(min(A),max(A),length=10))); 
 } 
}

#x=1:80; 
#A=outer(x,x,FUN=function(x,y) 0.737*exp(-0.01*(x-y)^2)); 
#px=seq(0,4,length=length(x)); 
#matrix.image(A,px,px,xlab="Size t",ylab="Size t+1",do.contour=TRUE,do.legend=TRUE); 
```
